# A ferredoxin bridge connects the two arms of plant mitochondrial complex I

**DOI:** 10.1101/2020.11.23.393975

**Authors:** Niklas Klusch, Jennifer Senkler, Özkan Yildiz, Werner Kühlbrandt, Hans-Peter Braun

## Abstract

Mitochondrial complex I is the main site for electron transfer to the respiratory chain and generates much of the proton gradient across the inner mitochondrial membrane. It is composed of two arms, which form a conserved L-shape. We report the structures of the intact, 47-subunit mitochondrial complex I from *Arabidopsis thaliana* and from the green alga *Polytomella* sp. at 3.2 and 3.3 Å resolution. In both, a heterotrimeric γ-carbonic anhydrase domain is attached to the membrane arm on the matrix side. Two states are resolved in *A. thaliana* complex I, with different angles between the two arms and different conformations of the ND1 loop near the quinol binding site. The angle appears to depend on a bridge domain, which links the peripheral arm to the membrane arm and includes an unusual ferredoxin. We suggest that the bridge domain regulates complex I activity.

**One sentence summary:** The activity of complex I depends on the angel between its two arms, which, in plants, is adjusted by a protein bridge that includes an unusual ferredoxin.

## 1. INTRODUCTION

Complex I is the largest enzyme complex of the mitochondrial electron-transfer chain. It catalyzes electron transfer from NADH onto ubiquinone, which is coupled to proton translocation across the inner mitochondrial membrane (Agip et al., 2019; Parey et al., 2020; Sazanov, 2015). Bacterial and mitochondrial complex I consist of two parts: the membrane arm and the peripheral arm. The membrane arm is integral to the mitochondrial or bacterial inner membrane, while the peripheral arm protrudes into the bacterial cytoplasm or the mitochondrial matrix. Together, the two arms form an L shape. Each arm has two functional domains: in the peripheral arm these are the NADH-oxidation (N) and the ubiquinone reduction (Q) domains, in the membrane arm they are the proximal (relative to the peripheral arm) and distal proton translocating domains (P_P_ and P_D_ domains).

The first high-resolution structures of complex I were of bacterial origin (Baradaran et al., 2013; Berrisford et al., 2016). *E. coli* complex I has a mass of about 500 kDa and is composed of 14 protein subunits, 7 of which reside in the membrane and 7 in the peripheral arm. This set of conserved subunits forms the core of complex I. Electrons are transferred from NADH to ubiquinone via one flavin mononucleotide (FMN) and 8 or 9 FeS clusters in the peripheral arm. The membrane arm has four potential proton translocation pathways. The molecular mechanisms that couple electron transfer to proton translocation are unknown but thought to involve long-range conformational changes between and within the two complex I arms.

Mitochondrial complex I is significantly larger. Apart from the 14 core subunits, it contains around 30 accessory subunits. The first higher-resolution structures of mitochondrial complex I were from the aerobic yeast *Yarrowia lipolytica* (Zickermann et al., 2015) and mammalian mitochondria (Fiedorczuk et al., 2016; Zhu et al., 2016). Recently, more detailed cryoEM structures were reported for fungal (Grba & Hirst, 2020; Parey et al., 2018) and mammalian complex I (Agip et al., 2018; Kampjut & Sazanov, 2020). Mammalian complex I has 45 subunits and a mass of about 970 kDa. The accessory subunits surround the core subunits and are thought to stabilize the complex. Some of the accessory subunits add new functions to complex I. For instance, two copies of a mitochondrial acyl carrier protein (ACP; also called the SDAP subunit) are integral parts of the mammalian and yeast complex I. Furthermore, a nucleoside kinase is attached to complex I in mammals and a sulfur transferase to that of *Yarrowia* (D’Imprima et al., 2016).

Plants have two different forms of complex I, one each for chloroplasts and mitochondria. The chloroplast complex resembles that of cyanobacteria, which transfers electrons from ferredoxin to plastoquinone. The high-resolution structure of cyanobacterial complex I has recently been resolved by cryoEM (Laughlin et al., 2019; Schuller et al., 2019; Zhang et al., 2020). The structure of plant mitochondrial complex I is less well characterized. Low-resolution single-particle EM revealed a second matrix-exposed domain, which is attached to the membrane arm at a central position (Dudkina et al., 2005). Plant complex I includes additional subunits not present in complex I from *Yarrowia* and mammals (Heazlewood et al., 2003), most notably proteins resembling γ-type carbonic anhydrases (γCAs) (Parisi et al., 2004; Perales et al., 2004). The γCAs form a heterotrimer and were shown to constitute the extra matrix-exposed domain (Fromm et al., 2016; Sunderhaus et al., 2006). It has been proposed that the γCA domain is involved in the transfer of mitochondrial CO_2_ to the chloroplasts for carbon fixation (Braun & Zabaleta, 2007). First insights into the structure of this domain come from single-particle cryoEM of a complex I assembly intermediate from mung bean, which includes 30 of its >45 subunits (Maldonado et al., 2020).

Here we report the high-resolution cryoEM structures of complete complex I from the model plant *Arabidopsis thaliana* in the open and closed state, and from the unicellular heterotrophic green alga *Polytomella* sp. in the closed state. We present new structural and functional insights into plant-specific features of mitochondrial complex I, most notably a protein bridge, which links the peripheral arm to the membrane arm. The bridge appears to adjust the angle between the two complex I arms and may be involved in regulating complex I activity. A recent cryoEM study (Soufari et al., 2020) provides insights into the structure of complex I from cabbage but the map is of lower resolution, shows only one state and the complex is incomplete. In particular, it lacks the bridge domain.

## 2. RESULTS and DISCUSSION

### 2.1 Structure of the intact *Arabidopsis* and *Polytomella* complex I

Intact mitochondrial complex I from *Arabidopsis* was purified as described (Supp. Fig. 1, (Klodmann et al., 2010); *Polytomella* sp. complex I was isolated by a similar protocol (Supp. Fig. 2). Single-particle cryoEM yielded a 3.4 Å map of *Arabidopsis* complex I and a 3.5 Å map for the *Polytomella* complex. Multibody refinement of the peripheral arm, the membrane arm and the P_P_ and P_D_ domains of the membrane arm improved the resolution of *Arabidopsis* complex I up to 3.2 Å and that of *Polytomella* complex I to 3.3 Å (Supp. Figures 3-6). Both have the typical L shape. The γCA domain, which is characteristic for plant mitochondrial complex I, shows up prominently on the matrix side in both maps. The membrane arm is connected by a protein bridge near the γCA domain to the Q domain of the peripheral arm.

#### 2.1.1 Subunit composition and complete model of *Arabidopsis* mitochondrial complex I

*Arabidopsis* complex I consists of 47 subunits (Figure 1, Video 1), including 14 core subunits and 33 accessory subunits (Table 1). We use the subunit designations for bovine complex I (Walker et al., 1992) wherever possible (see Supp. Table 1 for nomenclature). Of the 33 accessory subunits, 24 are conserved and found in both mammalian and plant complex I (Senkler et al., 2017a). Additionally, our structure indicates two copies of the acyl carrier protein (SDAP1 and SDAP2), which were assumed to be absent in plant complex I (Meyer et al., 2007), raising the number of conserved accessory subunits to 26. The remaining 7 accessory subunits seem to be plant-specific. These are three members of the γCA/CAL family; the so-called P1 and P2 proteins (Meyer, 2012); a small unknown hydrophobic subunit on the side of the membrane arm (Supp. Fig. 7); and a ferredoxin, which we refer to as C1-FDX (complex I ferredoxin).

**Figure 1.**
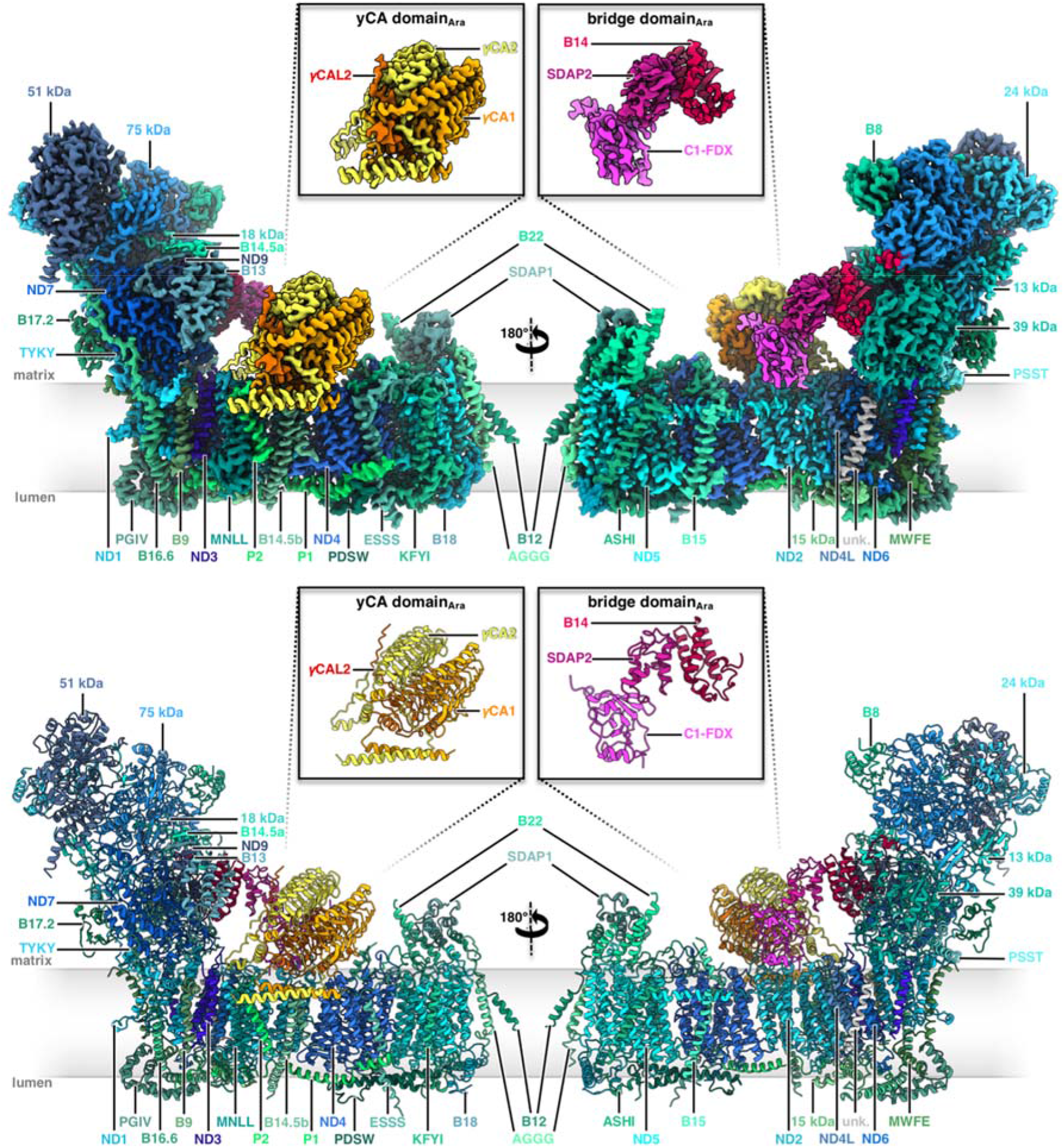
Structure of mitochondrial complex I from *Arabidopsis*. Above: Cryo-EM density; below: atomic model. The 14 core subunits conserved in complex I in bacteria and eukaryotes are drawn in shades of blue; the accessory subunits in shades of green; the three subunits of the carbonic anhydrase domain are yellow, light orange and orange; subunits of the bridge domain are pink, purple and red. Subunit nomenclature as for bovine complex I (Walker et al., 1992). Unknown (unk) refers to one subunit in the membrane arm that was not assigned due to limited amino acid sequence information. Figure insets: y-carbonic anhydrase (γCA) and bridge domains. For EM statistics see Supp. tables 2, 4 and 6.

**Table 1.**
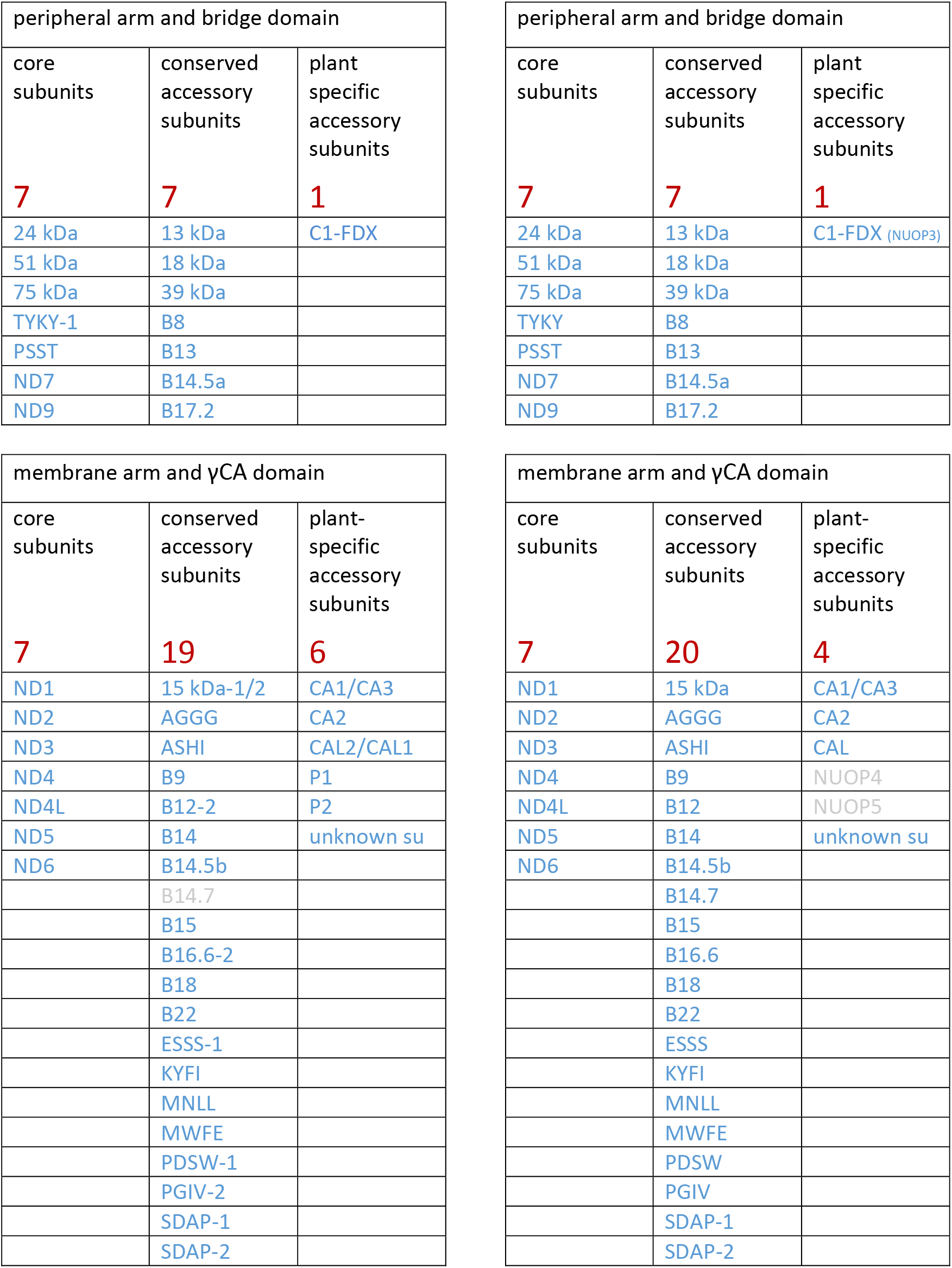
Complex I subunits of *Arabidopsis* and *Polytomella* in the structures shown in Figures 1 and 2. Red, total number of subunits in the subcomplexes. Grey, subunits detected by MS not identified in the cryoEM maps. Extensions - 1/-2 indicate isoforms for some *Arabidopsis* complex I subunits. The dominant isoform was fitted to the map. For protein accession numbers, see Sup. Table 1.

| peripheral arm and bridge domain |  |  |
| --- | --- | --- |
| core subunits | conserved accessory subunits | plant specific accessory subunits |
| **7** | **7** | **1** |
| 24 kDa | 13 kDa | C1-FDX |
| 51 kDa | 18 kDa |  |
| 75 kDa | 39 kDa |  |
| TYKY-1 | B8 |  |
| PSST | B13 |  |
| ND7 | B14.5a |  |
| ND9 | B17.2 |  |

| membrane arm and γCA domain |  |  |
| --- | --- | --- |
| core subunits | conserved accessory subunits | plant-specific accessory subunits |
| **7** | **19** | **6** |
| ND1 | 15 kDa-1/2 | CA1/CA3 |
| ND2 | AGGG | CA2 |
| ND3 | ASHI | CAL2/CAL1 |
| ND4 | B9 | P1 |
| ND4L | B12-2 | P2 |
| ND5 | B14 | unknown su |
| ND6 | B14.5b |  |
|  | B14.7 |  |
|  | B15 |  |
|  | B16.6-2 |  |
|  | B18 |  |
|  | B22 |  |
|  | ESSS-1 |  |
|  | KYFI |  |
|  | MNLL |  |
|  | MWFE |  |
|  | PDSW-1 |  |
|  | PGIV-2 |  |
|  | SDAP-1 |  |
|  | SDAP-2 |  |

Complex I subunits of *Polytomella*
| peripheral arm and bridge domain |  |  |
| --- | --- | --- |
| core subunits | conserved accessory subunits | plant specific accessory subunits |
| **7** | **7** | **1** |
| 24 kDa | 13 kDa | C1-FDX (NUOP3) |
| 51 kDa | 18 kDa |  |
| 75 kDa | 39 kDa |  |
| TYKY | B8 |  |
| PSST | B13 |  |
| ND7 | B14.5a |  |
| ND9 | B17.2 |  |

**Table 1** Complex I subunits of *Arabidopsis* and *Polytomella* in the structures shown in **Figures 1 and 2**.
| membrane arm and γCA domain |  |  |
| --- | --- | --- |
| core subunits | conserved accessory subunits | plant-specific accessory subunits |
| **7** | **20** | **4** |
| ND1 | 15 kDa | CA1/CA3 |
| ND2 | AGGG | CA2 |
| ND3 | ASHI | CAL |
| ND4 | B9 | NUOP4 |
| ND4L | B12 | NUOP5 |
| ND5 | B14 | unknown su |
| ND6 | B14.5b |  |
|  | B14.7 |  |
|  | B15 |  |
|  | B16.6 |  |
|  | B18 |  |
|  | B22 |  |
|  | ESSS |  |
|  | KYFI |  |
|  | MNLL |  |
|  | MWFE |  |
|  | PDSW |  |
|  | PGIV |  |
|  | SDAP-1 |  |
|  | SDAP-2 |  |

Four complex I subunits that were previously identified in *Arabidopsis* by mass spectrometry (Supp. Fig. 8) are absent in our structure: (i) a L-galactono-1,4-lactone dehydrogenase; (ii) a TIM—like protein *(Arabidopsis* accessions At1g18320 and At3g10110); (iii) subunit B14.7; (iv) subunit SGDH. Of these, GLDH only binds to assembly intermediates of complex I (Schertl et al., 2012) and is not expected in the holo complex, as recently confirmed by cryoEM (Soufari et al., 2020). The TIM-like protein may not be a true complex I subunit but might have been co-purified in earlier preparations. B14.7 is conserved in mammals, *Yarrowia* and *Polytomella*, where its main role is thought to be in supercomplex formation (Kampjut & Sazanov, 2020; Letts et al., 2016). As a large fraction of *Arabidopsis* complex I forms a supercomplex with complex III_2_ (Eubel et al., 2003), the B14.7 subunit may have dissociated during complex I preparation. The SGDH subunit, like B14.7, is a conserved accessory subunit of mitochondrial complex I. Its location in mammalian complex I corresponds to that of the plant-specific P1 subunit in *Arabidopsis*. Since mammalian SGDH and *Arabidopsis* P1 share some sequence similarity (Supp. Fig. 9), we conclude that P1 is a plant equivalent of SGDH. In total, *Arabidopsis* complex I consists of 47 subunits plus subunit B14.7 (Table 1), which makes it the largest complex I assembly characterized to date.

Apart from the 47 protein subunits, our structure of *Arabidopsis* complex I indicates 15 cofactors, including 11 in the peripheral arm (8 FeS clusters, 1 FMN, 1 Zn^2+^ and 1 NADPH), and four in the membrane arm (2 phosphopantetheine groups, 1 Zn^2+^ and 1 Fe ion), plus eight lipids and one bound LMNG detergent molecule (Supp. Fig. 10) in the membrane arm.

*Arabidopsis* complex I was prepared without added substrates, and therefore the NADH oxidation site is empty, as expected. However, the complex binds ubiquinone at site 2 in the Q tunnel between the lipid bilayer and the Q reduction site, as previously described for *Yarrowia* complex I (Parey et al., 2019).

#### 2.1.2 Subunit composition and model of *Polytomella* mitochondrial complex I

The subunit composition of complex I from *Polytomella* sp. has been less well defined. We analyzed purified *Polytomella* complex I by 2D SDS/SDS PAGE and mass spectrometry (Supp. Figures 11-14). In total, we identified peptides of 40 subunits that resembled known complex I components from other organisms, in particular *Chlamydomonas*. Shotgun MS analyses revealed another four polypeptides, raising the total number of subunits to 44 (Supp. Fig. 11; see also the GelMap at www.gelmap.de/2062 [password: Poly-C1]). Since the mass spectrometry data did not cover the complete amino acid sequences especially of the hydrophobic subunits, additional peptide sequence information was derived from genomes. The genome sequence of *Chlamydomonas* is known (Merchant et al., 2007). A partial genome sequence of *Polytomella* sp. has been determined recently (Murphy et al., 2019), but remains to be fully annotated. We used exon-intron prediction programs to identify open readings frames that encode the complete polypeptide sequences of complex I subunits in *Polytomella*.

We were able to assign 43 subunits in the 3.3 Å cryo-EM map of *Polytomella* complex I (Figure 2, Video 2), including the complete set of the 14 core subunits and 29 of the accessory subunits (Table 1). In addition, three accessory subunits not identified by MS were assigned on the basis of their map density (Supp. Figure 15), bringing the total to 46. 31 of the 32 accessory subunits are conserved between *Polytomella* and *Arabidopsis*, indicating a remarkably similar subunit composition of mitochondrial complex I in plants and algae. The plant-specific P1 and P2 subunits are absent in *Polytomella*. As in *Arabidopsis, Yarrowia* and mammals, the acyl carrier proteins (SDAP1 and 2) bind in two distinct locations. The *Polytomella* ferredoxin-like subunit C1-FDX is a homolog of the NUOP3 subunit previously identified in *Chlamydomonas* complex I (Cardol et al., 2008; Cardol et al., 2005; Cardol et al., 2004). The small unknown hydrophobic subunit at the side of the *Arabidopsis* membrane arm is conserved in *Polytomella*, but its sequence was not determined. In contrast to *Arabidopsis*, the B14.7 subunit is present in the *Polytomella* complex I structure. As in mammals and *Yarrowia*, it is found at the position where complex I interacts with the complex III dimer within the I+III_2_ supercomplex. Subunits NUOP4 and NUOP5, which were detected by mass spectrometry in *Polytomella* and *Chlamydomonas* complex I Table 1, (Cardol et al., 2005; Cardol et al., 2004), were not found in the *Polytomella* cryoEM map. A few regions of the *Polytomella* map were left unassigned (Supp. Fig. 16). These map regions may belong to unknown parts of the identified subunits, or to new subunits that remain to be identified. The overall structures of *Arabidopsis* and *Polytomella* complexes I are similar, including the trimeric γCA domain and the bridge connecting the Q domain of the peripheral arm to the membrane arm near the γCA domain. *Polytomella* complex I appears to be more compact, perhaps because some of its subunits are longer than in *Arabidopsis* and thus can form stronger contacts.

**Figure 2.**
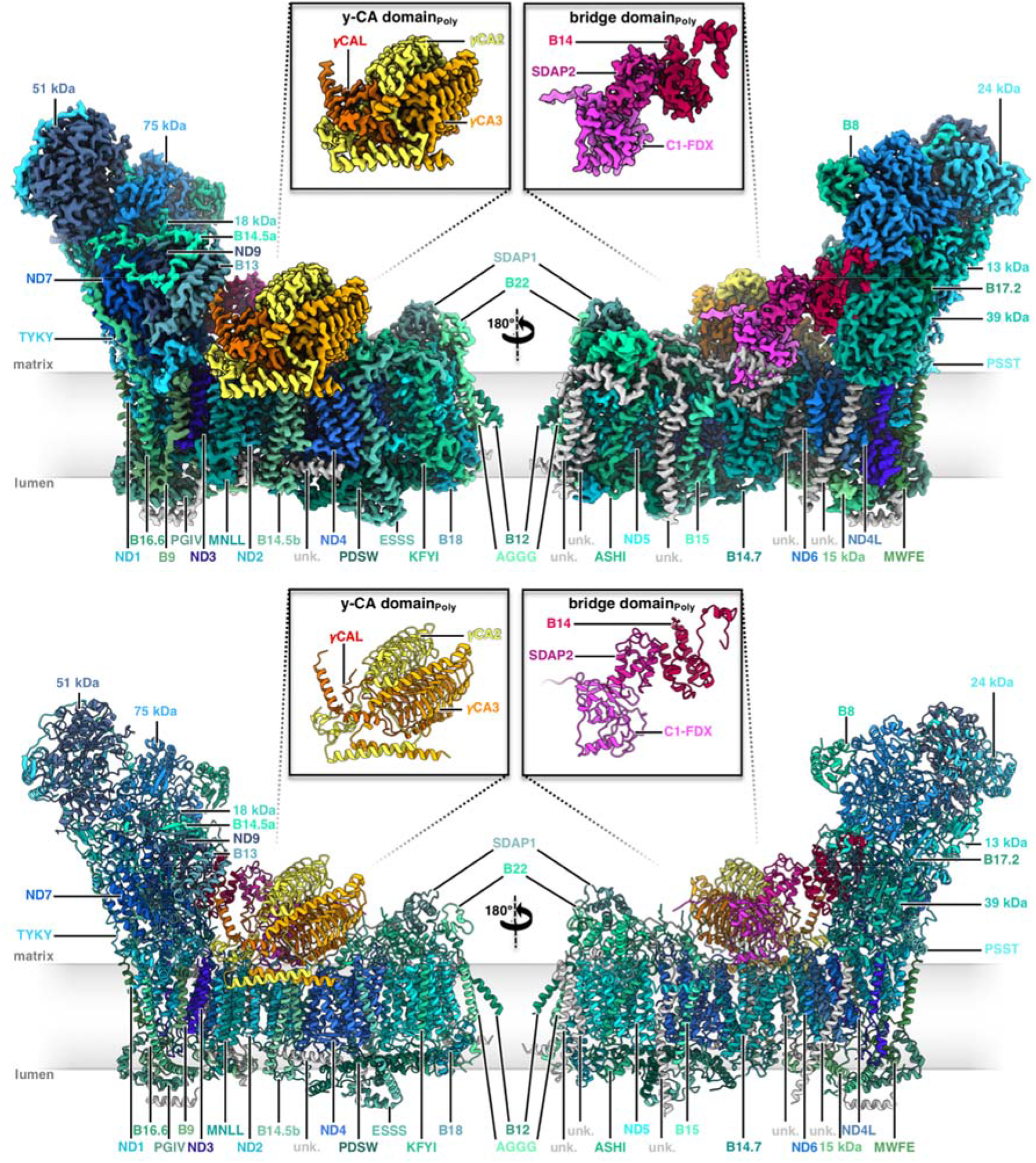
Structure of mitochondrial complex I from *Polytomella* sp. Above: Cryo-EM density; below: atomic model. Color scheme for the 14 conserved core subunits, accessory subunits and the subunits of the carbonic anhydrase and bridge domains as in Figure 1. Insets: y-carbonic anhydrase (γCA) and bridge domains. Unknown (unk) refers to six unassigned densities in the membrane arm with limited amino acid sequence information. For EM statistics see Supp. tables 3, 5 and 7.

Apart from its 46 protein subunits, our structure of *Polytomella* complex I indicates 13 bound co-factors (8 FeS clusters, 1 FMN, 1 Zn^2+^ and 1 NADPH in the peripheral arm, two phosphopantetheine groups in the two acyl carrier subunits of the membrane arm), plus 12 lipid molecules (Supp. Fig. 17).

### 2.2 The γ-carbonic anhydrase heterotrimer in *Arabidopsis* and *Polytomella*

In plant mitochondria, a heterotrimeric γCA domain is attached to the membrane arm of complex I (Braun, 2020; Fromm et al., 2016; Sunderhaus et al., 2006). The γCA domain is absent in complex I from mammals, fungi, bacteria and chloroplasts but seems to occur in several groups of protists (Gawryluk & Gray, 2010). This suggests that the γCA proteins are of ancient origin and most likely formed part of complex I in the earliest ancestors of the eukaryotic clade. The *Arabidopsis* genome encodes five different γCA subunits, all of them associated with mitochondrial complex I (Klodmann et al., 2010). Of these, three have the conserved amino acid residues that form the active site, as in the prototypic γ-carbonic anhydrase of the archaeon *Methanosarcina thermophila* (Kisker et al., 1996). They are referred to as γCA1, γCA2 and γCA3 (Parisi et al., 2004). Two others are known as gamma carbonic anhydrase-like proteins γCAL1 and γCAL2 (Perales et al., 2004). The complex I-integral γCA/CAL proteins of *Arabidopsis* assemble into heterotrimers of two γCA proteins and one γCAL (Braun, 2020; Fromm et al., 2016), but the precise composition of the trimers was unknown. Our complex I structure of *Arabidopsis* includes γCA2, γCA1 and γCAL2, as indicated by evaluation of amino acid positions that differ between the γCA and γCAL proteins (Figure 3a,c; Supp. Fig. 18). Note that the high-resolution map of the *A. thaliana* complex suggests a mixed occupancy of the γCA/CAL heterotrimer, because side chains of alternative but less abundant γCA subunits can be fitted at some positions. This is in line with the finding that γCA subunits can substitute for each other in *Arabidopsis* knockout lines (Fromm et al., 2016).

**Figure 3.**
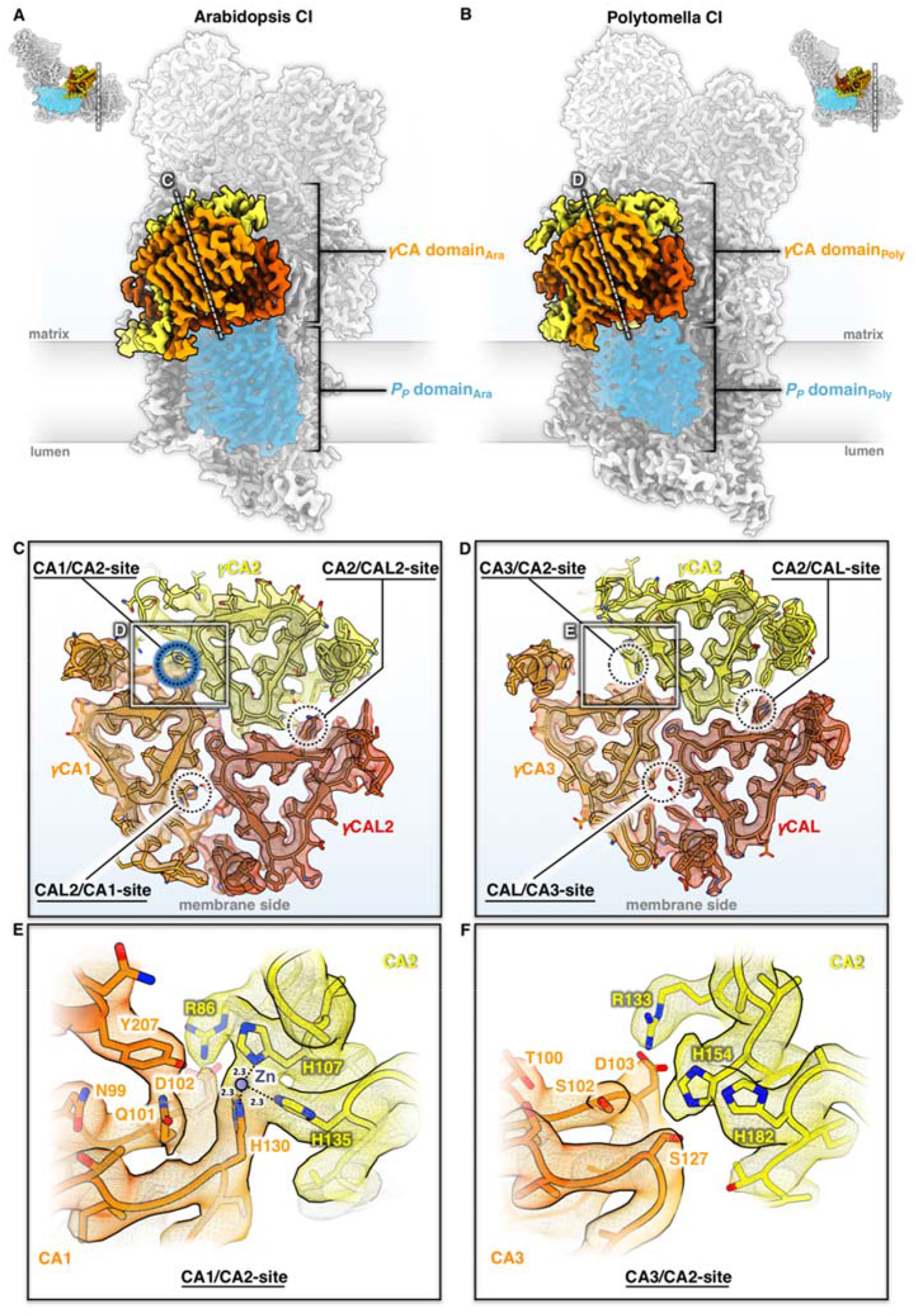
The γCA domain of *Arabidopsis* and *Polytomella* mitochondrial complex I. **A, B**: Position of the γCA domain on the membrane arm. Complex I (CI) is seen from the tip of the membrane arm and cut at the plane indicated by a dashed white line in the small insets. Subunit color scheme as in Figures 1 and 2. The P_P_ domain of the membrane arm is shaded blue. **C, D**: Cross-sections of the γCA domains of *Arabidopsis* (C) and *Polytomella* (D) at the level of the catalytic sites at the γCA/CAL subunit interfaces, as indicated by dashed circles. In *Arabidopsis*, one of the three possible catalytic sites coordinates a metal (presumably zinc) ion (blue) and is therefore potentially active, whereas none of the three sites of the *Polytomella* γCA domain are occupied, and therefore they are inactive. **E, F**: Details of catalytic site regions in *Arabidopsis* (E) and *Polytomella* (F) (white boxes in C and D). Amino acids are indicated by the one-letter code. For further details, see Sup. Figures 18-21.

The fold of the γCA/CAL proteins is highly conserved, consisting of a central left-handed triangular β-helix, which is laterally flanked by a C-terminal α-helix (Supp. Fig. 19a,b). In contrast to *M. thermophila*, γCA1 and γCA2 each have a long amphiphilic alpha helix at the N-terminus, which is absent in γCAL2. The two amphiphilic helices of γCA1 and γCA2 form a coiled coil parallel to the membrane arm on the matrix side of the inner mitochondrial membrane. A gap between the coiled coil and the membrane arm is filled with a distinct set of lipids, which might help to attach the γCA domain to complex I (Supp. Fig. 19c). The γCA domain interacts with the ND2 and B14.5b subunits, the plant-specific complex I protein P2 and C1-FDX (see below). Interaction of the γCA trimer and the membrane arm is restricted to the P_P_ module. The γCA/CAL subunits are part of early assembly intermediates; in their absence, assembly of plant mitochondrial complex I is arrested (Ligas et al., 2019).

The archaeal γCA of *M. thermophila* is a homotrimer with three active sites at the subunit interfaces, each with a zinc ion coordinated by three histidines, two of which belong to one and the third to another, neighboring subunit (Kisker et al., 1996). In *Arabidopsis*, only the γCA1/γCA2 interface has a complete set of active-site histidines (γCA1_HisH130, γCA2_HisH107 and γCA2_HisH135). Together these three side chains coordinate a metal (presumably Zn) ion in a non-peptide density (Figure 3c,e). The nearby conserved sidechains γCA1_N99, γCA1_Q101 γCA1_D102, γCA1_Y207 and γCA2_R86 are crucial for stability and the catalytic mechanism (Ferry, 2010; Iverson et al., 2000), suggesting that the γCA1/γCA2 site is active. The two other sites in the heterotrimer lack some of the conserved zinc-binding residues, do not show a non-peptide density, and are therefore presumably inactive.

In *Polytomella*, three γCA proteins and one γCAL protein were identified by MS and evaluation of the partial genome sequence. Based on sequence similarity to *Arabidopsis*, we refer to the *Polytomella* γCA subunits as γCA1, γCA2 and γCA3. Of these, γCA2 and γCA3 are present in *Polytomella* complex I, in addition to one copy of γCAL (Figure 3b and Supp. Fig. 20). As in *Arabidopsis*, the structure of *Polytomella* complex I indicates a degree of mixed occupancy of γCA subunits, because alternative, less abundant side chains can be fitted at some positions. It is therefore likely that a small fraction of *Polytomella* complex I contains γCA1, γCA2 and γCAL. The topological arrangement of the γCA/CAL subunits and the anchoring of the γCA domain by the coiled-coil N-terminal amphipathic helices is very similar to *Arabidopsis* (Supp. Fig. 21). Surprisingly, none of the three potential active sites at the subunit interfaces has the complete set of three zinc-coordinating residues (Figure 3d,f). At the γCA2/γCA3 interface, the third histidine is substituted by γCA3_S127 and nearby residues that participate in catalysis have been replaced. Since none of the three potential catalytic sites show any density for a bound metal ion, we conclude that the *Polytomella* γCA domain is inactive.

In photosynthetically active organisms, carbonic anhydrases play a role in carbon assimilation and carbon concentration mechanisms or pH stabilization. It has been suggested that the carbonic anhydrase of complex I is required for the transfer of carbon dioxide from mitochondria to the chloroplasts for carbon fixation in the Calvin-Benson cycle (Braun & Zabaleta, 2007). Since *Polytomella*, unlike its close relative *C. reinhardtii*, does not photosynthesize, it might not need this activity. It recently has been found that cyanobacterial complex I is involved in carbon transport and concentration (Schuller et al., 2020). However, the cyanobacterial carbonic anhydrase subunits belong to a different enzyme class, and they are attached to the P_D_ domain at the tip of the membrane arm. Furthermore, one of the proton transfer pathways in the cyanobacterial P_D_ domain appears to have adapted to CO_2_ transfer. In contrast, the γCA domain of *Arabidopsis* mitochondrial complex I is attached to the P_P_ domain and its active site is not close to a proton transfer path. Our structure thus does not suggest that plant mitochondrial complex I is directly involved in CO_2_ or bicarbonate transport across the inner mitochondrial membrane. However, bicarbonate formed at the γCA domain might be exported from the mitochondrial matrix by transporters unrelated to complex I.

### 2.3 A ferredoxin bridge between the ubiquinone-reduction and the γ-carbonic anhydrase domain

A striking feature of the *Arabidopsis* and *Polytomella* mitochondrial complex I is a three-subunit protein bridge, which forms a physical link between the Q domain of the peripheral arm and the membrane arm (Figure 4). The bridge consists of (i) subunit B14 in the peripheral arm, (ii) one of the two acyl carrier proteins (SDAP2) with a bound phosphopantetheine and (iii) a ferredoxin-like subunit (here referred to as complex 1 ferredoxin, C1-FRX). C1-FRX is connected to the core ND2 subunit of the membrane arm and the γCAL2 subunit of the γCA domain. The B14 and acyl carrier subunits are conserved in mammalian and *Yarrowia* complex I. The B14 subunit belongs to the eukaryotic LYR protein family (Angerer et al., 2014) that is defined by a [Leu, Tyr, Arg] motif close to the N-terminus. LYR proteins are comparatively small, mitochondria-specific and positively charged components of respiratory chain complexes or act as assembly factors. ISD11, another LYR protein, is part of the iron sulphur cluster (ISC) assembly complex. Mitochondrial acyl carrier proteins are confined to the mitochondrial matrix (Angerer et al., 2017), where they are involved in fatty acid biosynthesis, in particular lipoic acid, which is a prosthetic group of several mitochondrial enzymes, and possibly also longer fatty acids for membrane biogenesis. The two acyl carrier proteins bound to complex I both carry longer fatty acids, which interact with the LYR-like subunits B14 and B22. Finally, mitochondrial acyl carrier proteins are known to form part of the ISC assembly complex, bind to ISD11, and might have a regulatory role in FeS cluster biosynthesis (Lill, 2020).

**Figure 4.**
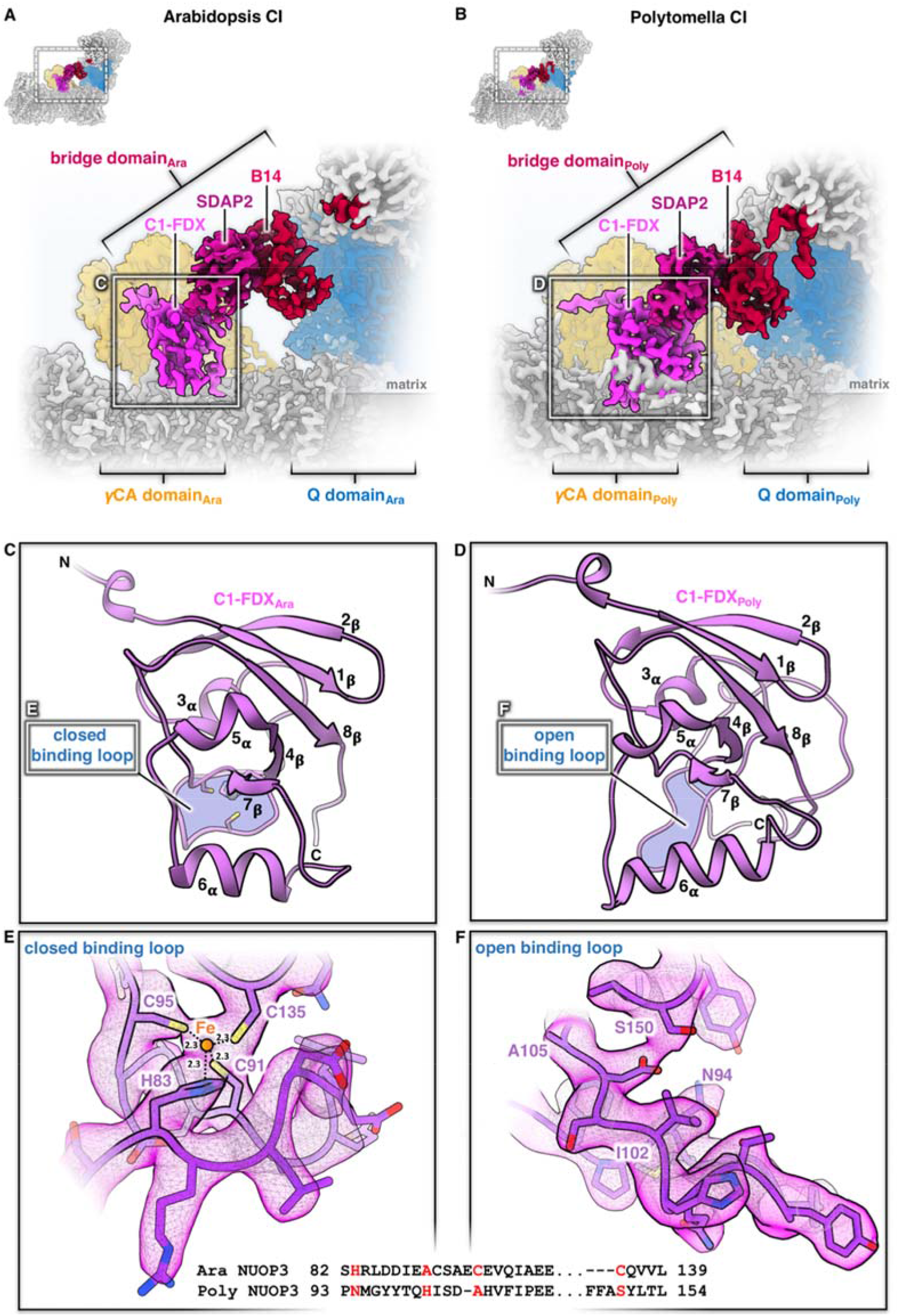
The bridge domain of *Arabidopsis* and *Polytomella* complex I (CI). **A, B:** Attachment of the bridge domain linking the membrane arm near the γCA domain (orange) to the Q domain of the peripheral arm (blue). **C, D:** Structure of the C1-FDX subunit in *Arabidopsis* and *Polytomella*. **E,F:** catalytic sites of C1-FDX in *Arabidopsis* and *Polytomella*. For details, see Sup. Figs. 16 and 22-24.

Blue-native/ SDS PAGE did not identify the C1-FDX subunit in plant complex I, but low levels of a ferredoxin were detected by complexome profiling (Senkler et al., 2017b; Takabayashi et al., 2017). The *Arabidopsis* C1-FDX subunit is homologous to NUOP3, which is part of *Chlamydomonas* complex I (Cardol et al., 2005; Cardol et al., 2004). An *Arabidopsis* mutant lacking the gene encoding C1-FDX has decreased complex I levels (Hansen et al., 2018) but remains to be fully characterized. The copy number of *Arabidopsis* C1-FDX is estimated to be 2600 per single mitochondrion by quantitative mass spectrometry, in excellent agreement with the copy number of average mitochondrial complex I subunits (2500 per single mitochondrion; (Fuchs et al., 2020). In our *Arabidopsis* complex I, prepared by sucrose density gradient centrifugation, C1-FDX clearly is an accessory complex I subunit. NUOP3, the *Polytomella* homolog of C1-FDX, is a structural subunit of the *Polytomella* bridge domain.

The 3D structure of *Arabidopsis* and *Polytomella* C1-FDX closely resembles mammalian and fungal mitochondrial ferredoxin 1 and 2, with its characteristic β-grasp fold (Supp. Fig. 22).

Furthermore, C1-FDX resembles two mitochondrial ferredoxins of *Arabidopsis* known as AtMFDX1 and AtMFDX2 (Takubo et al., 2003). Although sequence identity between C1-FDX and AtMFDX1/AtMFDX2 is low, the structure of C1-FDX and the predicted structures of AtMFDX1 and AtMFDX2 modelled on human mitochondrial ferredoxin are very similar (Supp. Fig. 23). Mitochondrial ferredoxins are involved in the formation of FeS clusters (Cai et al., 2017; Lange et al., 2000). Typically, they have a central 2Fe2S cluster themselves, coordinated by four conserved cysteine residues in the binding loop of the core domain. In *Arabidopsis* C1-FDX, one of these cysteines is substituted by a histidine (Figure 4). Judging from its map density, the ligand bound by the four side chains of C1-FDX_H83, C1-FDX_C91, C1-FDX_C95 and C1-FDX_C135 cannot be a 2Fe2S cluster, but must be a single metal ion, most likely iron. This would distinguish C1-FDX from all other mitochondrial ferredoxins of mammals, fungi and plants. In *Polytomella*, the C1-FDX equivalent NUOP3 lacks all four conserved cysteines. As a result, the core domain loop is locked in a state that cannot not bind a metal ion (Figure 4), and therefore the *Polytomella* ferredoxin is inactive.

What could be the functional role of the bridge domain of plant and algal complex I? Interestingly, homologs of B14, the SDAP protein and C1-FDX form a functional module within the ISC assembly machinery. Together with the cysteine desulfurase NFS1, the scaffold protein ISCU and frataxin, they can perform *de novo* FeS cluster biosynthesis (Boniecki et al., 2017; Cory et al., 2017; Fox et al., 2019; Lill, 2020). We conclude that elements of the ISC assembly machinery are part of complex I in algae and plants. Nevertheless, biosynthesis of complete FeS clusters on intact complex I is unlikely, because this would require the other components of the ISC assembly machinery to bind in positions where they would interfere with the peripheral arm (Supp. Fig. 24). However, it is possible that assembly intermediates of complex I lacking the peripheral arm bind the missing components of the ISC assembly machinery transiently and then catalyse the formation of FeS clusters. Presumably, our structure shows how these subunits interact within the ISC assembly machinery.

### 2.4 Ferredoxin binding to plant mitochondrial complex I sets the angle between its two arms

The angle between the membrane and peripheral arms of mammalian complex I varies. Two states have been described, referred to as open (112° angle between the two arms) and closed (105°) (reviewed in (Parey et al., 2020)). Our analysis of *Arabidopsis* complex I likewise revealed two states that resemble those of the mammalian complex. About one third of the intact *Arabidopsis* particles are in the closed state, with an angle of 106° between the membrane and peripheral arms, whereas two thirds are in the open state, with an angle of 112°. At 3.7 Å resolution, 3D maps calculated for these conformational classes reveal that complex I in the open state lacks the acyl carrier and C1-FDX subunits almost entirely (Figure 5a,b,c). In the closed state, both subunits are present and well-defined. Our data suggest that the binding of the acyl carrier and C1-FDX subunits to *Arabidopsis* complex I sets the angle between its two arms. In the more robust *Polytomella* complex I, a helix of one of its unidentified accessory subunits wraps around the lower part of the ferredoxin, holding it firmly in place and preventing its dissociation (Figure 4b; Sup. Fig. 16). This explains why the *Polytomella* complex is found exclusively in the closed conformation.

**Figure 5.**
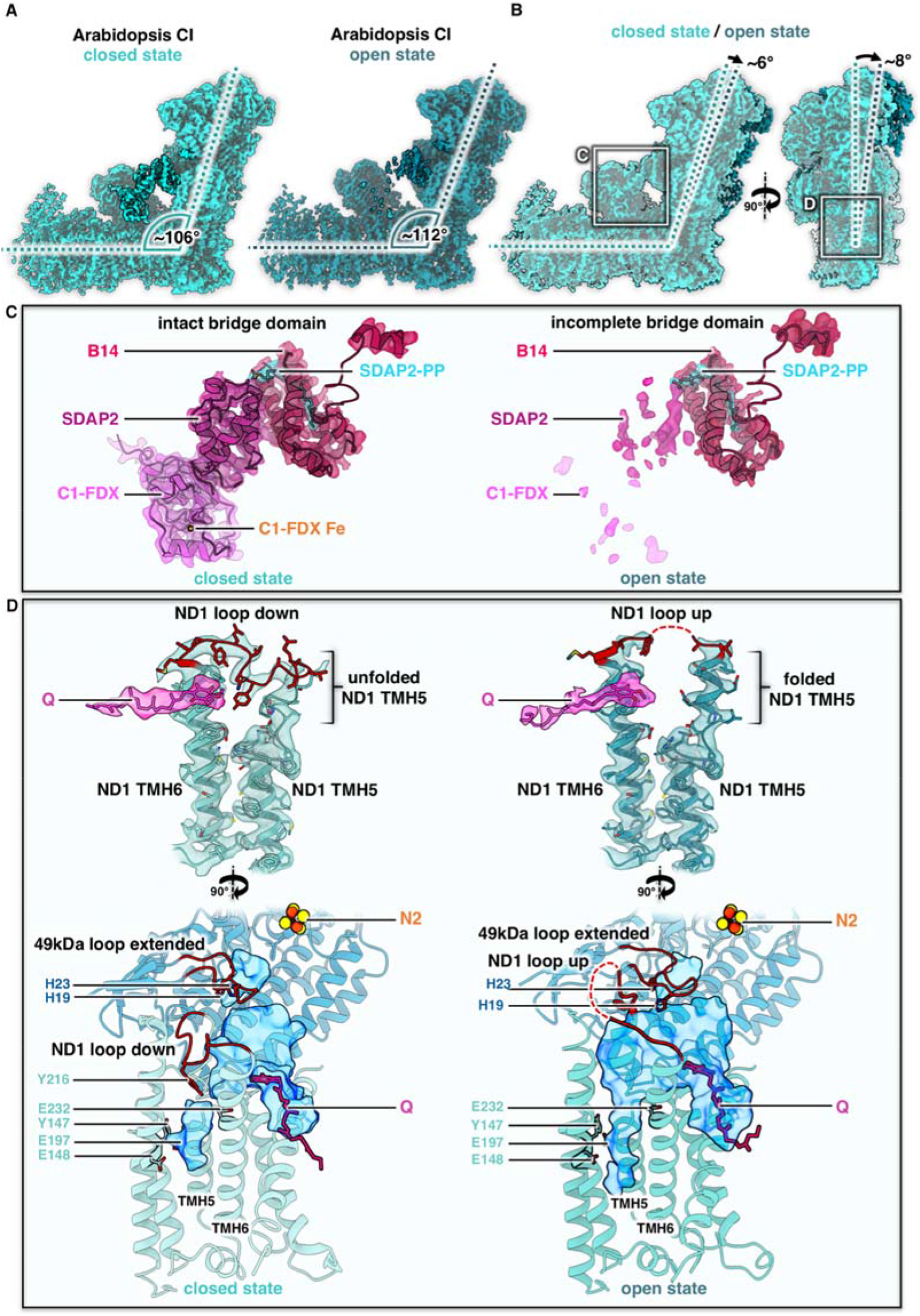
Conformations of *Arabidopsis* complex I (CI). **A:** closed (angle 106°) and open (112°) conformation. **B:** Front and side views of superposed maps shown in A. **C:** Map density and fitted atomic models of the bridge domain in *Arabidopsis*. Subunits color scheme as in Figures 1 and 2. **D**: Conformation of the Q-binding site in the open and closed complex I conformations. Top: orientation of the loop between transmembrane helix (TMH) 5 and TMH6 in ND1. Below: Q-binding channel (blue) with E channel on the left, indicating different conformations of the ND1 and 49 kDa loops with conserved amino acids drawn as sticks. For details, see Supp. Figure 25.

The transition from the open to the closed state in *Arabidopsis* complex I is associated with a conformational switch of the loop linking TM helices 5 and 6 of the core ND1 subunit in the membrane arm. The ND1 loop is in a well-defined “down” conformation in the closed state, while the upper part of TMH 5 near the matrix surface of the membrane arm is unfolded (Figure 5d). In the open state, the unfolded stretch folds into three turns of an alpha helix, extending TMH 5 towards the matrix, and the loop switches from the “down” to an “up” conformation. The same conformational switch of the ND1 loop has been reported for the ovine complex (Supp. Fig. 25b). In its catalytic cycle, complex I has been suggested to alternate between the open and closed state (Kampjut & Sazanov, 2020). Mammalian complex I can assume a number of different open conformations, one of which has been assigned to the deactive state (Kampjut & Sazanov, 2020). The deactive state of the ovine complex is arrested by TMH 4 of the core ND6 subunit, which tilts by ~35° to a new position on the outside of the membrane arm (Supp. Fig. 25e). In *Polytomella* (Supp. Fig. 16, unknown 5) and in both states of the *Arabidopsis* complex (Supp. Fig. 7), this position is occupied by the transmembrane helix of an unknown accessory subunit, and therefore deactivation of the plant complex must proceed by a different mechanism.

The transition between the deactive and active states of mammalian and *Yarrowia* complex I is accompanied by distinct conformational changes of several subunit loops (Agip et al., 2018; Grba & Hirst, 2020; Kampjut & Sazanov, 2020; Parey et al., 2019) near the Q reduction site (site 1) at the end of the Q channel and the so-called E channel, an aqueous cavity branching off the Q channel (Figure 5d). In both states of the *Arabidopsis* complex I, the native quinol substrate binds near the Q channel entrance (site 2) (Figure 5d). Apart from a major change in the ND1 loop (Figure 5d, Supp. Fig. 25a), detailed comparison of the closed and open states of the *Arabidopsis* complex revealed only minor changes in loop conformations. In both states, the 49 kDa loop is extended and blocks access to the Q reduction site (Figure 5d, Supp. Fig. 25c) and the PSST loop with its two Arg residues has the same conformation (Supp. Fig. 25d). In the closed state of the *Arabidopsis* complex, the E channel is blocked by Tyr216 in the ND1 loop in the “down” position (Figure 5d, Supp. Fig. 25a). In the open state of the *Arabidopsis* complex, the ND1 loop is in the “up” position and the E channel is clear. Both the ND1 loop and the 49 kDa loop are in the same respective conformations in the open states of the ovine and *Arabidopsis* complex (Supp. Fig. 25a,c). Notwithstanding their remarkable similarity, clear differences exist between the *Arabidopsis* and ovine complex I in the loop conformations that control access to the Q reduction site and E channel. These differences most likely reflect the occupation of the Q reduction site by quinol substrate or an inhibitor.

## CONCLUSION

The structures of mitochondrial complex I from *Arabidopsis* and *Polytomella* show that the heterotrimeric γCA domain is attached to the membrane arm by the coiled-coil amphipathic helices of the two γCA subunits. Only one of the three potential active sites of the γCA domain binds a metal ion in *Arabidopsis*, whereas none of them do in *Polytomella*. The γCA domain has been suggested to promote transfer of mitochondrial CO_2_ for carbon fixation by the Calvin-Benson cycle to the chloroplasts. The γCA domain is known to be essential for plant mitochondrial complex I assembly, but its connection to the bridge domain suggests an additional role in controlling the mutual orientation of the two complex I arms, and possibly their activity.

Of the three accessory subunits in the bridge domain, the peripheral arm protein B14 and the acyl carrier protein SDAP2 are conserved in mammals and *Yarrowia*. In the plant complex, the bridge it is completed by the special ferredoxin C1-FDX that appears to coordinate an iron ion instead of the usual FeS cluster. In the bridge, C1-FDX fills the gap between the acyl carrier protein and the core subunit ND2 in the membrane arm close to the γCAL subunit of the carbonic anhydrase domain. The three-subunit protein bridge seems to control the angle between the two complex I arms in the open and closed state (Figure 6). We assume that the active form of *Arabidopsis* complex I would be in a closed state, as it corresponds to the active state of mammalian complex I and the closed state of the ovine complex in the catalytic cycle (Kampjut & Sazanov, 2020). It is tempting to speculate that C1-FDX is involved in the regulation of complex I activity, either by dynamic interaction of the ferredoxin with the fully assembled complex, or by its incorporation in a final step of the assembly pathway. We expect that a similar ferredoxin bridge links the acyl carrier subunit to ND2 in mammals, but that this bridge is less stable and therefore breaks or dissociates upon isolation. Regulation by C1-FDX may depend on the oxidation state of the bound metal ion, which might act as a sensor to align complex I activity with the redox state of the mitochondrial matrix. Such a mechanism might be of special significance for photosynthetic organisms.

**Figure 6:**
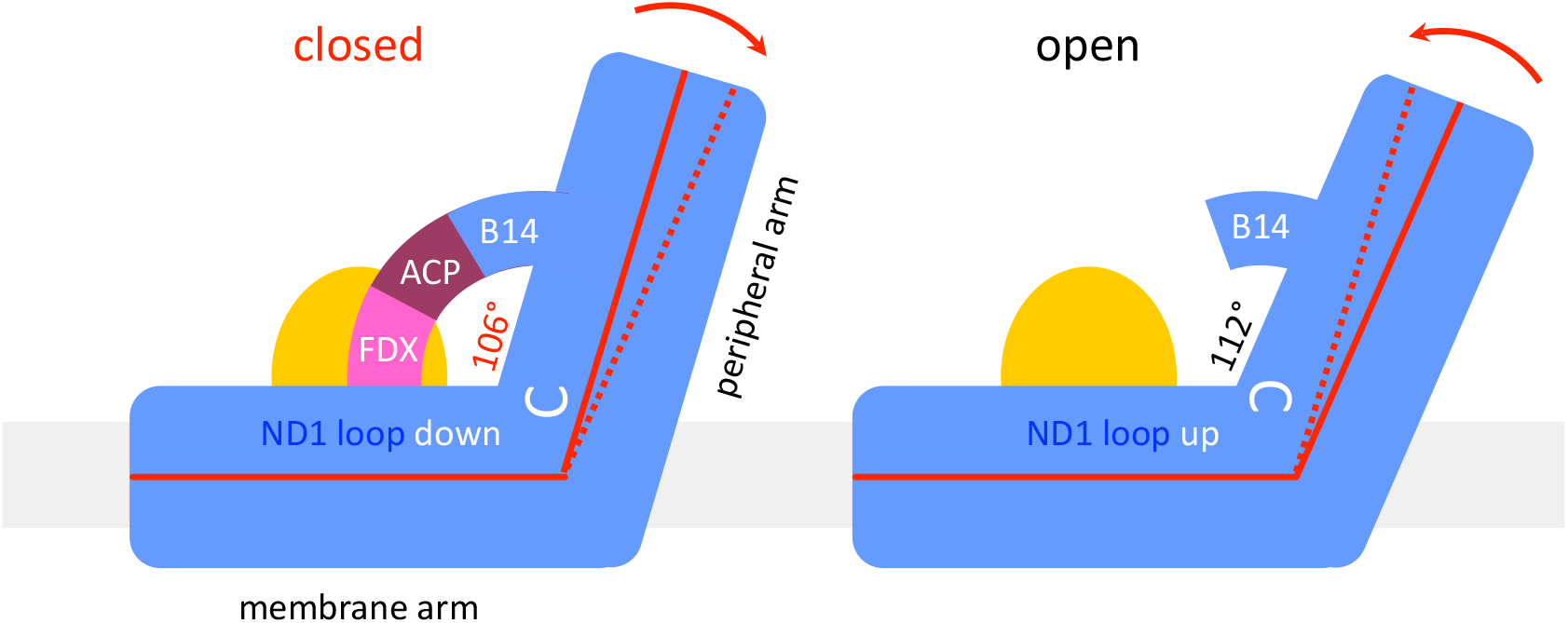
The ferredoxin bridge of *Arabidopsis* complex I. In the closed state, subunit B14 of the peripheral arm forms a bridge with the acyl carrier protein SDAP2 (purple) and the complex I ferredoxin C1-FDX (pink). Ferredoxin joins the bridge to the membrane arm near the gamma carbonic anhydrase domain (orange). The ND1 loop linking TMH5 and 6 of the ND1 subunit in the membrane arm (white) is down. The bridge sets the angle between the membrane and peripheral arms to 106°. In the open state, the ND1 loop flips to the “up” position, the acyl carrier protein and ferredoxin dissociate, and the angle relaxes to 112°. Membrane, grey. The position of the gamma carbonic anhydrase domain is taken by a 42 kDa nucleoside kinase in mammalian complex I.

## Author contributions

HPB and WK initiated the project. NK purified complex I from *Polytomella*, NK and JS purified complex I from *Arabidopsis*. NK collected cryoEM data, performed image processing and produced the figures. NK and ÖY built and analyzed the atomic models. JS carried out proteome analyses. All authors evaluated data. HPB and WK wrote the manuscript, with contributions from NK and JS.

## Acknowledgements

We thank Janet Vonck and Volker Zickermann for critical comments on the manuscript. The work was funded by the Max Planck Society (WK, ÖY, NK) and by the Deutsche Forschungsgemeinschaft (grant BR 1829/10-2 – HPB, JS; SFB 807 – WK, NK).

## Materials and Methods

### Plant material

*Arabidopsis thaliana* was cultivated as described (Farhat et al., 2019). In short, *Arabidopsis* plants (ecotype Columbia 0) were grown under sterile conditions in a growth chamber (16h light / 8h dark, 22°C) for one week. Leaves were cut into small pieces and placed on B5 medium (3.16 g/l B5-medium, 3% sucrose [w/v], 0.75% agar [w/v], 0.5 mg/l 2,4-D, 0.05 mg/l kinetin, pH5.7) to induce callus formation. After three weeks, callus tissue was transferred into liquid B5 medium, which was refreshed once per week. The cell suspension culture was maintained at 22°C on a shaker in the dark. *Polytomella* sp. cells (198.80, E.G. Pringsheim) were ordered from the SAG Culture Collection of Algae (Göttingen University, Germany) and cultivated in acetate medium (0.2% [w/v] sodium acetate, 0.1% [w/v] tryptone peptone, 0.1% [w/v] yeast extract) at 20°C in the dark. The medium was changed twice per week.

### *Arabidopsis thaliana* mitochondria isolation

Mitochondria were isolated from ~150 g of cells from *A. thaliana* cell suspension cultures. Cells were harvested with a sieve. All following steps were performed at 4°C or on ice. Cells were disrupted in grinding buffer (450 mM sucrose, 15 mM MOPS, 1.5 mM EGTA, 0.6% [w/v] PVP40, 0.2% (w/v) BSA, 10 mM sodium ascorbate, 10 mM cysteine, pH7.4, 0.2 mM PMSF) using a Waring blender. The suspension was centrifuged at 2,700 xg (twice) and 8,300 xg for 5 minutes to remove cell debris. Mitochondria were pelleted by centrifugation at 17,000 xg for 10 minutes, resuspended in washing buffer (300 mM sucrose, 10 mM MOPS, 1 mM EGTA, pH7.2, 0.2 mM PMSF) and carefully dispersed using a Dounce homogenizer. Isolated mitochondria were loaded onto discontinuous Percoll gradients (18%, 23% and 40% Percoll in gradient buffer [300 mM sucrose, 10 mM MOPS, pH7.2]). Percoll gradient ultracentrifugation was performed at 70,000 xg for 90 minutes. Mitochondria were collected from the 18%-23% interphase of the Percoll gradients. Percoll was removed by three cycles of pelleting the mitochondria by centrifugation at 14,500 xg for 10 minutes and resuspending the pellets in resuspension buffer (400 mM mannitol, 1 mM EGTA, 10 mM tricine, pH7.2, 0.2 mM PMSF). Washed mitochondrial pellets were finally resuspended at a concentration of 0.1 g organelle pellet per ml resuspension buffer and stored at − 80°C until further use.

### Purification of complex I from *Arabidopsis thaliana*

Purified mitochondria from *Arabidopsis thaliana* (corresponding to about 60 mg mitochondrial pellet) were sedimented by centrifugation at 14,300 xg for 10 minutes at 4°C, resuspended in membrane solubilization buffer (30 mM HEPES, 150 mM potassium acetate, 1% [w/v] lauryl maltose neopentyl glycol [LMNG]) and incubated for 5 minutes on ice. Solubilized protein complexes were separated from membrane debris by centrifugation for 20 minutes at 18,300 g and 4°C. Mitochondrial protein complexes were separated by sucrose gradient ultracentrifugation (Klodmann et al., 2010), modified). Sucrose gradients (volume: 15 ml) were prepared by a gradient mixer using 8 and 7 ml of a 1.5 M and 3 M sucrose solution (in 15 mM Tris, 20 mM KCl, 0.05 % [w/v] LMNG, pH 7.0), respectively. One mg mitochondrial protein was loaded per gradient. Centrifugation was at 146,000 xg and 4°C for 20h. Gradients were fractionated into aliquots of 500μl using a sample collector. To identify fractions containing complex I, 50 μl aliquots were analyzed by one-dimensional Blue-native PAGE (Wittig et al., 2006). Complex I was further purified by size-exclusion chromatography. Fractions containing complex I were pooled and loaded onto a Superose 6 Increase 10/300 column (GE Healthcare) equilibrated with buffer containing 30 mM HEPES-NaOH, pH 7.8, 50 mM KCl and 0.007% (w/v) LMNG. Fractions containing complex I were concentrated using a Vivaspin 500 column with a 100,000 molecular weight cutoff. To remove sucrose, the concentrated sample was resuspended in size exclusion buffer and finally concentrated to a protein concentration of 1.1 mg/ml which was used directly for cryo-EM specimen preparation.

### Purification of complex I from *Polytomella* sp

*Polytomella* sp. mitochondrial complex I was purified following the protocol for the preparation of *Polytomella* ATP synthase (Murphy et al., 2019) with modifications. Mitochondria (175mg mitochondrial protein) harvested from a *Polytomalla* culture in exponential growth phase were solubilized for 30 min at 4°C in a total volume of 12 ml buffer containing 30 mM Tris-HCl, pH 7.8, 50 mM NaCl, 2 mM MgCl_2_ and 2.9% (w/v) lauryl maltose neopentyl glycol (LMNG) to a final detergent:protin weight ratio of 2:1. Unsolubilized material was removed by centrifugation at 21,000g for 15 min at 4°. The supernatant was filtered and loaded onto a POROS GoPure HQ column (Thermo Fisher Scientific) connected to an Äkta purifier (GE Healthcare). The column was equilibrated in buffer A (30 mM Tris-HCl, pH 7.8, 50 mM NaCl, 2 mM MgCl_2_, 0.0085% (w/v) LMNG). After an initial wash with 100 mM NaCl in buffer A, complex I was eluted with a linear 100-300 mM NaCl gradient in buffer A. Fractions containing complex I were concentrated using an Amicon Ultra 4 column with 100,000 molecular weight cutoff and loaded onto a Superose 6 Increase 3.2/30 size exclusion column (GE Healthcare). Complex I was eluted in buffer B (30 mM Tris-HCl, pH 7.4, 60 mM NaCl, 0.007% LMNG) and used directly for cryo-EM specimen preparation.

### Analysis of the subunit composition of *Polytomella* complex I by two-dimensional SDS/SDS polyacrylamide gel electrophoresis

2D SDS/SDS polyacrylamide gel electrophoresis (PAGE) of *Polytomella* complex I was carried out as described (Rais et al., 2004). Briefly, purified complex I from *Polytomella* sp. was mixed 1:1 with SDS sample buffer (10% [w/v] SDS, 30% glycerol [v/v], 100 mM Tris, 4% mercaptoethanol, 0.006% bromophenolblue, pH6.8) and loaded onto a 10% polyacrylamide SDS gel containing 6M urea. After first dimension SDS PAGE, a gel lane with separated subunits of complex I was excised, washed in acidic solution (100 mM Tris, 150 mM HCl, pH2.0) and transferred horizontally onto a second dimension SDS gel (16% polyacrylamide, without urea). 2D gels were stained with Coomassie blue (Neuhoff et al., 1985).

### Protein analyses by mass spectrometry (MS)

Protein spots were excised from 2D SDS/SDS gels. Proteins were fragmented into peptides by tryptic in-gel digestion as described (Klodmann et al., 2010). Tryptic peptide mixtures were finally analyzed by coupled liquid chromatography (LC) / electrospray (ESI)-quadrupole (Q)-time of flight (ToF) mass spectrometry (MS) using the Easy nLC system (Thermo Scientific, Dreieich, Germany) and a micrOTOF Q II mass spectrometer (Bruker Daltonics, Bremen, Germany). Tryptic peptides were extracted (for details see (Klodmann et al., 2010)), resolved in solution P (0.1% formic acid, 2% acetonitrile in water) and transferred into the LC sample table. For peptide separation, a 2 cm C18 pre-column (ID 75μm, particle size 5μm, Thermo Scientific) and a 10 cm C18 analytical column (ID 75μm, particle size 3μm, Thermo Scientific) were used. A discontinuous elution gradient was applied by mixing solution A (0.1% formic acid in water) and solution B (0.1% formic acid in acetonitrile) as described (Klodmann et al., 2011). MS/MS parameters were applied as outlined before (Klodmann et al., 2011).

### Evaluation of MS data

For protein identification, the following databases were searched with an in-house Mascot server: (i) a modified *Arabidopsis thaliana* protein database, based on the TAIR database [www.arabidopsis.org] complemented with the edited sequences of mitochondrially encoded *Arabidopsis* proteins, (ii) a *Chlamydomonas reinhardtii* database and (iii) a *Polytomella* database (both downloaded from NCBI in 10/2019). In addition, a *Polytomella* protein database translated from genomic DNA was used (Murphy et al., 2019). Finally, a database integrating all sequences of complex I subunits from all databases was built and used to evaluate MS data.

### Shotgun mass spectrometry

For shotgun mass spectrometry, 50 mg purified *Polytomella* complex I was prepared by SDS PAGE and tryptic in-gel digestion as described (Thal et al., 2018). Extracted peptides were measured with an U3000 UPLC (Thermo Scientific, Dreieich, Germany) coupled to a Q Exactive Orbitrap MS system (Thermo Scientific, Dreieich, Germany) following a standard shotgun MS protocol (Thal et al., 2018).

### Reference map for 2D-separated subunits of *Polytomella* complex I

A reference map of a 2D SDS/SDS gel of *Polytomella* complex I was established at the GelMap platform (www.gelmap.de) as described (Peters et al., 2013). The map (https://gelmap.de/2062) (password: Poly-C1) summarizes all MS-based identifications of complex I subunits from *Polytomella*.

### Electron cryo-microscopy and image processing of *A. thaliana* complex I

A solution of 1.1 mg/ml purified complex I was applied onto C-flat 1.2/1.3 400 mesh copper grids (Science Services GmbH) that were glow-discharged for 45 s at 0.15 mA. Grids were frozen in liquid ethane after blotting for 4-7 s at blotforce 20 using a Vitrobot operating at 10°C and 70% humidity. Electron micrographs were collected at 300 kV in a Titan Krios G3i electron microscope equipped with a K3 detector operating in electron counting mode. The nominal magnification was 105,000x, giving a pixel size of 0.837 Å. 50-frame movies were recorded automatically with EPU software at an exposure rate of 15 e-pixel-1s-1. Particles were picked using crYOLO, motion-corrected with MotionCor2, and the CTF was estimated with CTFFind4.1.13. Further processing was performed in Relion3. For initial 3D classification to clean the dataset, particles were binned to a pixel size of 2.511 Å. After 3D refinement with C1 symmetry applied to the whole complex, particles were re-extracted at a pixel size of 0.837 Å. Two additional rounds of CTF refinement and an intermediate step of Bayesian polishing resulted in a 3D reconstruction with an overall resolution of 3.41 Å. Further multibody refinement with a soft mask around the peripheral arm, the PP domain with the CA and bridge domain, or the PD domain resulted in final resolutions of 3.21 Å, 3.39 Å or 3.43 Å, respectively. To separate the closed and open conformations, particles were aligned to the peripheral arm with a local mask applied during 3D refinement, and then 3D-classified with a soft mask applied to the membrane arm with a value of T=20 and without paricle alignment. Particle classes were further refined with a global mask, resulting in a resolution of 3.77 Å for the closed state and 3.72 Å for the open state. Final focussed 3D refinement around the PP, CA, bridge and Q domain improved the resolution to 3.72 Å and 3.69 Å.

### Electron cryo-microscopy and image processing of *Polytomella* sp. complex I

A solution of complex I at a final concentration of 1.3 mg/ was applied onto glow-discharged (0.15 mA for 45 s) C-flat 1.2/1.3 400 mesh copper grids (Science Services GmbH) and frozen in liquid ethane with a Vitrobot operating at 10°C and 70% with blotforce 20 (6 s blotting time). Electron micrographs were collected at 300 kV with a Titan Krios G3i equipped with a K3 detector in electron counting mode. 50-frame movies were recorded automatically at a pixel size of 0.837 Å and an exposure rate of 15 e^-^pixel^-1^s^-1^ with EPU software. Movies were motion-corrected with MotionCor2 and the CTF was estimated with CTFFind4.1.13. Particles were picked using crYOLO. Processing was performed in Relion3. For the first two rounds of 3D classification to clean the dataset, particles were binned to a pixel size of 2.511 Å. After 3D refinement with C1 symmetry applied to the whole complex, particles were re-extracted at a pixel size of 0.837 Å. The following 3D reconstruction resulted in an overall resolution of 3.53 Å. Additional multibody refinement with a soft mask around the peripheral and membrane arms resulted in final resolutions of 3.30 Å and 3.34 Å.

### Model building

Initial models for the *A. thaliana* and *Polytomella* sp. complex I were built using homology models for each individual subunit created with the SWISS-MODEL server (Guex et al., 2009). Homology models were then ridgid-body fitted into the cryo-EM density maps using USCF Chimera (Pettersen et al., 2004), followed by manual building in Coot (Emsley et al., 2010). Final models were refined using the phenix.real_space_refine tool in Phenix (Afonine et al., 2018). Model quality statistics were taken from the phenix.validation_cryoem tool and are summarized in table S3. For structural comparison, models were aligned using the Matchmaker tool of USCF Chimera. Water-accessible cavities were simulated with the program Hollow (Ho & Gruswitz, 2008) using an interior probe radius of 1.4 Å and a surface probe of 3.5 Å. Figures were made using UCSF Chimera and ChimeraX (Goddard et al., 2018).

