## Supplementary figures and tables for "A ferredoxin bridge connects the two arms of plant mitochondrial complex I"

### List of Supplementary figures and tables

Supplementary Figures

Supp. fig. 1: Purification of complex I from *Arabidopsis*.
Supp. fig. 2: Purification of complex I from *Polytomella*.
Supp. fig. 3: Cryo-EM of *Arabidopsis* complex I. Particle classification and processing scheme.
Supp. fig. 4: Cryo-EM of *Arabidopsis* complex I. Orientation distribution and FSC curves.
Supp. fig. 5: Cryo-EM of *Polytomella* complex I. Particle classification and processing scheme.
Supp. fig. 6: Cryo-EM of *Polytomella* complex I. Orientation distribution and FSC curves.
Supp. fig. 7: Non-assigned density of *Arabidopsis* complex I.
Supp. fig. 8: Subunit composition of *Arabidopsis* complex I as revealed by 2D SDS / SDS PAGE.
Supp. fig. 9: Alignment of *Arabidopsis* P1 and bovine SGDH.
Supp. fig. 10: Cofactors and lipids in *Arabidopsis* complex I.
Supp. fig. 11: Subunit composition of *Polytomella* complex I as revealed by 2D SDS / SDS PAGE.
Supp. fig. 12: Subunits of *Polytomella* complex I identified by mass spectrometry.
Supp. fig. 13: Complex I of *Polytomella*. Peptides identified by coupled LC-ESI-MS/MS mass spectrometry.
Supp. fig. 14: Partially assembled peptides of complex I subunits from *Polytomella* sp.
Supp. fig. 15: Densities of *Polytomella* complex I assigned by homologue subunits.
Supp. fig. 16: Non-assigned densities of Polytomella complex I.
Supp. fig. 17: Cofactors and lipids of *Polytomella* complex I.
Supp. fig. 18: γCA/γCAL proteins of the heterotrimeric γCA domain in *Arabidopsis*.
Supp. fig. 19: Structures of γCA and γCAL subunits in *Arabidopsis* complex I.
Supp. fig. 20: γCA/γCAL subunits in the heterotrimeric γCA domain of *Polytomella*.
Supp. fig. 21: Structures of γCA and γCAL subunits in *Polytomella* complex I.
Supp. fig. 22: *Arabidopsis* mitochondrial C1-FDX.
Supp. fig. 23: Comparison of *Arabidopsis* C1-FDX to *Arabidopsis* mitochondrial FDX1 and FDX2.
Supp. fig. 24: Comparison of the bridge domain of *Arabidopsis* and *Polytomella* complex I to the human iron sulfur cluster (ISC) assembly core complex.
Supp. fig. 25: Loop conformations in the closed and open state of *Arabidopsis* complex I.

Supplementary Tables

Supp. tab. 1: Nomenclature of complex I subunits in *A. thaliana*, *C. reinhardtii* and other model species.
Supp. tab. 2: EM Statistics *Arabidopsis thaliana* complex I.
Supp. tab. 3: EM Statistics *Polytomella sp.* complex I.
Supp. tab. 4: *Arabidopsis thaliana* complex I map identifiers and statistics.
Supp. tab. 5: *Polytomella* complex I map identifiers and statistics.
Supp. tab. 6: *Arabidopsis thaliana* complex I model identifiers and quality statistics.
Supp. tab. 7: *Polytomella* complex I model identifiers and quality statistics.

#### Supplementary Figures

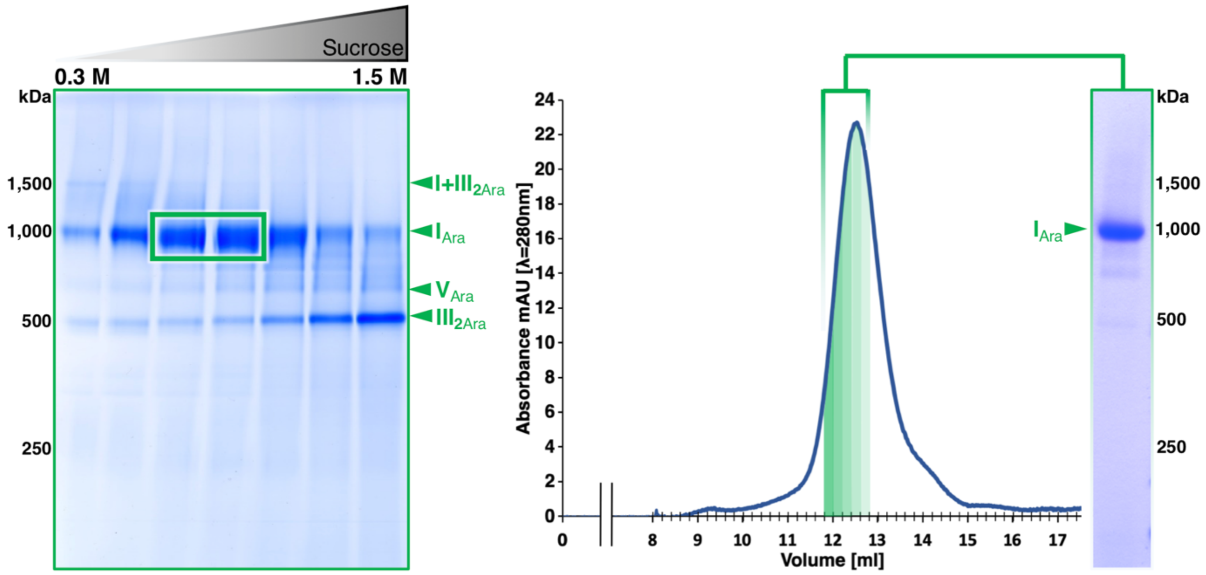

**Supp. fig. 1:** Purification of *Arabidopsis* complex I. Mitochondria isolated from *Arabidopsis* cell suspension culture were solubilized with 0.75 g LMNG / g mitochondrial protein. Protein complexes were separated by sucrose gradient ultracentrifugation and fractions were analysed by 1D blue native PAGE (left). For cryo-EM analysis, samples containing complex I were loaded onto a size-exclusion column (right). Peak fractions were pooled and concentrated to a final concentration of 1.1 mg/ml for grid freezing. Ara, *Arabidopsis thaliana*; I, complex I; $III_2$, dimeric complex III; I+$III_2$, supercomplex formed by complexes I and $III_2$; V: complex V.

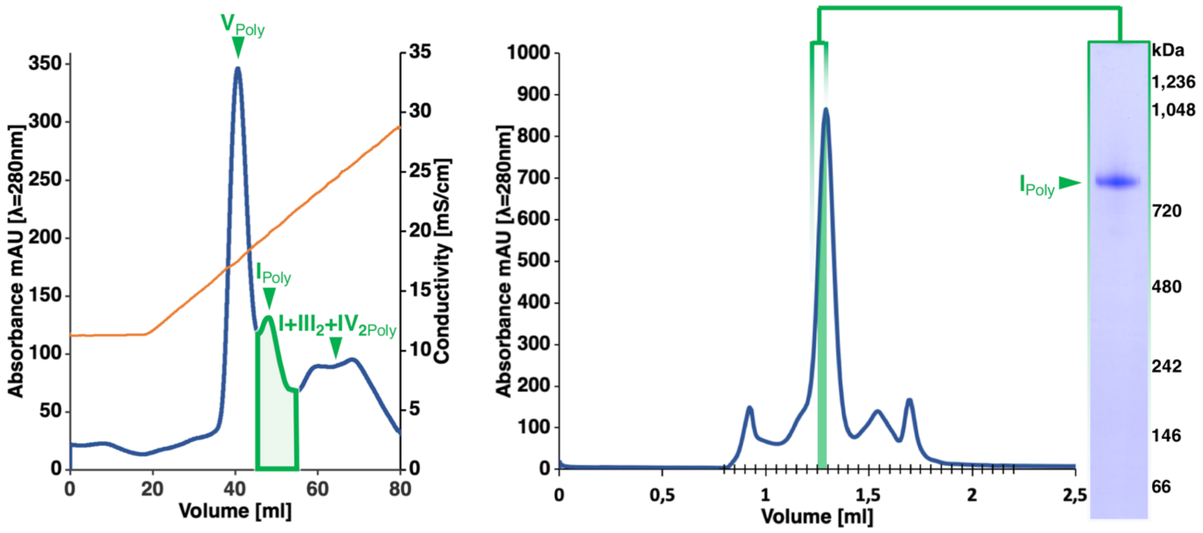

**Supp. fig. 2:** Purification of complex I from *Polytomella*. Isolated mitochondria from *Polytomella sp.* were solubilized in 2 g LMNG / g mitochondrial protein. Protein complexes were separated by ion-exchange chromatography (left). Fractions containing complex I were pooled, concentrated and loaded onto a size-exclusion column (right) for cryo-EM analysis. The peak fraction used for grid freezing had a final concentration of 1.3 mg/ml. For details see Materials and Methods. Poly, *Polytomella sp.*; I, complex I; I+$III_2$+$IV_2$, supercomplex formed by complexes I plus $III_2$ and $IV_2$; V: complex V.

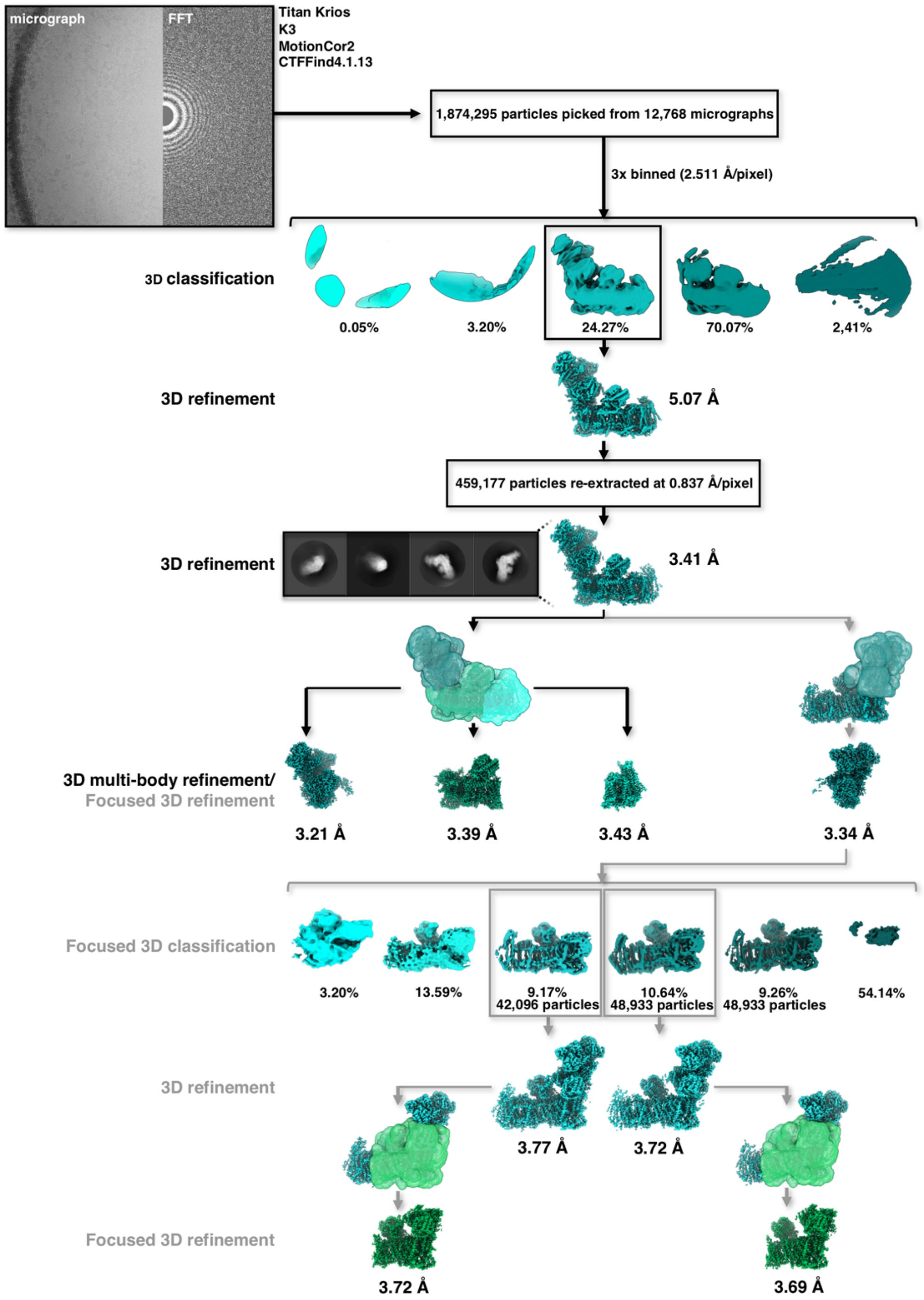

**Supp. fig. 3:** Cryo-EM of *Arabidopsis* complex I. Particle classification and processing scheme. Images were recorded with a Titan Krios G3i microscope equipped with a K3 camera in counting mode at an exposure rate of 15 $e^{-}pixel^{-1}s^{-1}$. In total, 12,768 50-frame movies were collected automatically using EPU software. After motion correction and defocus estimation, particles were picked using crYOLO and binned to a pixel size of 2.511 Å. Further processing was performed in Relion3. After initial 3D classification and 3D refinement, particles were re-extracted at a pixel size of 0.837 Å. 3D refinement applying a soft mask for the whole complex resulted in a global resolution of 3.41 Å after two rounds of CTF refinement and Bayesian polishing with C1 symmetry. Further multibody refinement with a soft mask around the peripheral arm, the $P_P$ domain with the CA and bridge domain or the $P_D$ domain resulted in final resolutions of 3.21 Å, 3.39 Å and 3.43 Å, respectively. For separation of closed and open states of the complex, particles were aligned by their peripheral arm with a local mask during 3D refinement and then 3D classified with a soft mask applied to the membrane arm using a value of T=20 without particle alignment. Particles were further refined with a global mask resulting in a resolution of 3.77 Å for the closed state and 3.72 Å for the open state. Final focused 3D refinement around the $P_P$, CA, bridge and Q domain improved the resolution to 3.72 Å and 3.69 Å, respectively.

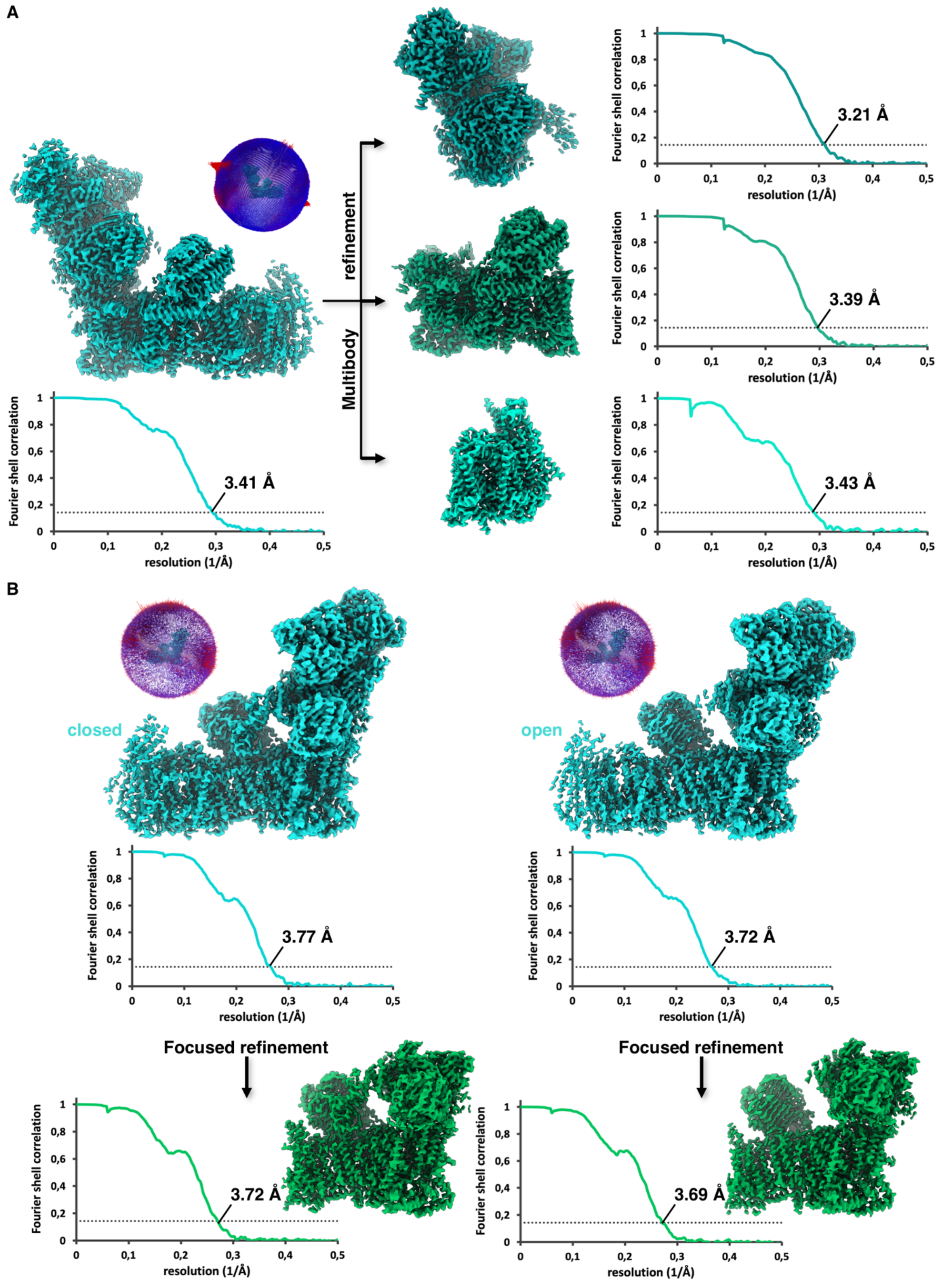

**Supp. fig. 4:** Cryo-EM of *Arabidopsis* complex I. **A:** Orientation distribution and FSC curves for the multi-body refinement. **B:** Orientation distribution and FSC curves for the open and closed state. Resolution was estimated by the FSC 0.143 criterion.

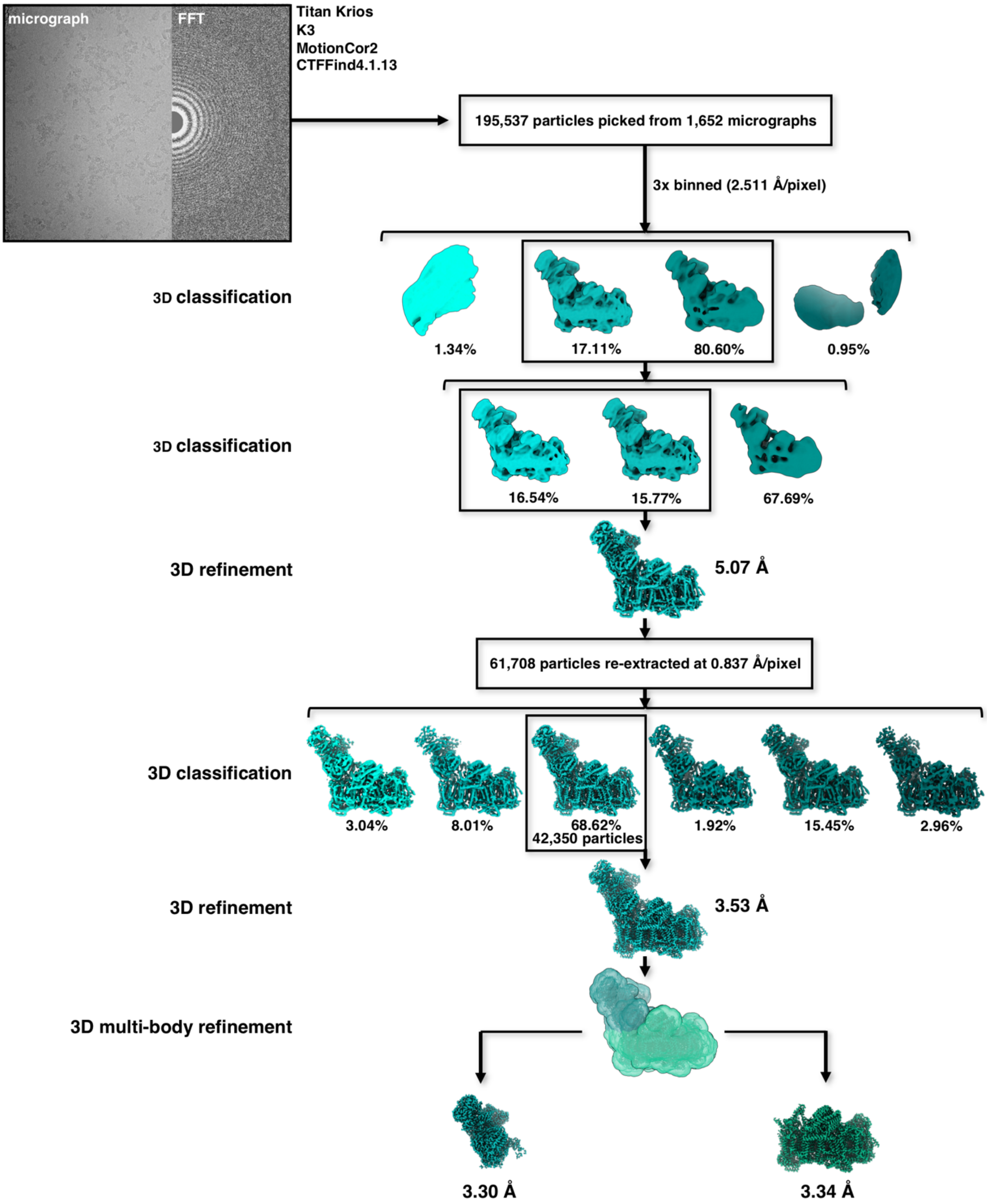

**Supp. fig. 5:** Cryo-EM of *Polytomella* complex I. Particle classification and processing scheme. Images were recorded with a Titan Krios G3i microscope equipped with a K3 camera in counting mode at an exposure rate of 15 $e^{-}pixel^{-1}s^{-1}$. 1,652 50-frame movies were collected automatically using EPU software. Particles were picked from motion corrected and defocus estimated micrographs using crYOLO. Further processing was performed in Relion3. After initial binning to a pixel size of 2.511 Å, two rounds of 3D classification and further 3D refinement, particles were re-extracted at a pixel size of 0.837 Å. After further 3D classification of the whole complex, 3D refinement resulted in a global resolution of 3.53 Å with C1 symmetry. Further multibody refinement with a soft mask around the peripheral arm and the membrane arm resulted in final resolutions of 3.30 Å and 3.34 Å, respectively.

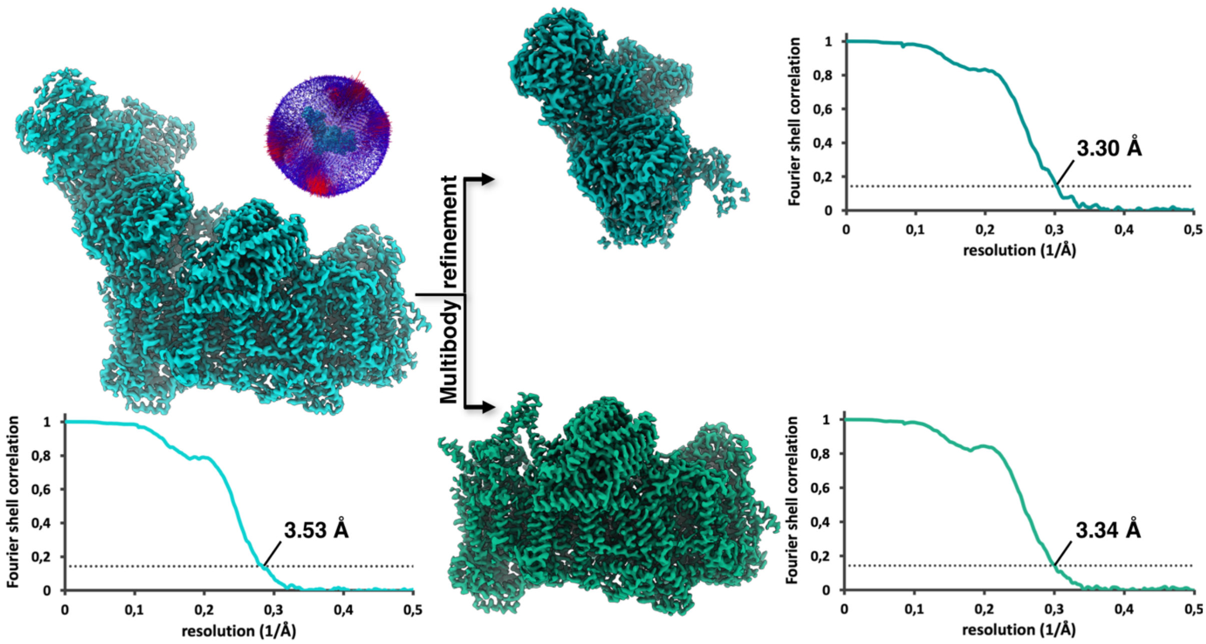

**Supp. fig. 6:** Cryo-EM of *Polytomella* complex I. Orientation distribution and FSC curves. Resolution was estimated by the FSC 0.143 criterion.

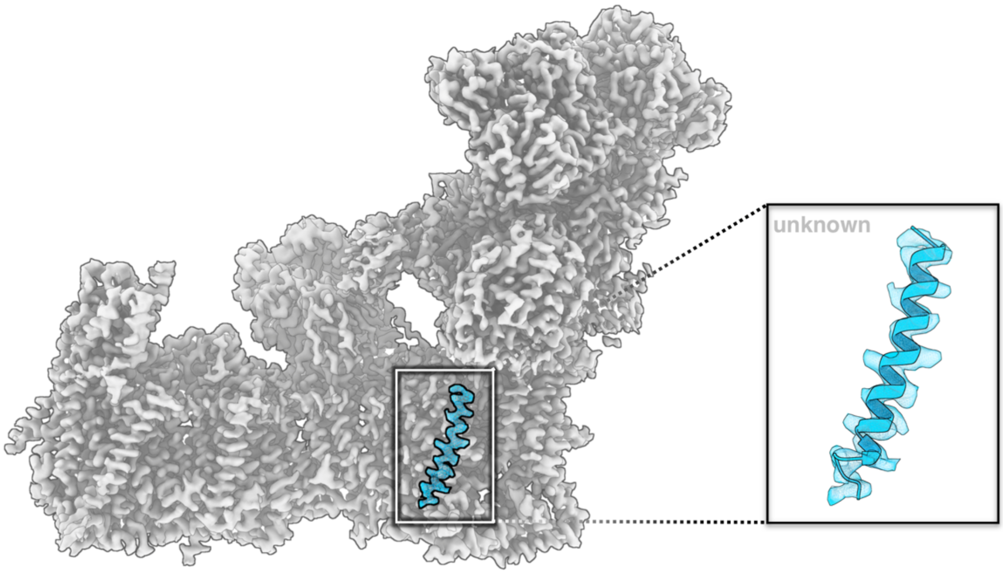

**Supp. fig. 7:** Non-assigned density in the *Arabidopsis* complex I membrane arm at the $P_P$ domain.

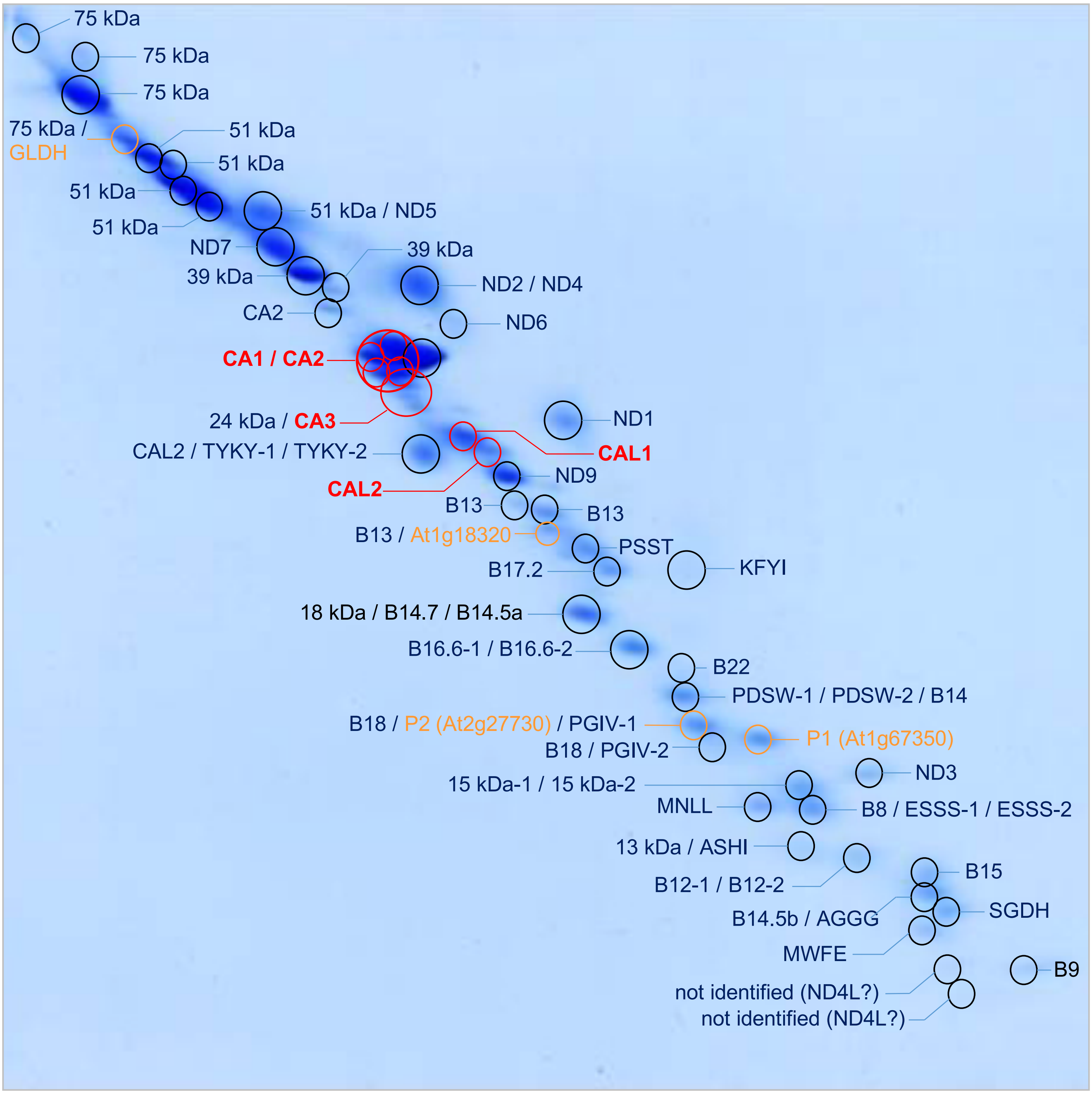

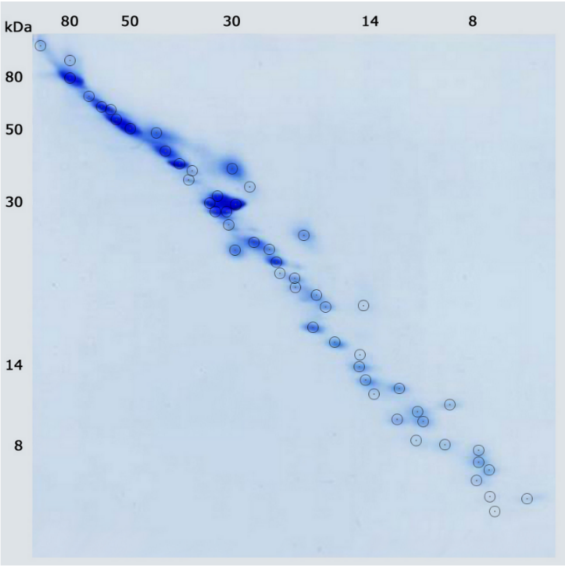

**Supp. fig. 8:** Subunit composition of *Arabidopsis* complex I as revealed by 2D SDS / SDS PAGE (Figure from Peters et al. 2014). Isolated complex I from *Arabidopsis* was analyzed by 2D SDS / SDS PAGE. Proteins were identified by LC-ESI-MS/MS. Subunits labelled red: carbonic anhydrase subunits. Subunits labelled orange: other complex I subunits not present in complex I of animals and fungi.

https://gelmap.de/235

```
Ara. P1          1 -----------------------------MGFIM-EFAENLV-----------LRLMENP 19
B. taurus SGDH   1 MAAMSLLHRASVSAVAALSGRRLGTRLGFGGFLTRDFPKTVAPVRHSGDHGKRLFIIKPS 60
                                                 **:  :*.:.:.           * ::: .

Ara. P1         20 EERDRK---------------------------AREHIYEMHERCKKIKEMWALPIRPYG 52
B. taurus SGDH  61 GFYDKRFLKLLRFYILLTGIPVAIGITLINVFIGEAELAEIPEGY--VPEHWEYFKHPIS 118
                      *..                           .   : *: *    : * *    .* .

Ara. P1         53 FW-----------TFERHNAQLRWDPQ-----ISQVAGRR-------DPYDDL--LEDNY 87
B. taurus SGDH 119 RWIARTFFDGPEKNYERTMAILQIEAEKAELRLKELEVRRLMRARGDGPWFHYPTIDKEL 178
                    *           .:**  * *. :.:     :.::  **       .*:     ::.:

Ara. P1         88 TPPSSSSSSSD 98
B. taurus SGDH 179 IDHSPKATPDN 189
                      *..::..:
```

**Supp. fig. 9:** Alignment of *Arabidopsis* P1 and bovine SGDH.

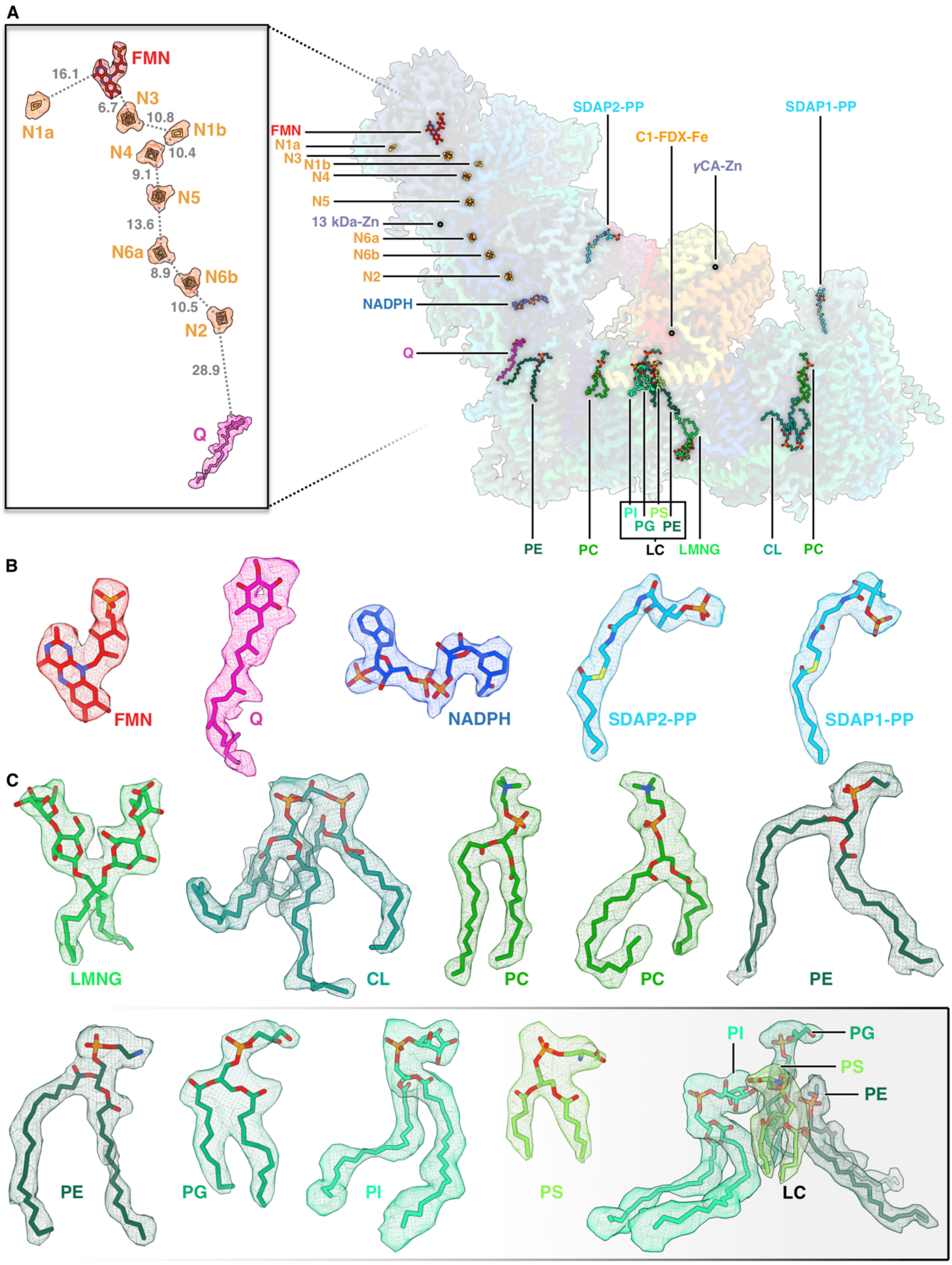

**Supp. fig. 10:** Cofactors and lipids in *Arabidopsis* complex I. FMN: Flavin mononucleotide; N1a, N1b, N2, N3, N4, N5, N6a, N6b: FeS clusters in the peripheral arm; Q: ubiquinone; NADPH: Nicotinamide adenine dinucleotide phosphate; SDAP1-PP, SDAP2-PP: 4'-phosphopantetheine bound to SDAP1 and SDAP2 subunits; LMNG: Lauryl Maltose Neopentyl Glycol; CL: cardiolipin; PC: phosphatidylcholine; PE, phosphatidylethanolamine; PG: phosphatidylglycerol; PI: phosphatidylinositol; PS: phosphatidylserine; LC: lipid complex.

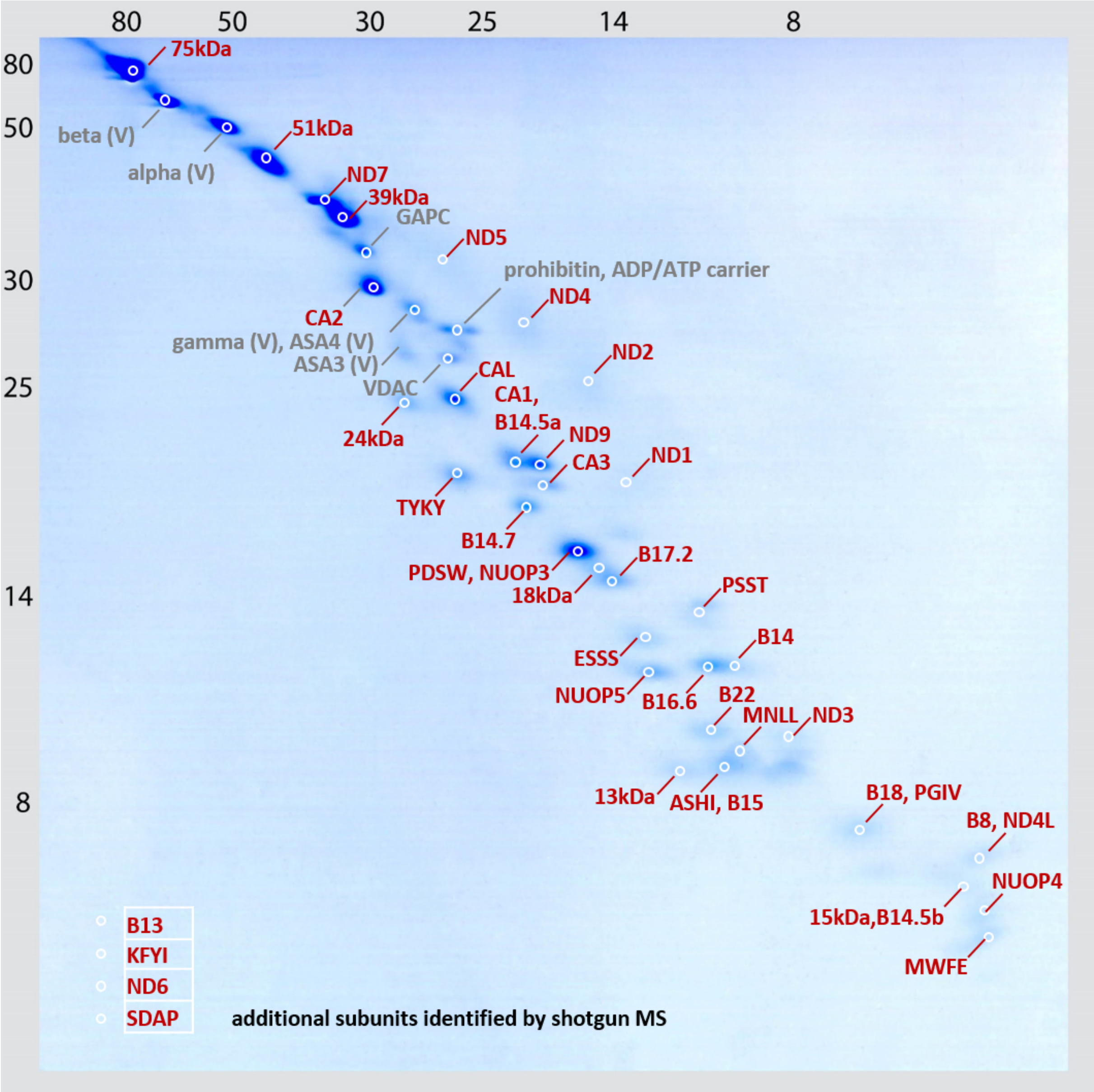

**Supp. fig. 11:** Subunit composition of *Polytomella* complex I as revealed by 2D SDS / SDS PAGE. Proteins were identified by mass spectrometry. For subunit assignments see supplementary Figure S12. Additional complex I subunits were identified by direct shotgun proteome analyses of the purified protein complex. A GelMap was generated, which can be accessed at www.gelmap.de/2062 (password: Poly-C1). Contaminating proteins that are not part of complex I are labelled in grey. Of these, four belong to complex V (ATP synthase, V). Furthermore, a voltage dependent anion channel VDAC, an ADP/ATP carrier, a prohibitin and a glyceraldehyde-3-phosphate dehydrogenase (GAPC) were identified.

| Subunit | homolog in *C. reinhardtii* | homolog in *A. thaliana* | Localization |
|---|---|---|---|
| 15 kDa | Cre12.g511200 | At3g62790, At2g47690 | membrane |
| ASHI | Cre01.g007850 | At5g47570 | membrane |
| B14 | Cre12.g555250 | At3g12260 | membrane |
| B14.5b | Cre13.g571150 | At4g20150 | membrane |
| B14.7 | Cre14.g617826 | At2g42210 | membrane |
| B15 | Cre03.g204650 | At2g31490 | membrane |
| B16.6 | Cre16.g664600 | At1g04630, At2g33220 | membrane |
| B18 | Cre06.g278188 | At2g02050 | membrane |
| B22 | Cre11.g467668 | At4g34700 | membrane |
| ESSS | Cre05.g240800 | At2g42310, At3g57785 | membrane |
| KFYI | Cre17.g725400 | At4g00585 | membrane |
| MNLL | Cre06.g267200 | At4g16450 | membrane |
| MWFE | Cre10.g459750 | At3g08610 | membrane |
| ND1 | AAB93446 | AtMg00516 / AtMg01120 / AtMg01275 | membrane |
| ND2 | AAB93444 | AtMg00285 / AtMg01320 | membrane |
| ND3 | Cre08.g378900 | AtMg00990 | membrane |
| ND4 | AAB93441 | AtMg00580 | membrane |
| ND4L | Cre09.g402552 | AtMg00650 | membrane |
| ND5 | AAB93442 | AtMg00060 / AtMg00513 / AtMg00665 | membrane |
| ND6 | AAB93445 | AtMg00270 | membrane |
| NUOP3 | Cre02.g100200 | At3g07480 | membrane |
| NUOP4 | Cre08.g378550 | n.i. | membrane |
| NUOP5 | Cre08.g378050 | n.i. | membrane |
| PDSW | Cre12.g555150 | At3g18410, At1g49140 | membrane |
| PGIV | Cre07.g333900 | At3g06310, At5g18800 | membrane |
| SDAP | Cre16.g673109 | At1g65290, At2g44620, At5g47630 | membrane |
| CA1 | Cre12.g516450 | At1g19580 | CA domain |
| CA2 | Cre09.g415850 | At1g47260 | CA domain |
| CA3 | Cre12.g516450 | At5g66510 | CA domain |
| CAL | Cre06.g293850 | At5g63510, At3g48680 | CA domain |
| 13 kDa | Cre03.g178250 | At3g03070 | peripheral |
| 18 kDa | Cre03.g146247 | At5g67590 | peripheral |
| 24 kDa | Cre10.g450400 | At4g02580 | peripheral |
| 39 kDa | Cre10.g434450 | At2g20360 | peripheral |
| 51 kDa | Cre10.g422600 | At5g08530 | peripheral |
| 75 kDa | Cre12.g535950 | At5g37510 | peripheral |
| B8 | Cre16.g679500 | At5g47890 | peripheral |
| B13 | Cre12.g484700 | At5g52840 | peripheral |
| B14.5a | Cre12.g484700 | At5g08060 | peripheral |
| B17.2 | Cre11.g467767 | At3g03100 | peripheral |
| ND7 | Cre09.g405850 | AtMg00510 | peripheral |
| ND9 | Cre07.g327400 | AtMg00070 | peripheral |
| PSST | Cre12.g492300 | At5g11770 | peripheral |
| TYKY | Cre12.g496750 | At1g79010, At1g16700 | peripheral |

**Supp. fig. 12:** Subunits of *Polytomella* complex I identified by mass spectrometry. First column: abbreviated subunit names (bovine nomenclature; Walker et al. 1992). Second column: accession numbers of corresponding genes in *Chlamydomonas*. Third column: Accessions of homologs in *Arabidopsis*; fourth column: assignments to complex I arms (membrane arm, peripheral arm, γ-carbonic anhydrase domain).

**15 kDa**
APMITLR
APMITLRK
KEPMIALR
SGFGLQSGTAR
SGFGLQSGTSR
**ASHI**
EFLALFK
QSGLLTPIY
QYPSEVR
NGFVLNDYSQPDYEEMK
NGFVLNDYSQPDYEEMKR
RQSGLLTPIY
**B14**
AAAPAAPVTCQK
AYPLVHSK
LEEITSLK
VTDLLIFK
FAYVAK
LEEITSLKEMR
NIYKNCAR
SSNSAFLDSFYEK
VNSPVTDSR
AAAPAAPVTCQKAR
CLPFIHR
FRVNSPVTDSR
KSSNSAFLDSFYEK
LHKLEEITSLK
**B14.5b**
LFSAY
GLQVTEEHKK
LLDEVQDTYYLEHLK
LLDEVQDTYYLEHLKR
**B14.7**
AEEVTVDFSK
KLAPIVINK
AEEVTVDFSKEPSASSLK
EPSASSLK
GYGTPYGFENLDDAVPTR
LAPIVINK
RTYNIIK
TPIYGK
TYNIIK
TYNIIKK
CGFALK
FGFALK
GYGTPYGFENLDDAVPTRNPIR
SYTSGMVAGAATGAVFAIGR
TPIYGKR
TAYATLLGAIFCATDALYENYSGK
**B15**
SNFVIPDER
NYYFSDSK
SNFVIPDEREFY
YTPNIFSPETPMDR
SFSHVPK
**B16.6**
GFPGLK
LEAEEVYTAR
NVPGWK
SIKDLPK
VQDVPPPGGFPSIR
WLPPVESFDVRPLI
VQDVPPPGGFPSIRIER
YLEHQR
IMQDIFQK
ASFIK
IIELK
LKGECSYK
SILHPILQAEWDLR
**B18**
CSYIEFK
VAIAIEEK
CSYIEFKR
INCAHELHHYNK
RVAIAIEEK
VAIAIEEKK
YGHPLTWR
**B22**
EIVNWSYGR
FVPNPAYIDEFK
LVLHGEDLLAR
ASEAVLR
HEFYPR
MEFEANKELSDR
QGDTVLLY
ASEAVLRLPHK
ELSDREQIR
HWEPTIR
KLVLHGEDLLAR
SEFVGGTVYGVWPQHCK
**ESSS**
AIYEELEA
AKDEYAVDMK
VLLGLHSGR
YGEIPLPNGQPR
AGGASGHPNGSFWSEGTQVGLNGFK
DEYAVDMK
DLNMIEEEAYDLIMMR
LSIVLK
VPADLK
**KFYI**
GWIPSQLWNDNVPTPVDYR
SHYCPPDFHYSR
**MNLL**
NVPAALVPEHY
PINPYDVPSSPAYHR
LQEPSYPVIDK
PSWNQNK
**MWFE**
EGKPSWNQNKDQFQR
**ND1**
GGVSSQCVDASR
HVAFVFSTLPK
ITANENQR
LMYLVSPVK
AVMGSLQR
DGNGFPCLNLMETASQAK
DGNGFPCLNLTETASQTK
LKGGVSSQCVDASR
LVFYLR
YAFLGCLR
YDMFMQIGWK
LDVISWNPLVLATVVTGIGYM
MGPAVSGAFGILQPFWDGFK
**ND2**
HVAFVFSTLPK
ITANENQR
LMYLVSPVK
**ND3**
MGALEWRE
YFVEGEHVVQAEANR
MGALEWR
**ND4**
GGISSQCVDASR
GGVSSQCVDASR
LVFYLR
LKGGVSSQCVDASR
LDVISWNPLVLATVVTGIGYM
AANEVEALNLLK
**ND4L**
AANEVEALNLLK
**ND5**
ENNNFLNR
GGFDVFYAR
IFTGSSLSQNMTTNLPAHIK
SLIVNK
ATLFLSAGLSIAK
QGSPTLSFATTIASLNLLGFPELGGFYSK
TSTSLEWLLPATPAFHTFSEVPVLR
VLAQLYLFPYGNGR
**ND6**
IQVSTLSTK
**NUOP3**
ALIGTPLNK
ETSFFASYLTLNK
LETGPFIE
LVVQDYQR
MFDRLETGPFIE
TDDAEAIK
YVENVDLK
YVENVDLKTDDAEAIK
TDDAEAIKLMFK
WSFNGHHNVK
EMDNMTIGFGPIKPWHITPK
TVFEDTITINVLDYDGK
**NUOP4**
ADTSMDHVFGDSTNK
**NUOP5**
AQAEVDTLSAAGR
TEVDPTGK
DPNELQK
DSDSRDEYAINAEEPVIYIR
LGPLSPF

RDPNELQK
DEYAINAEEPVIYIR
**PDSW**
EEPHVYYHNNNFR
ELGLNNPFPPSSPR
IFEDELDK
DVGFIYQNNK
EVLDIVQSK
QLIHFEK
SEKEEPHVYYHNNNFR
NSEWARPEEPILHNLK
VASNLERGPNAR
IFEDELDKCMR
RDVGFIYQNNK
**PGIV**
QAAFEAAFPLSE
APEEFKDYAK
AQHDWPEDCKPESEAIVR
NEKEHNAILECK
VPYSHELYAMAK
EHNAILECK
EKQAAFEAAFPLSE
NDPEKEQFQFIDR
**SDAP**
EQNLAAGIK
**CA1**
ASVWYNAVVR
EIQYLSTLPER
LQGNYSFK
ETGLALETLGCK
FGFALK
HSTLTQVQYK
HSTLTQVQYKEPSLGK
NVSIGHGAVLK
YVESSAEHSAMMELLSLK
NTFIAPSALVAGDVIIGDK
TLVWR
GCTIGDNVLVGMNAVISEK
LRDLRPK
**CA2**
AESTFIAPNANVLGDVK
NLAVHQLYIFDPQTQLAAR
NYPPSYLYPFR
VESGSIVGAGSIVPPNTVIPAGQVWV
GNPAK
AGIVAPIPASAQR
EIVLPSK
HPKPEGVIEK
NVLPEENGFIASSANNYDLLGQQHK
NVLPEENGFIASSANNYDLLGQQHKFE
NSK
SSIWYGAVLR
VFEEMLVEEEIAK
VFEEMLVEEEIAKDR
VLFNLGSLFR
VLGVK
YNPAAAPK
DRELLEDK
ELLEDK
ELLEDKNLAVHQLYIFDPQTQLAAR
EIVLPSKAESTFIAPNANVLGDVK
**CA3**
APSVGESTFVAPSALVSGDVIIGEK
ATVEDNTILAPGSYVPEDVVVK
LQGNYSFR
ETGLALETLGCK
HTPLTQYQFK
LLELK
SGELWSGSPAQK
SSVLYNAVVR
CGFAIK
ELALFNTLSVGATELAADHAVIMK
NLDEKELALFNTLSVGATELAADHAVIM
K
TIVWR
SVTIGEGSTISDNAYVGSSSEFSPETVIGS
NVSVGSGAVLK
**CAL**
GDLNTIR
IEDEWYNR
IGDNCIVGAR
LVDYK
ESAEHYYDIATGFR
IGENSTILDK
SVVCEGAIMHPESVLAPGSVLAPGR
TEFNILNK
TEFNILNKQPSK
VSIFFGAVVR
ADDVFIAPNAVVCGDVDLYQK
DAAELEK
DAEAYR
EYNWVNYR
LVDYKEYNWVNYR
NETFDHGSAWR
QIPSGELWAGSPAK
QKLVDYK
QRTEFNILNK
NVNFYDAPNGPAVEQVK
QPSKADDVFIAPNAVVCGDVDLYQK
**13 kDa**
GTSYENPAVCK
NKIPETVGGAAFK
SPMDYINEVEPIK
GTSAVGLGDGLYSHTSK
IPETVGGAAFK
**18 kDa**
AASQSGLGR
AIEATCQYR
GVPDLAHLPSNR
HGWEYVVDEPNPR
STLLFYTK
APQWK
AVADVEIDFNNR
STLLFYTK
APQWK
AVADVEIDFNNR
FIMSINSLLEAVK
GVPDLAHLPSNRS
IVFENLSK
KGVPDLAHLPSNR
SSLFGR
IFCPAK
KAIEATCQYR
WENPLMGWTSTADPLENVGR
YVGVDHLYK
KYVGVDHLYK
YVGVDHLYKEQATK
**24 kDa**
AEAIMSNYPANYK
IEEAITK
YHVCICGTTPCR
IATILEVPPIK
KAEAIMSNYPANYK
QTAEPAGAVVGDK
ASAMIPLLDLAQQQNGGVVSLAVMNR
EIEAISLAK
IMSNYPSNYK
NGVEGFSYNYYEDLTPQDAVSILEK
TSAMIPLLDLAQQQNGGFVSLAVMNR
VAQILEVPPIK
VYEVATFFTMFNR
LQGAQKIEEAITK
NGVEGFSYNYYEDLTPQDAVNILEK
QTAEPAGAVVGDKWVPSSGENTLMGE
LPGPYCR
WIPSSGEQTLMGELPGPYCR
WVPSSGENTLMGELPGPYCR
**39 kDa**
AVGDEAIMK
DVHVDFPTR
GNQVVCPYR
DLSISPAK
FIQVSEMGADVSHASR
IVAESGKVER
LSNDDTVNLR
MLYKPLDTLR
NYSYK
QIYDHLIDTLR
RGNQVVCPYR
SNVVINCVGIR
TYYLGGPETLTMR
VGGYDWGDMNK
VQPTYILDVADAVVK
VSSNIPEHIR
VSSNIPEHIRK
VTEGLAIEGVR
YELAK
YIPDATIIR
YYNLK
KYYNLK
LSNDDTVNLRYELAK
NYSYKDVHVDFPTR
**51 kDa**
DTDVIDAIAR

GYLGK
LVEGCLVAGTAMGAR
TTFGGLRDQDR
WSFMPK
AGYIYIR
AVAEAYAK
GDWYMTK
GEFVNER
GGAGFASGLK
HEPHKLVEGCLVAGTAMGAR
IFTNIYGR
KMCDDVIMDFDALR
LSYFYK
MCDDVIMDFDALR
SEMEDRIK
TAQSGLGTAAVIVMNK
TAQSGLGTAAVIVMNKDTDVIDAIAR
DWIIDQIK
DWIIDQIKK
EGTGWLYDIMSR
ELIER
GGWDNLLAIIPGGSSVPLLPK
GPEWFSSFGR
GRDWIIDQIK
LEEIDMLWEITK
RGPEWFSSFGR
SIIER
TTFGGLR
ASDGRPSYLVVNGDESEPGTCK
GRGGAGFASGLK
LKPPFPAGMGLYGCPTTVTNVETVAVSPTILR
QIEGHTICALGDAAAWPVQGLIR

**75 kDa**

AESALWLNGQYFK
ANVVLPGAAYTEK
IKTDTPVIK
LSIAGNCR
NFTVLQACDAAGVDVPR
ADVILLVGTNPR
ALSEVVGQQLPYDSQPQVR
ASLFANTEGR
DDREAVLK
DSEVGTYIEK
FCYHQR
FESPVFNAR
FIEYK
FIEYKR
FQYDGLKR
FTHEVAGTSELGITGR
GAFFEALK
GLENATWSAAFDAIR
GRDSEVGTYIEK
GTETIDVSDALGSNIK
ISLNEIQTK
KVFLDGAK
LADAESMIALK
LADAESMIALKDLFNK
LAEVAPHFAEIGK
LGSGNLIHEDGSATLSADVR
LNDAINEEWLSDK
LNDAINEEWLSDKGR
LNEAVNEEWLSDK
LNEAVNEEWLSDKGR
LNTPLVK
NLGPLIK
NPVVIVGSSVLR
SSYIANTTIASIEK
STASLATNISNYYMTDAISR
TAVPVLGDAR
TDTPVIKK
TVNDLVDAAGVVK
VAALDIGFVPSASAR
DIFNK
DLFNK
DSEIGTYVEK
EGWNGFNVLHDNASR
FQYDGLK
GRDSEIGTYVEK
GVKDLVAK
MEVFVNGEAVVVPK
PVASCAFPAGPGMK
PVASCAMPAGPGMK
PYAFTAR
QRLNTPLVK
SWELK
TAVPVLGDAREDWK
TDTPVIK
VDLTYAYQHLGADVAALESLASGK
VFLDGAK
VGLVGEK
VVYLLGSDDFKDEEIPADAFVIYQGHHGDK
APKPVASCAFPAGPGMK
AVADKNLGPLIK
LHSSELSGNVIDLCPVGALLSKPYAFTAR
SWELKGTETIDVSDALGSNIK
TAIAGAKGNELK

**B8**

ESNAAQASLTAR
AANPLLPILVR
EFVLGQYAELK
KVSIENDSASGILSK
LEGLVK
VSIENDSASGILSK

**B13**

AIEATCQYR
SVPLPK
VVDTGSPIK
AVADVEIDFNNR
AVADVEIDFSNR
EGNDLYR
EMIESGVIK
FIMSINSLLEAVK
FTQANSEVSALLGR
IPSAVGYQPTLATDLGGLQER
KAIEATCQYR
LVLEVAQHMGDNTVR
LVSEIK
MLNPNVIGAEHYNVAR
SIAELGIYPAVDPLDSTSR
SVPETAEYR
SVPETAEYRK
TREGNDLYR
VALTGLTVAEYFR
VLQDYK
VSEELKDIPSLDK
YDDLPEMAFYMVGDIK
YVGVDHLSK
AKFIVSLNSLLEAVK
DIEGQDVLLFVDNIFR
FDGELPSILSSLEVEGHSVR
FIVSLNSLLEAVK
GSITSVQAVYVPADDLTDPAPATTFAHLDATTVLSR
KYVGVDHLSK
NLQDIIAILGMDELSEEDKLTVAR
VCGENNSDAAIEEVLDAHLEELIAESK
YVGVDHLYK
AEALSSETVVLNEEGK
CIAMDSTDGLVR
CTLVYGQMNEPPGAR
EVVIEEDDGLEEDFKAEALSSETVVLNEEGK
YVGVDHLSKEQGTK

**B14.5a**

MAEALK
LYAGMPTK
AVVPHDVPK
IPEVMPHR
SLPDFEYYLPFGLTYR
TPFDYSK
TVQSIFYSVGLKEPWK
VFGSAPLKPADVK

**B17.2**

GSWFNPAK
PQSGAQVGIDK
YLNNLVSTSGK
KVEIWQPPQV
VEIWQPPQV
WIVFADK
WIVFADKFDYK
YLNNLVSTSGKK
SLAGFFVSWEFAK
HNNVVRPQSGAQVGIDK

**ND7**

GEFGVYLVSR
LFEFYER
VLFSEITR
VSNFTINFGPQHPAAHGVLR
DIYDWSR
GSGIDWDLR
GSGIDWDLRK
IIYQCLNEMPDGLYK

LFSEGYHVPAGETYR
LLEYK
LVLEMDGEIIK
LVLEMDGEIIKR
MDFNVPIAGHGDCYDR
MHAAYFR
NQPYDVYGR
QSMESLIHHFK
SPGYAHLQMLDMVAK
TYNQGIPYFDR
VDEMEELLTGNR
VGGVAQDLPIGLLR
AFFLLK
EKLFEFYER
LDEMEELLTGNR
LVLEMQGEIIMR
QSMEALIHHFK
TIDVGLVTAQQAWDWGCSGPILR
GAMLADVVTIIGTLDVVFGEIDR
KNQPYDVYGR

**ND9**

IWNAANWFER
NFGDNYLTDYIIK
VISEPLELTQEFR
DHVNLQYK
DYIAYDNIK
FEVVYHLLSPR
MTDKDYIAYDNIK
TLIDITAVDYPER
VLTDYGFTGHPLR
YFDFSSPWDTLSR
VVSEPLELTQEFR
YDYGK
NSQALLGR
VLTDYGFTGHPLRK

**PSST**

GADFIVSK
FGIIFRPSPR
KVYDQMPEPK
NMYYR
NTQLWWNK
QSDVMIVAGTLTNK
VDTVVNWAR
VYDQMPEPK
GNFLVCGEGQGGEGGGEK
IRNSGNGAAK
MAPALR
SKNTQLWWNK
VVPVDVYVPGCPPTAEGLLYGLLQLQK
YDLDR
AGSMWPMTFGLACCAVEMMHAGAS
R
TNISLVIDASK
VVMLSQGSSK

**TYKY**

EELLYDK
RYPTGEER
YSSEWEQDPTFK
FIEYKR
FLEKYR
YDIDMTK
YPTGEER
EELLYDKQK
FRGEHALR
GMMYSASGFFDDK
INSVAKR
LCEAICPAQAITIEAEER
LCEAICPAQAITIEAEEREDGSR
LLENGDKWEQEIAANLR
RVQICMGLLK
TESLYR
TTRYDIDMTK
RPGQSGAWK
WETEIATNLR

**Supp. fig 13:** Complex I of *Polytomella*. Peptides identified by coupled LC-ESI-MS/MS mass spectrometry. Subunit names are blue. For details see Materials and Methods.

| | |
|---|---|
| **15 kDa** | SGFGLQSGTAR, APMITLRK, KEPMIALR (C.r.) |
| **ASHI** | QYPSEVREFLALFKNGFVLNDYSQPDYEEMKRRQSGLLTPIY |
| **B14** | CLPFIHRLHKLEEITSLKEMR, FRVNSPVTDSR, KSSNSAFLDSFYEKAYPLVHSK |
| **B14.5b** | LLDEVQDTYYLEHLKR, GLQVTEEHKKLFSAY |
| **B14.7** | AEEVTVDFSKEPSASSLK, TPIYGKRTYNIIKKLAPIVINKTAYATLLGAIFCATDALYENYSGK, SYTSGMVAGAATGAVFAIGR, GYGTPYGFENLDDAVPTRNPIR |
| **B15** | YTPNIFSPETPMDRSFSHVPK, NYYFSDSKSNFVIPDEREFY |
| **B16.6** | GFPGLK, SIKDLPKVQDVPPPGGFPSIR, LEAEEVYTARSILHPILQAEWDLR, NVPGWK, WLPPVESFDVRPLI |
| **B18** | YGHPLTWRINCAHELHHYNKCSYIEFKRRVAIAIEEKK |
| **B22** | MASEAVLRLPHK, KEIVNWSYGRHEFYPRANALRMEFEANKELSDREQIRKLVLHGEDLLARFRHWEPTIRSEFVGGTVYGVWPQHCK, KFVPNPAYIDEFKQGDTVLLY |
| **ESSS** | AGGASGHPNGSFWSEGTQVGLNGFKYGEIPLPNGQPR, AKDEYAVDMK, LSIVLKDLNMIEEEAYDLIMMR, VLLGLHSGRVPADLKAIYEELEA |
| **KFYI** | SHYCPPDFHYSR, GWIPSQLWNDNVPTPVDYR |
| **MNLL** | PINPYDVPSSPAYHRLQEPSYPVIDK, LMGFKENSR, NVPAALVPEHY |
| **MWFE** | EGKPSWNQNKDQFQR |
| **ND1** | AVMGSLQR, MGPAVSGAFGILQPFWDGFK, YAFLGCLR, DGNGFPCLNLTETASQTK, YDMFMQIGWK |
| **ND2** | ITANENQR, HVAFVFSTLPK, LMYLVSPVK |
| **ND3** | YFVEGEHVVQAEANR, MGALEWRE |
| **ND4** | LVFYLR, LKGGVSSQCVDASRLDVISWNPLVLATVVTGIGYM |
| **ND4L** | AANEVEALNLLK |
| **ND5** | SLIVNK, ATLFLSAGLSIAKENNNFLNR, QGSPTLSFATTIASLNLLGFPELGGFYSK, VLAQLYLFPYGNGR, IFTGSSLSQNMTTNLPAHIK, GGFDVFYAR |
| **ND6** | IQVSTLSTK |
| **NUOP3** | TVFEDTITINVLDYDGK, ALIGTPLNK, YVENVDLKTDDAEAIKLMFKLVVQDYQRETSFFASYLTLNKEMDNMTIGFGPIKPWHITPKWSFNGHHNVK, MFDRLETGPFIE |
| **NUOP4** | ADTSMDHVFGDSTNK |
| **NUOP5** | DSDSRDEYAINAEEPVIYIR, TEVDPTGK, RDPNELQKAQAEVDTLSAAGRLGPLSPF |
| **PDSW** | EVLDIVQSK, NSEWARPEEPILHNLKSEKEEPHVYYHNNNFR, QLIHFEK, IFEDELDKCMR, VASNLERGPNAR, RDVGFIYQNNK, ELGLNNPFPPSSPR |
| **PGIV** | VPYSHELYAMAK, NEKEHNAILECKAQHDWPEDCKPESEAIVR, APEEFKDYAK, EKQAAFEAAFPLSE |
| **SDAP** | EQNLAAGIK |
| **CA1** | MSSLAKTLVWRFGFALKETGLALETLGCKLQGNYSFK, NVSIGHGAVLKGCTIGDNVLVGMNAVISEK, LRDLRPKEIQYLSTLPERYVESSAEHSAMMELLSLK |
| **CA2** | NYPPSYLYPFRHPKPEGVIEKVLFNLGSLFR, YNPAAAPKAGIVAPIPASAQRVLGVKEIVLPSKAESTFIAPNANVLGDVK, SSIWYGAVLR, VESGSIVGAGSIVPPNTVIPAGQVWVGNPAK, NVLPEENGFIASSANNYDLLGQQHKFENSKVFEEMLVEEEIAKDRELLEDKNLAVHQLYIFDPQTQLAAR |
| **CA3** | MASFLKTIVWRCGFAIKETGLALETLGCKLQGNYSFR, HTPLTQYQFKAPSVGESTFVAPSALVSGDVIIGEKSSVLYNAVVR, SVTIGEGSTISDNAYVGSSSEFSPETVIGSNVSVGSGAVLKGCTVGNNVLIGNNVIISEKATVEDNTILAPGSYVPEDVVVKSGELWSGSPAQK, NLDEKELALFNTLSVGATELAADHAVIMKLLELK |

**CAL** NVNFYDAPNGPAVEQVKIEDEWYNRQRTEFNILNKQPSKADDVFIAPNAVVCGDVDLYQK, VSIFFGAVVRGDLNTIRIGENSTILDK, IGDNCIVGARSVVCEGAIMHPESVLAPGSVLAPGRQIPSGELWAGSPAK, ESAEHYYDIATGFRNETFDHGSAWRDAEAYRQKLVDYKEYNWVNYR, DAAELEKLL

**13 kDa** NKIPETVGGAAFK, GTSAVGLGDGLYSHTSK, SPMDYINEVEPIK, GTSYENPAVCK

**18 kDa** IFCPAKAASQSGLGR, APQWKIVFENLSK, WENPLMGWTSTADPLENVGRSTLLFYTK, HGWEYVVDEPNPR

**24 kDa** KIMSNYPSNYKASAMIPLLDLAQQQNGGVVSLAVMNRVAQILEVPPIKVYEVATFFTMFNR, YHVCICGTTPCR, RLQGAQKIEEAITK, NGVEGFSYNYYEDLTPQDAVNILEK, QTAEPAGAVVGDKWIPSSGEQTLMGELPGPYCR

**39 kDa** RGNQVVCPYR, SNVVINCVGIR, NYSYKDVHVDFPTR, IVAESGKVERFIQVSEMGADVSHASR, AVGDEAIMKYIPDATIIR, VQPTYILDVADAVVK, TYYLGGPETLTMRQIYDHLIDTLRLSNDDTVNLRYELAKMLYKPLDTLR, DLSISPAKVTEGLAIEGVR, VGGYDWGDMNKVSSNIPEHIRKYYNLK

**51 kDa** TTFGGLRDQDRIFTNIYGR, GDWYMTK, GRDWIIDQIKK, GRGGAGFASGLKWSFMPK, ASDGRPSYLVVNGDESEPGTCKDREIMRHEPHKLVEGCLVAGTAMGARAGYIYIR, GEFVNER, AVAEAYAKGYLGK, LKPPFPAGMGLYGCPTTVTNVETVAVSPTILRRGPEWFSSFGRK, KLFAISGHVNRPVTVEEEMSIPLRELIER, RGGWDNLLAIIPGGSSVPLLPKKMCDDVIMDFDALRTAQSGLGTAAVIVMNKDTDVIDAIARLSYFYK, REGTGWLYDIMSR, LEEIDMLWEITKQIEGHTICALGDAAAWPVQGLIR, SEMEDRIK

**75 kDa** MEVFVNGEAVVVPKNFTVLQACDAAGVDVPR, FCYHQRLSIAGNCR, APKPVASCAFPAGPGMKIKTDTPVIKK, GRDSEIGTYVEK (C.r.), LNEAVNEEWLSDKGR (C.r.), FIEYKRAVADKNLGPLIK, FTHEVAGTSELGITGRGRDSEVGTYIEKLHSSELSGNVIDLCPVGALLSKPYAFTARSWELKGTETIDVSDALGSNIK, LNDAINEEWLSDKGRFQYDGLKRQRLNTPLVK, GLENATWSAAFDAIRTAIAGAKGNELK, LADAESMIALKDLFNKLGSGNLIHEDGSATLSADVRSSYIANTTIASIEKADVILLVGTNPRFESPVFNAR, KVFLDGAKVGLVGEKVDLTYAYQHLGADVAALESLASGKGAFFEALK, NPVVIVGSSVLR, DDREAVLKTVNDLVDAAGVVKEGWNGFNVLHDNASRVAALDIGFVPSASAR, VVYLLGSDDFKVR, LAEVAPHFAEIGKAESALWLNGQYFKGVKDLVAK, STASLATNISNYYMTDAISR

**B8** EFVLGQYAELKAANPLLPILVRESNAAQASLTAR, KVSIENDSASGILSKLEGLVK

**B13** AVADVEIDFSNR, KYVGVDHLSKEQGTKAKFIVSLNSLLEAVKSVPETAEYRKAIEATCQYR, VCGENNSDAAIEEVLDAHLEELIAESK, KYVGVDHLYKEQATK, FIMSINSLLEAVK, KAIEATCQYR (blue: isoform)

**B14.5a** MAEALK (C.r.), TVQSIFYSVGLKEPWK, SLPDFEYYLPFGLTYR, AVVPHDVPK, TPFDYSK, LYAGMPTK (C.r.), VFGSAPLKPADVK, IPEVMPHR

**B17.2** YLNNLVSTSGKKSLAGFFVSWEFAK, HNNVVRPQSGAQVGIDK, WIVFADKFDYK, GSWFNPAK, KVEIWQPPQV

**ND7** VSNFTINFGPQHPAAHGVLRLVLEMDGEIIKR, LLEYKTYNQGIPYFDR, VLFSEITR, EKLFEFYER, MHAAYFRVGGVAQDLPIGLLRDIYDWSR, VDEMEELLTGNR, TIDVGLVTAQQAWDWGCSGPILRGSGIDWDLRKNQPYDVYGRMDFNVPIAGHGDCYDR, IIYQCLNEMPDGLYK, QSMESLIHHFKLFSEGYHVPAGETYR, GEFGVYLVSR, SPGYAHLQMLDMVAKGAMLADVVTIIGTLDVVFGEIDR

**ND9** NSQALLGR, MTDKDYIAYDNIKNFGDNYLTDYIIK, DHVNLQYKTLIDITAVDYPER, FEVVYHLLSPR, IWNAANWFER, VLTDYGFTGHPLRK, YDYGK, VISEPLELTQEFRYFDFSSPWDTLSR

**PSST** GADFIVSKVDTVVNWARAGSMWPMTFGLACCAVEMMHAGASRYDLDRFGIIFRPSPRQSDVMIVAGTLTNKMAPALRK-VYDQMPEPK, VVPVDVYVPGCPPTAEGLLYGLLQLQK, SKNTQLWWNK

**TYKY** RPGQSGAWK, YSSEWEQDPTFK, GMMYSASGFFDDK, FRGEHALRRYPTGEER, AQAITIEAEEREDGSR, YDIDMTK, REELLYDKQKLLENGDKWEQEIAANLRTESLYR

**Supp. fig. 14:** Partially assembled peptides of complex I subunits from *Polytomella sp.* Assignments are based on the *Chlamydomonas reinhardii* genome sequence (Merchant et al. 2007) and a partial genome sequence of *Polytomella* sp. (Murphy et al. 2019). Peptides labeled (*C.r.*) exactly match sequences in the *Chlamydomonas* genome that were not covered by the *Polytomella* partial genome sequence.

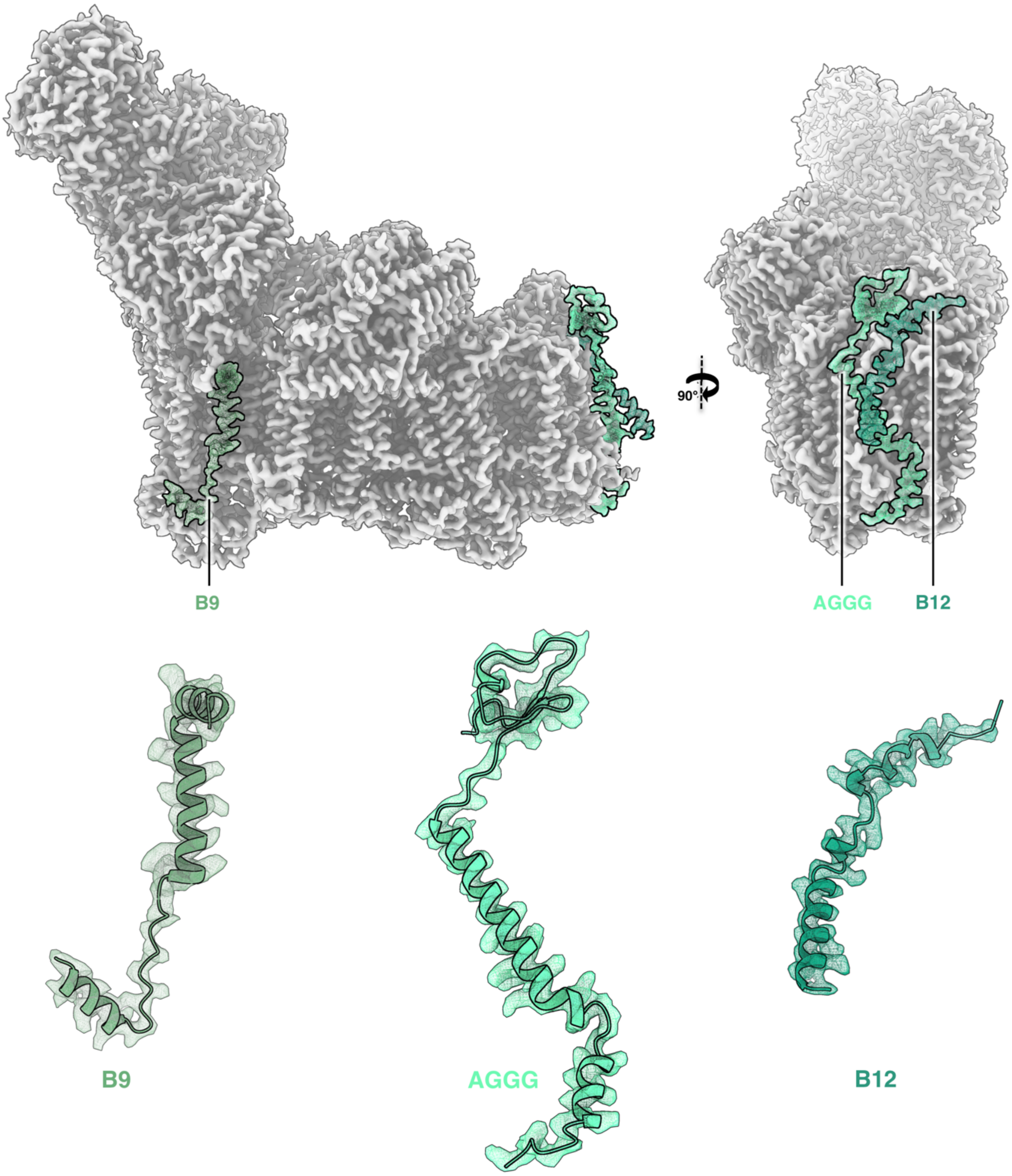

**Supp. fig. 15:** Subunits of *Polytomella* complex I assigned by structural homology. Accessory subunits B9, AGGG and B12 of *Polytomella* complex I were assigned by their position within the *Arabidopsis* or bovine complex I.

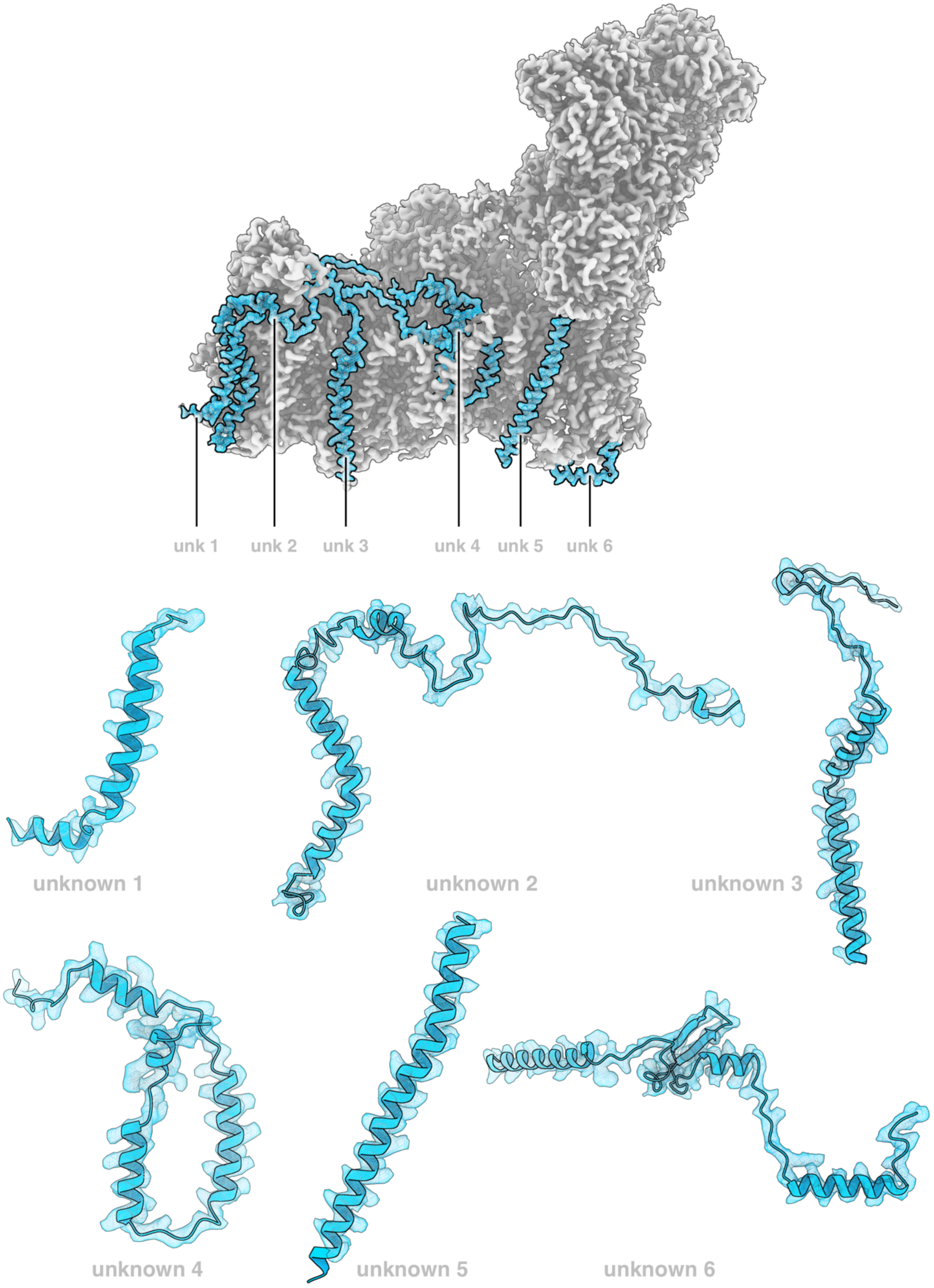

**Supp. fig. 16:** *Polytomella* complex I. Non-assigned densities of 6 unknown subunits in the membrane arm.

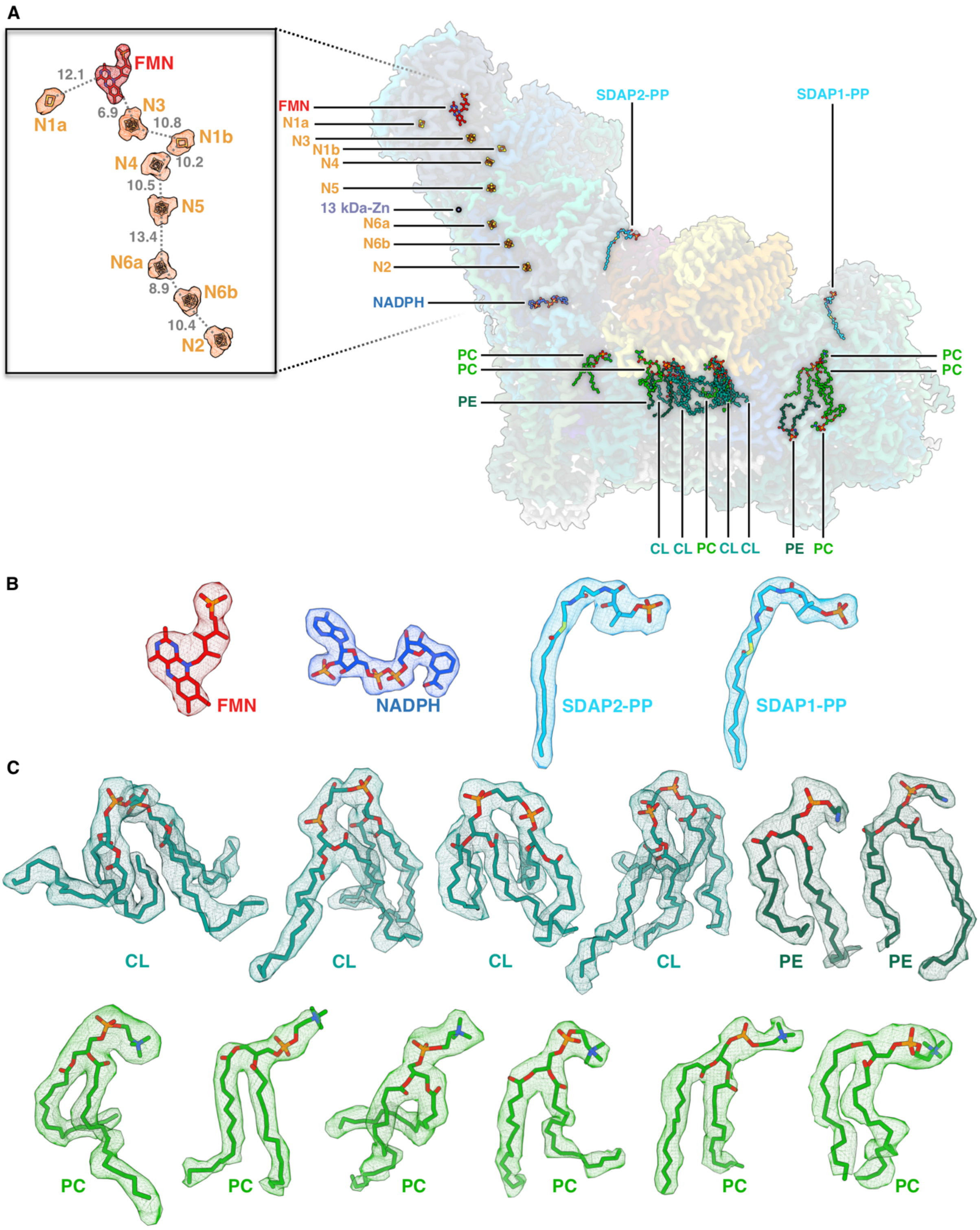

**Supp. fig. 17:** Cofactors and lipids of *Polytomella* complex I. FMN: Flavin mononucleotide; N1a, N1b, N2, N3, N4, N5, N6a, N6b: FeS clusters in the peripheral arm; NADPH: Nicotinamide adenine dinucleotide phosphate; SDAP1-PP, SDAP2-PP: 4'-phosphopantetheine bound to SDAP1 and SDAP2; CL: cardiolipin; PC: phosphatidylcholine; PE, phosphatidylethanolamine.

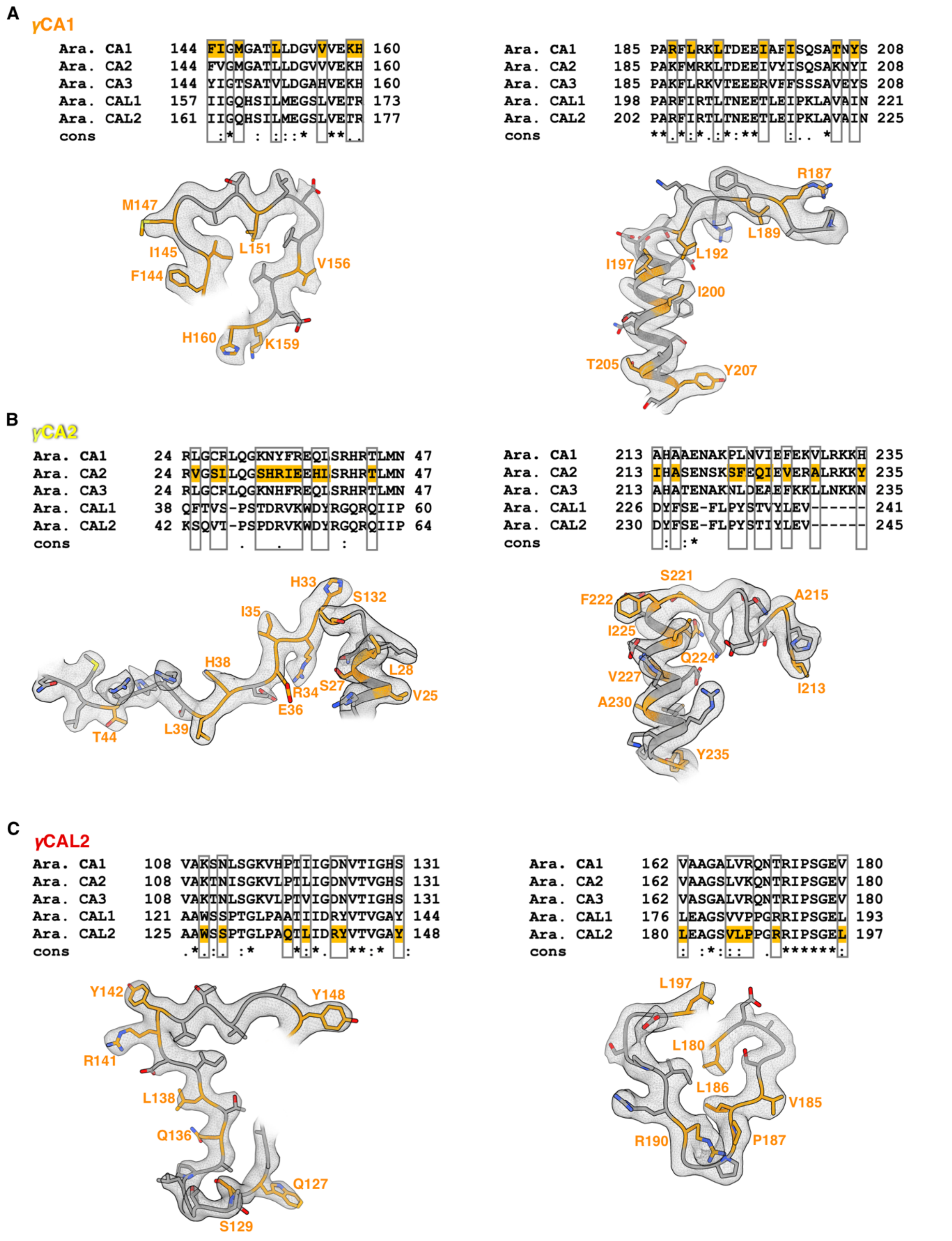

**Supp. fig. 18:** γCA/γCAL proteins of the heterotrimeric γCA domain in *Arabidopsis*. Sequence regions differing between the five Arabidopsis γCA/γCAL proteins are aligned and used to interpret the cryo-EM density. **A:** Evidence that γCA1 is localized at one defined position within the heterotrimer. **B:** Evidence that γCA2 is localized at the second position within the heterotrimer. **C:** Evidence that γCAL2 is localized at the third position. The numbers indicate the first and the last amino acid position of the compared protein sequences, respectively. Amino acids used to identify the different proteins are indicated in orange. Maps are drawn in ChimeraX at a density threshold level of 0.007 for γCA1, or 0.008 for γCA2 and γCAL2.

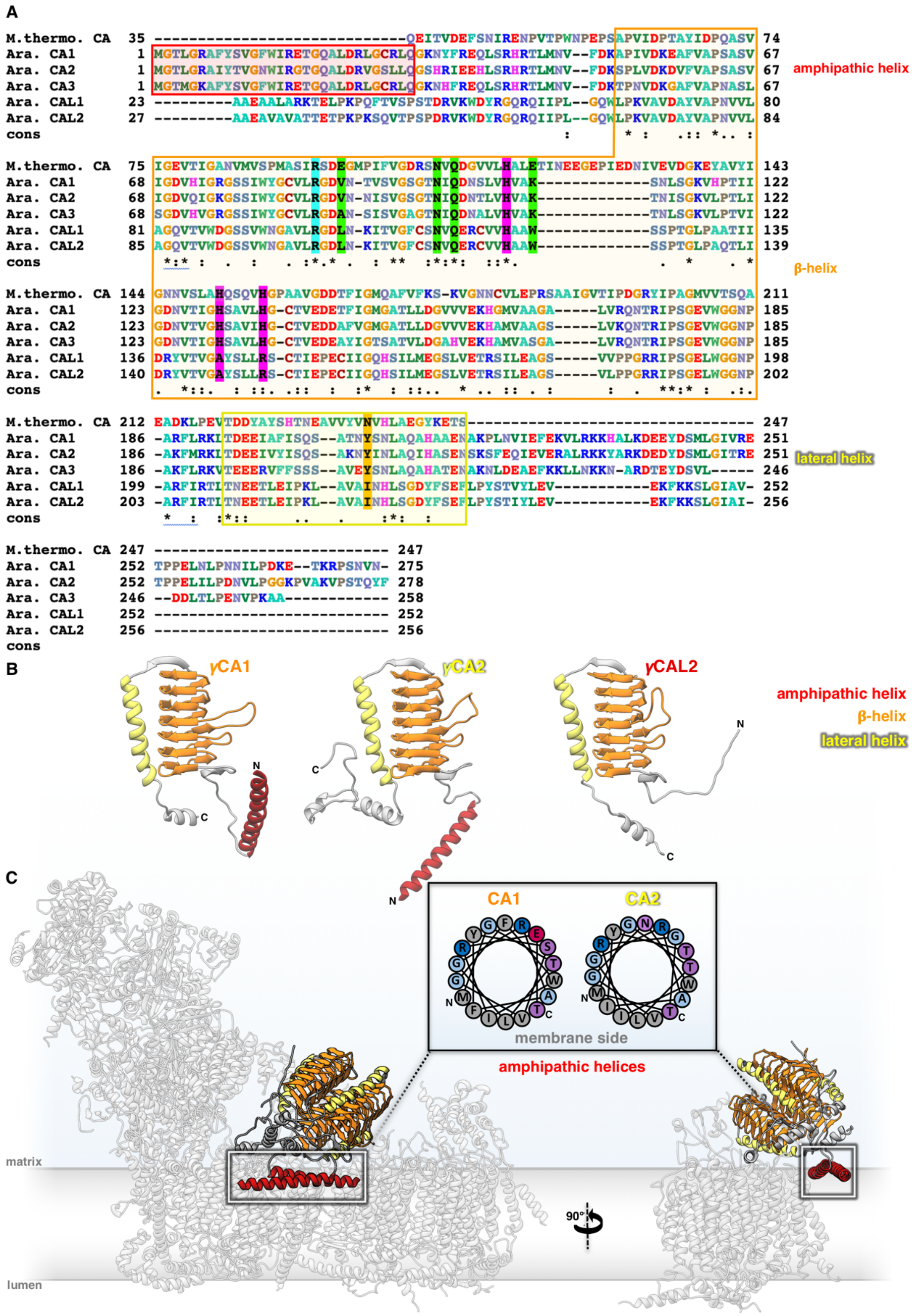

**Supp. fig. 19:** Structures of γCA and γCAL subunits in *Arabidopsis* complex I. **A:** Alignment of the five γCA/γCAL proteins to the prototypic γCA from *Methanosarcina thermophila*. The three histidines coordinating the zinc ion in the *M. thermophila* active site are highlighted in magenta. Note that two of the three histidines are not conserved in γCAL1 and γCAL2. Catalytically important residues are highlighted in green and orange. Residues that stabilize the active site are highlighted in cyan. Secondary structure elements are boxed in color as indicated. **B:** Structures of γCA1, γCA2 and γCAL2 subunits in the cryo-EM map. γCA1 and γCA2 have an N-terminal amphipathic helix (red). **C:** The two amphipathic helices form a coiled coil, which anchors the γCA domain to the membrane. Helix wheel projections (inset) indicate that hydrophobic residues orient towards the membrane.

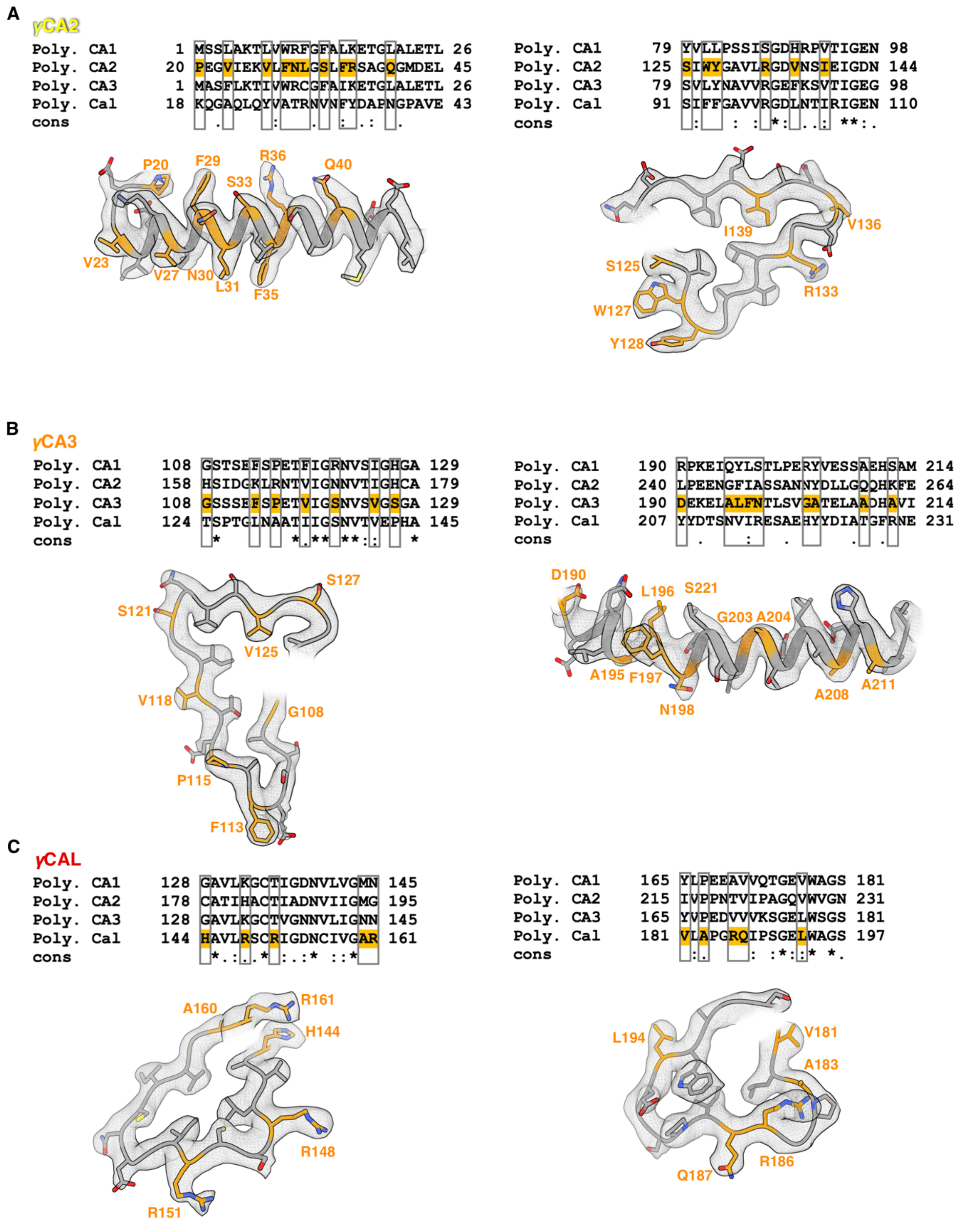

**Supp. fig. 20:** γCA/γCAL subunits in the heterotrimeric γCA domain of *Polytomella*. The γCA/γCAL subunits in the cryo-EM density were identified by their different sequences. **A:** One defined position within the heterotrimer is occupied by γCA2. **B:** A second position is occupied by γCA3. **C:** The third position is occupied by γCAL. Numbers indicate the first and the last residues of the compared protein sequences. Amino acids used to specify the different proteins are indicated in orange. All maps are drawn in ChimeraX at a density threshold level of 0.01.

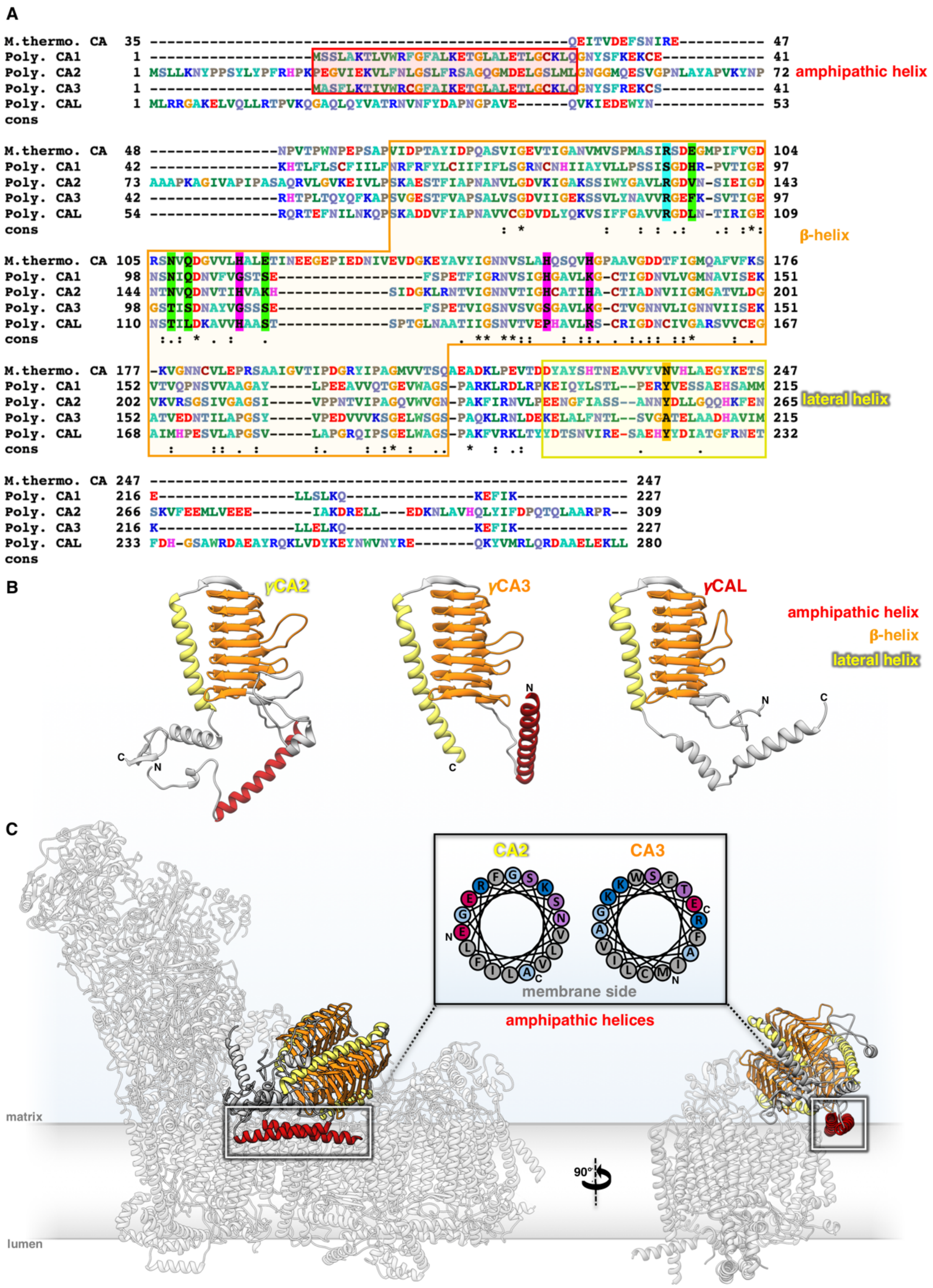

**Supp. fig. 21:** Structures of γCA and γCAL subunits in *Polytomella* complex I. **A:** Alignment of the four γCA/γCAL proteins to the prototypic γCA from *M. thermophila*. Histidines that coordinate the zinc ion are highlighted in magenta. Note that the three histidines are conserved only in *Polytomella* γCA2. Catalytically important residues are highlighted in green and orange. Residues that stabilize the active site are highlighted in cyan. Secondary structure elements are boxed in color as indicated. **B:** Structures of γCA1, γCA2 and γCAL2 subunits in the cryo-EM map. γCA2 and γCA3 have an N-terminal amphipathic helix (red). **C:** The two amphipathic helices form a coiled coil, which anchors the γCA domain to the membrane. Helix wheel projections (inset) indicate that hydrophobic residues orient towards the membrane.

A

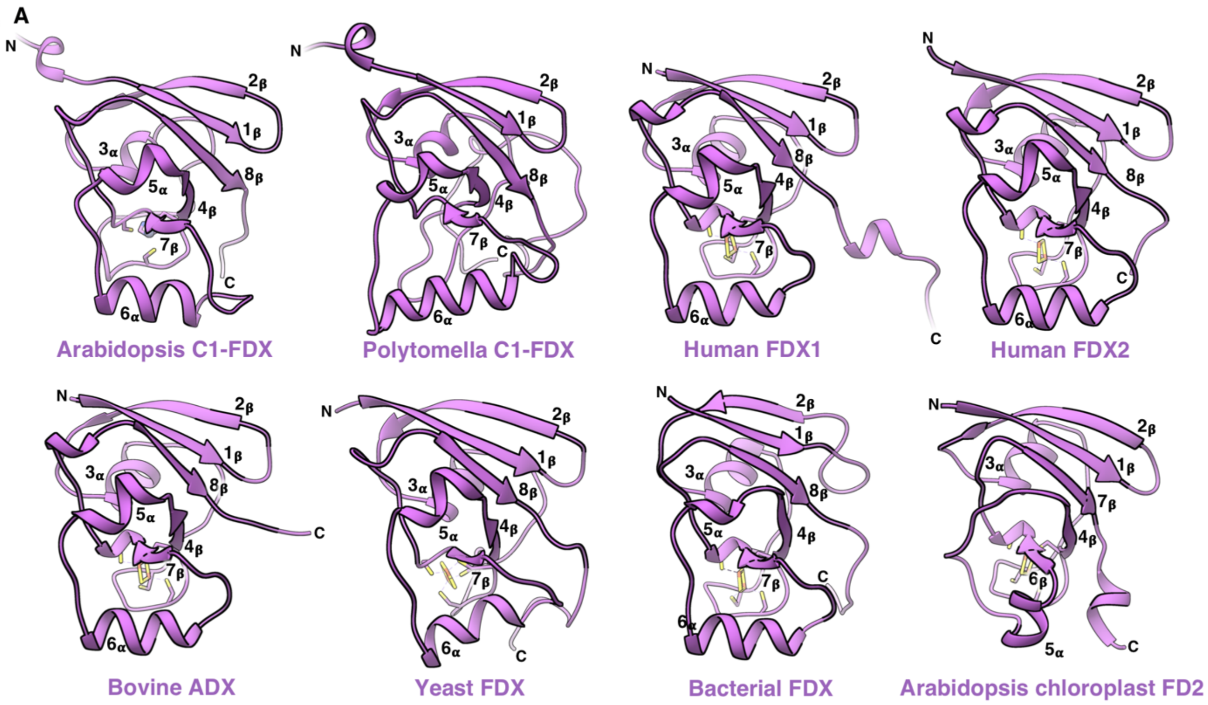

B

```
Ara. C1-FDX          1 -------------MATTLQKLSSQIHRLSPFTRS--------LIVR---------TSATSAPSPSLGSKKVSDRIVKLSAID 52
Poly. C1-FDX         1 ---------------MALLRALAKPLRSLQAVSSVAQ-----VSLRQFGAA-SHHDDHHDDHDHYTPPKTVFEDTITINVLD 61
H. sapiens FDX1      1 --MAAAGGARLLRAASAVLGGPAGRWLHHAGSRAGSS-----GLLRNRGP--GGSAEASRSLSVSARARSSSEDKITVHFIN 73
H. sapiens FDX2      1 MHVMAASMARGGVSARVLLQAARGTWWNRPGGTSGSGEGVALGTTRKFQA-----TGSRPAGEEDAGGPERPGDVVNVVFVD 77
B. taurus ADX        1 ------MAARLLRVASAALGDTAGRWRLLARPRAGAG-----GLRGSRGPGLGGGAVATRTLSVSGRAQSSSEDKITVHFIN 71
S. cerevisiae FDX    1 ----------MLKIVTRAGHTARISNIAAHLLRTSPS-----LLTRTTTTTRFLPFSTSSFLNHGHLKKPKPGEELKITFIL 67
P. putida FDX        1 -------------------------------------------------------------------MP--LVTFLPHEKFC 13
Ara. chloro. FD2     1 ---------------MASTALSSAIVGTSFIRRSPAP-----ISLRSLPSA-NTQSLFGLKSGTARGGRVTAMATYKVKFIT 61

Ara. C1-FDX         53 PDGYKQDIIGLSGQTLLRALTHTGLIDPA--SHRLDDIEACSAECEVQIAEEWLEKLPPRT--YDEEYVLKRSSRSRILNKH 130
Poly. C1-FDX        62 YDGKKHAVKALIGTPLNKALVEYGFSSTYFFPNMGYYTQHISD-AHVFIPEEYWKYVENVDLKTDDAEAIKLMFKL-VVQDY 141
H. sapiens FDX1     74 RDGETLTTKGKVGDSLLDVVVENNLDIDGFGACEG--TLACST-CHLIFEDHIYEKLDAIT--DEENDMLDLAY---GLTDR 147
H. sapiens FDX2     78 RSGQRIPVSGRVGDNVLHLAQRHGVDLEG--ACEA--SLACST-CHVYVSEDHLDLLPPPE--EREDDMLDMAP---LLQEN 149
B. taurus ADX       72 RDGETLTTKGKIGDSLLDVVVQNNLDIDGFGACEG--TLACST-CHLIFEQHIFEKLEAIT--DEENDMLDLAY---GLTDR 145
S. cerevisiae FDX   68 KDGSQKTYEVCEGETILDIAQGHNLDMEG--ACGG--SCACST-CHVIVDPDYYDALPEPE--DDENDMLDLAY---GLTET 139
P. putida FDX       14 PEG--LTVEVKPGTNILELAHDHHIEMES--ACGG--VKACTT-CHCIVRKG-FDSLEEAD--ELEEDMLDKAW---GLEAQ 82
Ara. chloro. FD2    62 PEGE-LEVECDDDVYVLDAAEEAGIDLPY--SCRAGS---CSS-CAGKVVSGSVDQSDQS---FLDDEQIGEGF-------- 125
                        .*         . :        .                :  .   .     .          :   :

Ara. C1-FDX        131 SRL----GCQVVLTQELQGMVVAVPEAKPWDIP-------------------------- 159
Poly. C1-FDX       142 QRETSFFASYLTLNKEMDNMTIGFGPIKPWHITPKWSFNGHHNVKDRMFDRLETGPFIE 200
H. sapiens FDX1    148 SRL----GCQICLTKSMDNMTVRVPETVADARQSIDVGKTS------------------ 184
H. sapiens FDX2    150 SRL----GCQIVLTPELEGAEFTLPKITRNFYVDGHVPKPH------------------ 186
B. taurus ADX      146 SRL----GCQICLTKAMDNMTVRVPDAVSDARESIDMGMNSSKIE-------------- 186
S. cerevisiae FDX  140 SRL----GCQIKMSKDIDGIRVALPQMTRNVNNNDFS---------------------- 172
P. putida FDX       83 SRL----GCQVFVAD--EDLTIEIPKYSLNHAAEA----PH------------------ 113
Ara. chloro. FD2   126 VLT-----CAAYPTSD-----VTIETHKEEDIV-------------------------- 148
```

**Supp. fig. 22:** *Arabidopsis* mitochondrial C1-FDX. *Arabidopsis* mitochondrial C1-FDX was compared to two human mitochondrial ferredoxins (FDX1, pdb: 3P1M; FDX2, pdb: 2Y5C), a bovine adrenodoxin (ADX, pdb: 1CJE) and a yeast mitochondrial FDX (pdb: 2MJD), a bacterial FDX (pdb: 3AH7) and *Arabidopsis* chloroplast FD2 (pdb: 4ZHO). **A:** Structural comparison, **B:** Sequence comparison. The four cysteines conserved in most sequences bind the FDX Fe2S2 cluster. Note that only three of the four cysteines are conserved in *Arabidopsis* C1-FDX, and none of them in *Polytomella* C1-FDX.

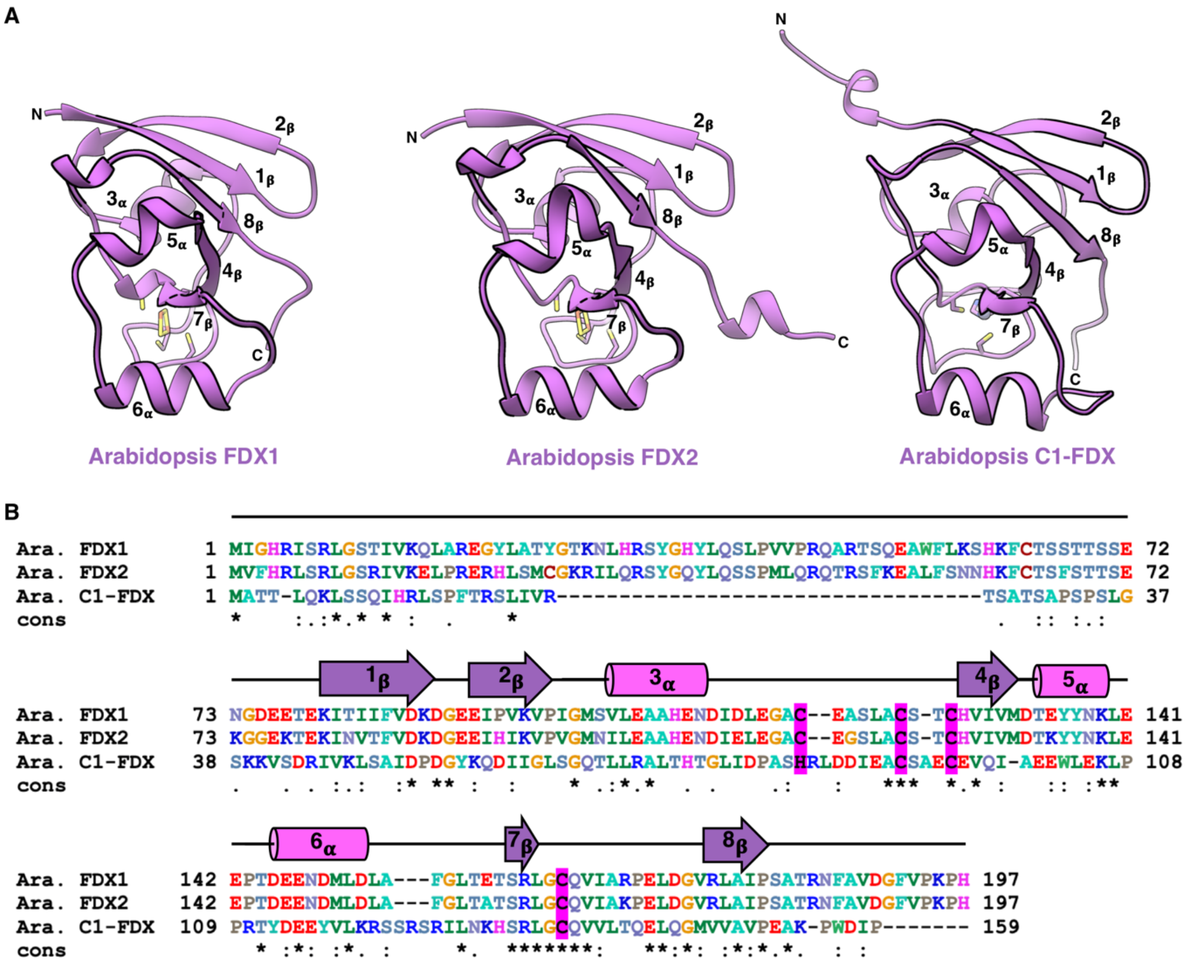

**Supp. fig. 23:** Comparison of *Arabidopsis* C1-FDX to *Arabidopsis* mitochondrial FDX1 and FDX2. **A:** Structural comparison. *Arabidopsis* FDX1 (accession number At4g05450) and FDX2 (At4g21090) were modeled on the X-ray structure of human mitochondrial ferredoxin (pdb: 2Y5C, 3P1M) using the SWISS MODEL server. **B:** Sequence comparison. The four cysteines coordinating the 2Fe2S cluster (magenta) are conserved in FDX1 and FDX2. One of the four cysteines is substituted by a histidine in C1-FDX.

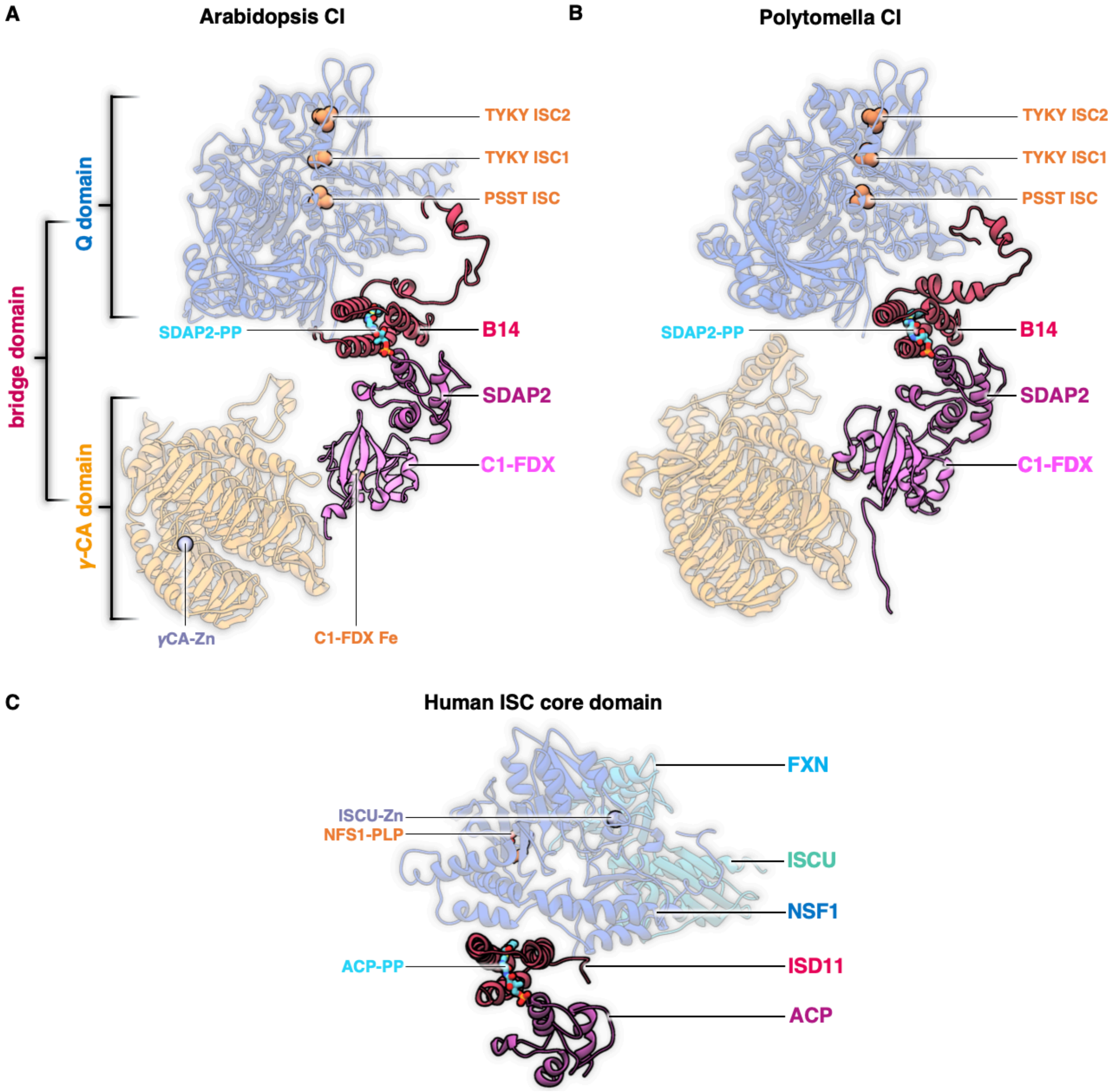

**Supp. fig. 24:** Comparison of the bridge domain of *Arabidopsis* and *Polytomella* complex I to the human iron sulfur cluster (ISC) assembly core complex (Fox et al., 2019). **A:** Bridge domain in *Arabidopsis* complex I located between the Q domain of the peripheral arm and the γCA domain of the membrane arm. The bridge consists of subunits B14, SDAP2 and C1-FDX. Iron sulfur clusters of the Q domain and the position of the metal ion (presumably Fe) in the γCA domain are orange. 4'-phosphopantetheine (SDAP2-PP) bound to SDAP2 is light blue. Zinc ion (γCA-Zn) bound in the active site of the γCA domain is purple. **B:** In *Polytomella* complex I, the bridge domain is located between the Q domain and the γCA domain. **C:** Core domain of the human iron sulfur cluster (ISC) machinery. ISCU, scaffold protein of the human ISC assembly core complex; NSF1, mitochondrial cysteine desulfurase; ISD11, LYR family protein; ACP, acyl carrier protein; FXN: frataxin; ISCU Zn, zinc bound to the ISCU protein; NFS1-PLP: pyridoxal phosphate bound to NSF1; ACP-PP, 4'-phosphopantetheine bound to ACP. Note that subunits B14 and SDAP2 of the *Arabidopsis* and *Polytomella* bridge domain are homologous to ISD11 and ACP in the human ISC core domain, respectively, and that there is as yet no structure for the mitochondrial ferredoxin that is known to form part of the human ISC assembly core complex.

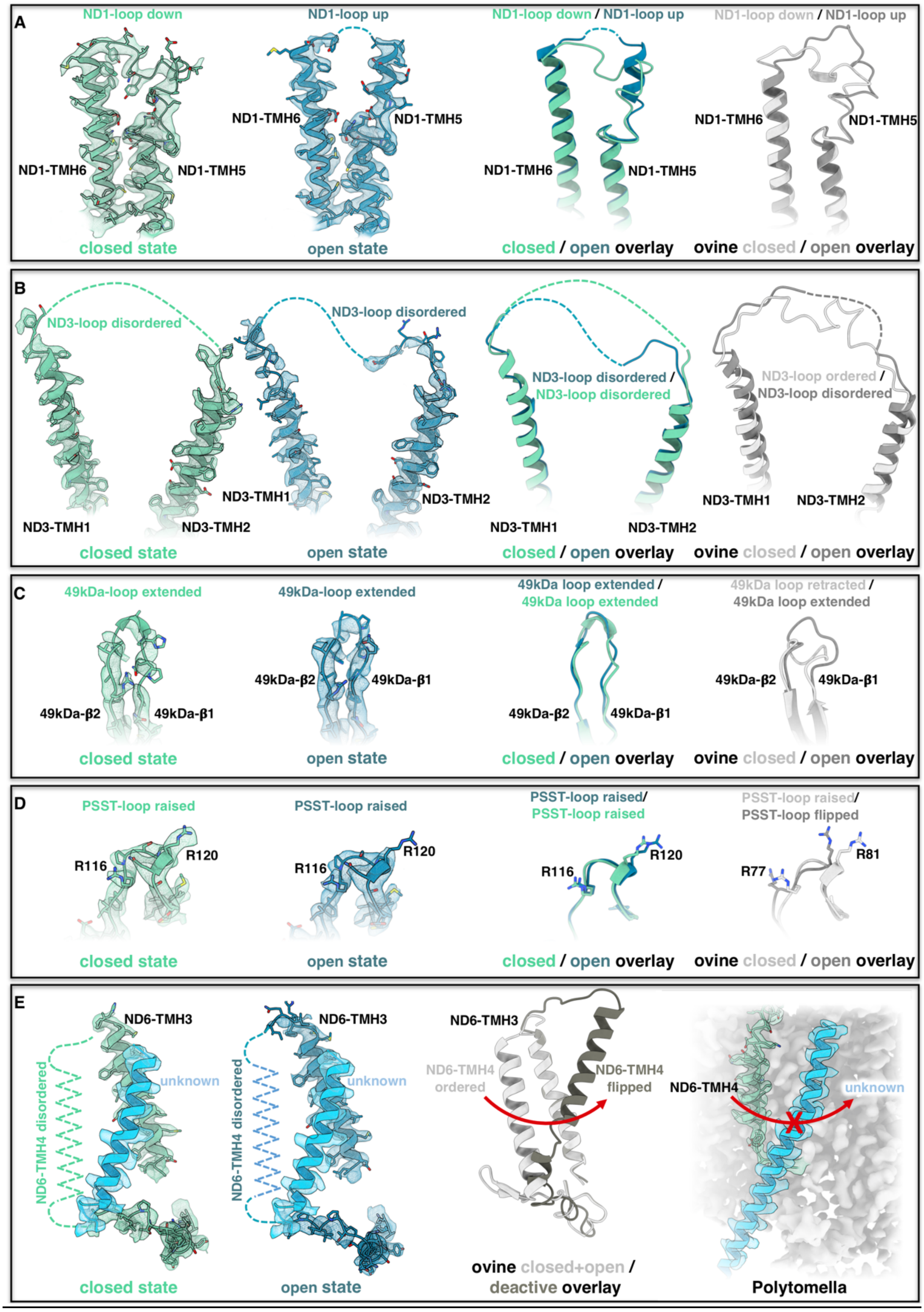

**Supp. fig. 25:** Loop conformations in the closed and open state of *Arabidopsis* complex I. Loops in the closed (green) and open (blue) state are shown with their map densities. For comparison, the loop structures are shown side by side to the corresponding loops in ovine complex I (light and dark grey) (Kampjut and Sazanov, 2020). **A:** ND1 loop between TMH5 and 6 (see also Figure 5D). **B:** Loop connecting TMH1 and 2 of ND3. **C:** 49kDa subunit loop between β-sheet 1 and 2. **D:** PSST loop with conserved arginine residues. **E:** ND6 TMH4. In the deactive state of ovine complex I, ND6 TMH4 flips to a position that is occupied by a non- assigned helix density in the membrane arm of *Arabidopsis* and *Polytomella* (light blue).

**Supplementary Tables:** (following pages)

#### Supp. table 1: Nomenclature of complex I subunits in *A. thaliana*, *C. reinhardtii* and other model species

| *A. thaliana* | accession | *C. reinhardtii* | accession | *B. taurus* | *H. sapiens* | *Yarrowia* | *E. coli* | *T. therm.* |
|---|---|---|---|---|---|---|---|---|
| **► *Membrane arm (incl. bridge domain)*** | | | | | | | | |
| 15 kDa-1<br>15 kDa-2 | At3g62790<br>At2g47690 | 15 kDa | Cre12.g511200 | 15kDa | NDUFS5 | NIPM | - | - |
| | | | | 42 kDa | NDUFA10 | - | | |
| AGGG | At1g76200 | AGGG | n.d. | AGGG | NDUFB2 | NIGM | - | - |
| ASHI | At5g47570 | ASHI | Cre01.g007850 | ASHI | NDUFB8 | NIAM | - | - |
| B9 | At2g46540 | B9 | Cre12.g537050 | B9 | NDUFA3 | NI9M | - | - |
| B12-1<br>B12-2 | At1g14450<br>At2g02510 | B12 | Cre05.g244850 | B12 | NDUFB3 | NB2M | - | - |
| B14 | At3g12260 | B14 | Cre12.g555250 | B14 | NDUFA6 | NB4M | - | - |
| B14.5b | At4g20150 | B14.5b | Cre13.g571150 | B14.5b | NDUFC2 | N4BM | - | - |
| B14.7 | At2g42210 | B14.7 | Cre14.g617826 | B14.7 | NDUFA11 | NUJM | - | - |
| B15 | At2g31490 | B15 | Cre03.g204650 | B15 | NDUFB4 | NB5M | - | - |
| B16.6-1<br>B16.6-2 | At1g04630<br>At2g33220 | B16.6 | Cre16.g664600 | B16.6 | GRIM19 | NB6M | - | - |
| | | | | B17 | NDUFB6 | - | | |
| B18 | At2g02050 | B18 | Cre06.g278188 | B18 | NDUFB7 | NB8M | - | - |
| B22 | At4g34700 | B22 | Cre11.g467668 | B22 | NDUFB9 | NI2M | - | - |
| ESSS-1<br>ESSS-2 | At2g42310<br>At3g57785 | ESSS | Cre05.g240800 | ESSS | NDUFB11 | NESM | - | - |
| KFYI | At4g00585 | KFYI | Cre17.g725400 | KFYI | NDUFC1 | - | - | - |
| MNLL | At4g16450 | MNLL | Cre06.g267200 | MNLL | NDUFB1 | - | - | - |
| MWFE | At3g08610 | MWFE | Cre10.g459750 | MWFE | NDUFA1 | NIMM | - | - |
| ND1 | AtMg00516<br>AtMg01120<br>AtMg01275 | ND1 | AAB93446 | ND1 | ND1 | NU1M | NUOH | NQO8 |
| ND2 | AtMg00285<br>AtMg01320 | ND2 | AAB93444 | ND2 | ND2 | NU2M | NUON | NQO14 |
| ND3 | AtMg00990 | ND3 | Cre08.g378900 | ND3 | ND3 | NU3M | NUOA | NQO7 |
| ND4 | AtMg00580 | ND4 | AAB93441 | ND4 | ND4 | NU4M | NUOM | NQO13 |
| ND4L | AtMg00650 | ND4L | Cre09.g402552 | ND4L | ND4L | NULM | NUOK | NQO11 |
| ND5 | AtMg00060<br>AtMg00513<br>AtMg00665 | ND5 | AAB93442 | ND5 | ND5 | NU5M | NUOL | NQO12 |
| ND6 | AtMg00270 | ND6 | AAB93445 | ND6 | ND6 | NU6M | NUOJ | NQO10 |
| PDSW-1<br>PDSW-2 | At3g18410<br>At1g49140 | PDSW | Cre12.g555150 | PDSW | NDUFB10 | NIDM | - | - |
| PGIV-1<br>PGIV-2 | At3g06310<br>At5g18800 | PGIV | Cre07.g333900 | PGIV | NDUFA8 | NUPM | - | - |
| SGDH | At1g67785 | SGDH | Cre03.g178250 | SGDH | NDUFB5 | NUNM | - | - |
| SDAP-1<br>SDAP-2<br>SDAP-3 | At1g65290<br>At2g44620<br>At5g47630 | SDAP | Cre16.g673109 | SDAP-α<br>SDAP-β | NDUFAB1<br>NDUFAB1 | ACPM1<br>ACPM2 | | |
| P1 | At1g67350 | - | | - | - | - | - | - |
| P2 | At2g27730 | - | | - | - | - | - | - |
| Ferredoxin | At3g07480 | NUOP3 | Cre02.g100200 | | | | | |
| | | NUOP4 | Cre08.g378550 | | | | | |
| | | NUOP5 | Cre08.g378050 | | | | | |
| | | | | | | NUXM | | |
| | | | | | | NUUM | | |
| | | | | | | ST1 | | |
| **► *Carbonic anhydrase domain*** | | | | | | | | |
| CA2 | At1g47260 | CA2 | Cre09.g415850 | - | - | - | - | - |
| CA1 | At1g19580 | CA1/CA3 | Cre12.g516450 | - | - | - | - | - |
| CA3 | At5g66510 | | | - | - | - | - | - |
| CAL1 | At5g63510 | CAL | Cre06.g293850 | - | - | - | - | - |
| CAL2 | At3g48680 | | | - | - | - | - | - |
| **► *Peripheral arm*** | | | | | | | | |
| | | 10 kDa | Cre03.g178250 | 10 kDa | NDUFV3 | - | | |
| 13 kDa | At3g03070 | 13 kDa | Cre03.g146247 | 13 kDa | NDUFS6 | NUMM | - | - |
| 18 kDa | At5g67590 | 18 kDa | Cre10.g450400 | 18 kDa | NDUFS4 | NUYM | - | - |
| 24 kDa | At4g02580 | 24 kDa | Cre10.g434450 | 24 kDa | NDUFV2 | NUHM | NUOE | NQO2 |
| 39 kDa | At2g20360 | 39 kDa | Cre10.g422600 | 39 kDa | NDUFA9 | NUEM | - | - |
| 51 kDa | At5g08530 | 51 kDa | Cre12.g535950 | 51 kDa | NDUFV1 | NUBM | NUOF | NQO1 |
| 75 kDa | At5g37510 | 75 kDa | Cre16.g679500 | 75kDa | NDUFS10 | NUAM | NUOG | NQO3 |
| B8 | At5g47890 | B8 | Cre13.g568800 | B8 | NDUFA2 | NI8M | - | - |
| B13 | At5g52840 | B13 | Cre12.g484700 | B13 | NDUFA5 | NUFM | - | - |
| B14.5a | At5g08060 | B14.5a | Cre11.g467767 | B14.5a | NDUFA7 | NUZM | - | - |
| B17.2 | At3g03100 | B17.2 | Cre09.g405850 | B17.2 | NDUFA12 | N7BM | - | - |
| ND7 | AtMg00510 | ND7 | Cre07.g327400 | 49 kDa | NDUFS2 | NUCM | NUOD | NQO3 |
| ND9 | AtMg00070 | ND9 | Cre12.g492300 | 30 kDa | NDUFS3 | NUGM | NUOC | NQO5 |
| PSST | At5g11770 | PSST | - | PSST | NDUFS7 | NUKM | NUOB | NQO6 |
| TYKY-1<br>TYKY-2 | At1g79010<br>At1g16700 | TYKY | Cre12.g496750 | TYKY | NDUFS8 | NUIM | NUOI | NQO9 |
| | | | | | | | | NQO15 |

Subunits specific for clades are highlighted by colors.

**Supplementary table 2. EM Statistics *Arabidopsis thaliana* complex I**

| | |
|---|---|
| **Data Collection Arabidopsis** | |
| Electron Microscope | Titan Krios G3i |
| Camera | Gatan K3 (electron-counting mode) |
| Data collection software | EPU (Thermo Fisher) |
| Voltage | 300 kV |
| Nominal magnification | 105,000 x |
| Calibrated physical pixel size | 0.837 Å |
| Total exposure | 64 $e^-$Å$^{-2}$ / 43 $e^-$Å$^{-2}$ |
| Exposure rate | 15 $e^-$pixel$^{-1}$s$^{-1}$ |
| Number of frames | 50 |
| Defocus range | −0.5 to −2 µm |
| **Image Processing Arabidopsis** | |
| Motion correction software | MotionCor2 |
| CTF estimation software | CTFFind4.1.13 |
| Particle selection software | crYOLO |
| Micrographs used | 12,768 |
| Particles selected | 1,874,295 |
| Cassification and refinement software | Relion3 |
| Particles contributing to final dataset | 459,177 |
| **Model Building Arabidopsis** | |
| Modeling software | Coot |
| Refinement software | Phenix (phenix.real_space_refine) |

**Supplementary table 3. EM Statistics *Polytomella sp.* complex I**

| | |
|---|---|
| **Data Collection Polytomella** | |
| Electron Microscope | Titan Krios G3i |
| Camera | Gatan K3 (electron-counting mode) |
| Data collection software | EPU (Thermo Fisher) |
| Voltage | 300 kV |
| Nominal magnification | 105,000 x |
| Calibrated physical pixel size | 0.837 Å |
| Total exposure | 64 $e^{-}Å^{-2}$ |
| Exposure rate | 15 $e^{-}pixel^{-1}s^{-1}$ |
| Number of frames | 50 |
| Defocus range | −0.7 to −2.5 µm |
| **Image Processing Polytomella** | |
| Motion correction software | MotionCor2 |
| CTF estimation software | CTFFind4.1.13 |
| Particle selection software | crYOLO |
| Micrographs used | 1,652 |
| Particles selected | 195,537 |
| Cassification and refinement software | Relion3 |
| Particles contributing to final dataset | 42,350 |
| **Model Building Polytomella** | |
| Modeling software | Coot |
| Refinement software | Phenix (phenix.real_space_refine) |

**Supp. table 4: *Arabidopsis thaliana* complex I map identifiers and statistics**

| Description | PDB ID | EMDB ID | Resolution (0.143) | Applied B-factor | # of particles | Symm. | Details |
|---|---|---|---|---|---|---|---|
| Full complex | 7ARB | 11878 | 3.41 Å | -80 | 459,177 | C1 | Generated from 3D classification and 3D refinement by applying a softmask around the entire complex |
| Peripheral arm | 7AQR | 11873 | 3.21 Å | -96 | 459,177 | C1 | Generated by 3D multi-body refinement by applying a softmask around the peripheral arm, the membrane arm core and the membrane arm tip |
| Membrane arm core | 7AQQ | 11872 | 3.39 Å | -92 | 459,177 | C1 | Generated by 3D multi-body refinement by applying a softmask around the peripheral arm, the membrane arm core and the membrane arm tip |
| Membrane arm tip | 7AQW | 11874 | 3.43 Å | -101 | 459,177 | C1 | Generated by 3D multi-body refinement by applying a softmask around the peripheral arm, the membrane arm core and the membrane arm tip |
| Closed state | 7AR8 | 11876 | 3.77 Å | -93 | 42,096 | C1 | Generated by focused 3D Classification and 3D refinement by applying a softmask around the membrane arm and the entire complex |
| Open state | 7AR7 | 11875 | 3.72 Å | -95 | 48,933 | C1 | Generated by focused 3D Classification and 3D refinement by applying a softmask around the membrane arm and the entire complex |

**Supp. table 5: *Polytomella* sp. complex I map identifiers and statistics**

| Description | PDB ID | EMDB ID | Resolution (0.143) | Applied B-factor | # of particles | Symm. | Details |
|---|---|---|---|---|---|---|---|
| Full complex | 7ARD | 11880 | 3.53 Å | -83 | 42,350 | C1 | Generated from 3D classification and 3D refinement |
| Peripheral arm | 7ARC | 11879 | 3.30 Å | -69 | 42,350 | C1 | Generated by 3D multi-body refinement by applying a softmask around the peripheral and membrane arm |
| Membrane arm | 7AR9 | 11877 | 3.34 Å | -73 | 42,350 | C1 | Generated by 3D multi-body refinement by applying a softmask around the peripheral and membrane arm |

**Supp. table 6: *Arabidopsis thaliana* complex I model identifiers and quality statistics**

| Description | PDB ID | EMDB ID | Residues built | RMS Bonds Length/Angles | | Ramachandran Outliers (%) | Ramachandran favoured (%) | Rotamer outliers (%) | Clashscore | EMRinger score |
|---|---|---|---|---|---|---|---|---|---|---|
| Full complex | 7ARB | 11878 | 7,789 | 0.006 | 0.792 | 0.13 | 90.32 | 0.03 | 14.55 | 2.22 |
| Peripheral arm | 7AQR | 11873 | 3,271 | 0.014 | 0.986 | 0.25 | 87.67 | 0.00 | 13.03 | 3.55 |
| Membrane arm core | 7AQQ | 11872 | 2,938 | 0.005 | 0.728 | 0.07 | 90.48 | 0.00 | 11.43 | 2.27 |
| Membrane arm tip | 7AQW | 11874 | 1,580 | 0.005 | 0.690 | 0.00 | 89.82 | 0.07 | 11.56 | 2.60 |
| Closed state | 7AR8 | 11876 | 7764 | 0.004 | 0.746 | 0.04 | 90.25 | 0.18 | 15.64 | 2.02 |
| Open state | 7AR7 | 11875 | 7625 | 0.008 | 0.831 | 0.07 | 91.28 | 0.06 | 15.51 | 1.98 |

**Supp. table 7: *Polytomella* sp. complex I model identifiers and quality statistics**

| Description | PDB ID | EMDB ID | Residues built | RMS Bonds Length/Angles | | Ramachandran Outliers (%) | Ramachandran favoured (%) | Rotamer outliers (%) | Clashscore | EMRinger score |
|---|---|---|---|---|---|---|---|---|---|---|
| Full complex | 7ARD | 11880 | 8,731 | 0.005 | 0.730 | 0.05 | 92.49 | 0.04 | 11.93 | 2.83 |
| Peripheral arm | 7ARC | 11879 | 3,408 | 0.007 | 0.724 | 0.06 | 89.75 | 0.07 | 11.13 | 3.34 |
| Membrane arm | 7AR9 | 11877 | 5,323 | 0.005 | 0.702 | 0.11 | 94.26 | 0.02 | 9.17 | 3.36 |
